# Sodium channels expressed in nociceptors contribute distinctly to action potential subthreshold phase, upstroke and shoulder

**DOI:** 10.1101/2023.09.03.556095

**Authors:** Phil Alexander Köster, Enrico Leipold, Jenny Tigerholm, Anna Maxion, Barbara Namer, Thomas Stiehl, Angelika Lampert

**Affiliations:** University Hospital, RWTH Aachen University, Institute for Neurophysiology, Pauwelsstrasse 30, 52074 Aachen, Germany; Scientific Center for Neuropathic pain Aachen SCNAACHEN, Uniklinik RWTH Aachen University, Aachen, Germany; University of Luebeck, Department of Anesthesiology and Intensive Care and CBBM-Center of Brain, Behavior and Metabolism, Ratzeburger Allee 160, 23562 Lübeck, Germany; University Hospital, RWTH Aachen University, Joint Research Center for Computational Biomedicine (JRCC), Pauwelsstrasse 30, 52074 Aachen, Germany; University Hospital, RWTH Aachen University, Interdisciplinary Center for Clinical Research (IZKF), Faculty of Medicine, Research Group Neurosciences, Pauwelsstrasse 30, 52074 Aachen, Germany; University Hospital, RWTH Aachen University, Institute for Computational Biomedicine and Disease modelling with focus on phase transitions between phenotypes, Pauwelsstrasse 30, 52074 Aachen, Germany

## Abstract

Voltage-gated sodium channels (VGSC) in the peripheral nervous system shape action potentials (AP) and thereby support the detection of sensory stimuli. Most of the nine mammalian VGSC subtypes are expressed in nociceptors, but predominantly, three are linked to several human pain syndromes: while Na_v_1.7 is suggested to be a (sub-)threshold channel, Na_v_1.8 is thought to support the fast AP upstroke. Na_v_1.9, as it produces large persistent currents, is attributed a role in determining the resting membrane potential.

We characterized gating of Na_v_1.1-Na_v_1.3 and Na_v_1.5-Na_v_1.9 in manual patch clamp with focus on the AP subthreshold depolarization phase. Na_v_1.9 exhibited the most hyperpolarized activation while its fast inactivation resembled the depolarized inactivation of Na_v_1.8. For some VGSCs (e.g., Na_v_1.1 and Na_v_1.2), a positive correlation between ramp current and window current was detected.

Using a modified Hodgkin-Huxley model which accounts for the time needed for inactivation to occur, we used the acquired data to simulate two nociceptive nerve fiber types (an Aδ-and a mechano-insensitive C-nociceptor) containing VGSC conductances according to published human RNAseq data. Our simulations suggest that Na_v_1.9 is supporting both the AP upstroke and its shoulder. A reduced threshold for AP generation was induced by enhancing Na_v_1.7 conductivity or shifting its activation to more hyperpolarized potentials, as observed in Na_v_1.7-related pain disorders.

Here, we provide a comprehensive, comparative functional characterization of VGSCs relevant in nociception and describe their gating with Hodgkin-Huxley-like models, which can serve as a tool to study their specific contributions to AP shape and sodium channel-related diseases.

**Disclaimer:** Parts of this study were published as a preprint on bioRxiv: Köster, P.A., T. Stiehl, J. Tigerholm, A. Maxion, B. Namer, and A. Lampert. 2023. Biophysics of sodium channels during subthreshold depolarization *in vitro* and *in silico*. bioRxiv. doi.org/10.1101/2023.09.03.556095 (Preprint posted September 6, 2023)

**Summary:** Subthreshold gating of seven sodium channels (Na_v_1.1-3, Na_v_1.5-8) is determined by manual patch clamp and, together with Na_v_1.9, integrated into a computer model of an Aδ-and a mechano-insensitive nociceptor (CMi). Simulations reveal contribution of Na_v_1.9 to the action potential upstroke and shoulder and prove useful for Na_v_1.7-related disease modelling.

## Introduction

Next to their well-known role to initiate the upstroke of the action potential (AP), voltage-gated sodium channels (VGSC) fine-tune cellular excitability by contributing to subthreshold depolarizations and fluctuations of the membrane potential. Thus, they are involved in the detection and amplification of sensory stimuli e.g. in peripheral sensory neurons such as nociceptors (Bhattacharjee et al., 2018; Bennett et al., 2019; Goodwin and McMahon, 2021; Tian et al., 2023). Nociceptive stimuli depolarize the cell membrane of the nerve fiber ending, but if they are small, they do not initiate an AP, leading to subthreshold depolarizations. Subthreshold depolarizations directly preceding an AP are described in rats to be supported by VGSC currents sensitive to the pufferfish neurotoxin tetrodotoxin (TTX) (Enomoto et al., 2006; Vasylyev and Waxman, 2012), but the specific contribution of each channel isoform is yet to be fully understood.

To date, nine isoforms of mammalian VGSC α subunits have been described (Na_v_1.1 – Na_v_1.9), which are encoded by the genes *SCN1A-SCN5A* and *SCN8A-SCN11A* (Yu and Catterall, 2003; Catterall et al., 2005; Dib-Hajj et al., 2010). They differ in their biophysical properties, such as kinetics and voltage-dependence of activation, inactivation, and deactivation (Ahern et al., 2016; Körner and Lampert, 2020; Catterall, 2023) and are classified according to their sensitivity to TTX as either TTX-resistant (TTXr; Na_v_1.5, Na_v_1.8, and Na_v_1.9) or TTX-sensitive (TTXs; all remaining isoforms) (de Lera Ruiz and Kraus, 2015).

Depending on their type and physiological function, excitable human cells express different VGSC isoforms (Catterall et al., 2005; Ahern et al., 2016). For example, Na_v_1.1 – Na_v_1.3 and Na_v_1.6 are widely expressed in neurons of the central (CNS) and peripheral nervous system (PNS) (Whitaker et al., 2000; Vacher et al., 2008; Liang et al., 2021), while Na_v_1.7-Na_v_1.9 are expressed in PNS neurons including dorsal root ganglion (DRG) neurons, sympathetic ganglion neurons or olfactory sensory neurons (Fukuoka et al., 2008; Wang et al., 2017). The skeletal and cardiac muscle express their own specific subtypes: Na_v_1.4 and Na_v_1.5, respectively (Sheets and Hanck, 1999). Na_v_1.5 can also be detected in PNS neurons during development or injury (Renganathan et al., 2002) and at low levels in adults as revealed by recent transcriptomics studies of human sensory neurons (Tavares-Ferreira et al., 2022).

The VGSC expression patterns of different excitable cell populations can overlap and are subject to dynamic changes during development or in response to injury or sensitization. Expression of Na_v_1.3, for example, was described in embryonic, but not in healthy adult rat DRG neurons. However, expression of Na_v_1.3 and associated currents have found to be upregulated in rodent DRG neurons after spinal cord injury (Felts et al., 1997; Hains et al., 2003; Lampert et al., 2006).

The sensation of pain via DRG neurons is associated with the activation of their afferent C-or Aδ-nerve fibers. Among them, the subclass of C-fibers known as “silent”, “sleeping” or mechano-insensitive (CMi) fibers is assumed to be involved in the generation of neuropathic pain by exhibiting abnormal spontaneous activity (Namer and Handwerker, 2009; Kleggetveit et al., 2012). Cell excitability of different fiber types and therefore their function in pain signaling are subject to various factors of influence. Recent transcriptomic studies including human tissue revealed distinct VGSC expression patterns in each sensory nerve fiber type (Usoskin et al., 2015; Kupari et al., 2021; Nguyen et al., 2021; Tavares-Ferreira et al., 2022; Jung et al., 2023; Bhuiyan et al., 2024). To date, the influence of these expression patterns on nerve fiber excitability is yet to be fully understood. It is therefore beneficial to first investigate the contribution of each VGSC isoform to nerve fiber excitability in isolation.

*Gain-of-function* variants of Na_v_1.7 are linked to inherited pain syndromes such as erythromelalgia (Yang et al., 2004; Lampert et al., 2010; Bennett and Woods, 2014; Brouwer et al., 2014; Tang et al., 2015; Dib-Hajj et al., 2017) or paroxysmal extreme pain disorder (Fertleman et al., 2006; Fertleman et al., 2007; Jarecki et al., 2008; Stępień et al., 2020). Studies of these mutations in heterologous expression systems show a prominent shift of voltage-dependence of activation to more hyperpolarized potentials and suggest that Na_v_1.7 plays an important role in the initiation of APs. Recently, however, in recordings of human sensory neurons derived from induced pluripotent stem cells (iPSC), the contribution of Na_v_1.7 to subthreshold depolarizations was observed to be limited, and the channel was considered a threshold channel rather than a subthreshold channel (Meents et al., 2019).

Functional studies in rodents and heterologous expression systems so far suggest that Na_v_1.8 is the main contributor to the upstroke of the AP and determines its duration by shaping the AP’s shoulder (Renganathan et al., 2001; Blair and Bean, 2002; Matsutomi et al., 2006; Han et al., 2015a). Na_v_1.9, on the other hand, is thought to modulate the resting membrane potential by eliciting large persistent currents (Herzog et al., 2001; Maingret et al., 2008; Tian et al., 2023). A contribution of Na_v_1.9 to the AP shoulder was nicely demonstrated in enteric neurons of Na_v_1.9 knockout mice (Osorio et al., 2014) and observed in DRG neurons of mice carrying a Na_v_1.9 missense mutation (Leipold et al., 2013), while another study failed to see an influence of Na_v_1.9 on AP shape (Priest et al., 2005). Thus, the question of the role of Na_v_1.9 in shaping AP properties is left unresolved.

Na_v_1.9 is long known to show poor heterologous expression (Vanoye et al., 2013; Goral et al., 2015; Sizova et al., 2020). Various attempts to express Na_v_1.9 in heterologous expression systems yielded limited success with very few exceptions (Tate et al., 1998; Leipold et al., 2013; Vanoye et al., 2013; Leipold et al., 2015). Although chimeric Na_v_1.9 channels containing the C-terminus from Na_v_1.4 show improved expression, they exhibit altered voltage dependence of channel activation and inactivation (Goral et al., 2015; Bothe and Lampert, 2021) and are therefore unsuitable for studying Na_v_1.9 gating and its contribution to AP genesis.

Once the gating parameters of each channel subtype have been analyzed, their contribution to cellular excitability can be reconstructed and investigated using mathematical and *in silico* models. In 1952, Hodgkin and Huxley provided a mathematical description of VGSC gating and APs, which still today is often used to model electrophysiological data and cellular excitability (Hodgkin and Huxley, 1952; Stiles and Gray, 2021; Wang et al., 2022). However, classic Hodgkin-Huxley models implement VGSC inactivation right at the beginning of a voltage step stimulus and therefore do not account for the duration of protein-level conformational changes that are assumed to occur within the channel for the so-called inactivation particle to bind to its receptor and thereby stopping the associated sodium influx.

C-fiber excitability in particular has been modelled and used successfully to describe the biophysical properties of this fiber type (Petersson et al., 2014; Tigerholm et al., 2014; Tigerholm et al., 2015; Maxion et al., 2023). In most cases, only select VGSC isoforms are included in models of excitable cells, often because of a lack of a standardized comprehensive data set which is needed as basis for model generation. Larger studies comparing VGSC gating *in vitro* in a comparable experimental setting have focused on suprathreshold gating of a subset of VGSCs expressed in the CNS and related mutations leading e.g. to epilepsy (Thompson et al., 2023), or on sensory neuron VGSCs with a focus on their temperature dependence (Kriegeskorte et al., 2023).

As conclusive and complete data on the sub-and suprathreshold activity of all VGSC isoforms are not yet available, we here conducted a standardized and comparative examination of VGSC isoforms involved in nociception (Na_v_1.1-1.3 and Na_v_1.5-1.8) with consistent experimental conditions using patch-clamp experiments in heterologous expression systems (HEK293(T) and ND7/23 cells).

With these data, we developed a computational model which is based on the Hodgkin-Huxley framework and parameterized using our patch-clamp data. We added Na_v_1.9 to this model using previously published data obtained under similar experimental conditions using transfected ND7/23 cells (Leipold et al., 2013; Leipold et al., 2015). We implemented the VGSC isoform proportions as reported for human CMi-fibers and Aδ-fibers (Tavares-Ferreira et al., 2022) to investigate the contributions of individual VGSC isoforms to excitability and AP generation in two sensory fiber types.

## Materials and Methods

### Cell culture and cell preparation

#### Cell lines

Plasmids of VGSC isoforms used in this study were either transfected stably in human embryonic kidney cell line HEK293 (rat Na_v_1.3 = rNa_v_1.3, human Na_v_1.5 = hNa_v_1.5, mouse Na_v_1.6 = mNa_v_1.6, hNa_v_1.7) or transiently in human embryonic kidney cell line HEK293T (hNa_v_1.1, hNa_v_1.2) or mouse/rat hybridoma nerve cell line ND7/23 (hNa_v_1.8).

All used cell lines were cultivated adherently under standard cell culture conditions (i.e., 37°C and 5 % CO2) in their respective cell culture media and supplements (Table S1). Passaging for all cell lines was performed twice per week at approximately 80% confluency using Accutase (Sigma-Aldrich, St. Louis, Missouri, USA) with a 1:5 – 1:10 passaging rate onto cultureware coated with Geltrex (LDEV-Free, hESC-Qualified; Gibco, Thermo Fisher Scientific, Waltham, Massachusetts, USA). Only cells with passage numbers below 30 were used for electrophysiological experiments. All cell lines were regularly tested negative for mycoplasma contamination.

#### Plasmids

Plasmid constructs used for transient transfection were amplified using XL10-gold ultracompetent cells (Agilent Technologies Inc., Santa Clara, California, USA) and verified by restriction pattern analysis and commercial DNA sequencing (Eurofins Genomics GmbH, Ebersberg, Germany). The plasmids for hNa_v_1.1 (pCMV6-AC-hNa_v_1.1 WT RRSSV 3’ UTR, Myc-DDK) and hNa_v_1.2 (pCMV6-XL5-hNa_v_1.2 WT RRSSV 3’ UTR) were kindly gifted by Frank Bosmans, Ghent University, Belgium. hNa_v_1.1 contained the common natural variation A1067T (allele frequency for T: 0.728). The hNa_v_1.8 plasmid (pIRES puro3-hNa_v_1.8 WT) contained the V1073A variant (allele frequency for A: 0.58).

#### Transient transfection

HEK293T and ND7/23 cells were seeded in 35 mm petri dishes 20-24 hours prior to transient transfection to reach a 70-90% confluency. Transfection was then performed using 3 µl jetPEI (Polyplus-transfection, Illkirch, France), 0.25 µg reporter plasmid pMax-GFP (Lonza, Basel, Switzerland) and 1.25 µg plasmid encoding the VGSC to be examined. Cells were incubated for another 18-24 hours before being split onto fresh 35 mm petri dishes. About 2 hours later, after the seeded cells attached to the cultureware, cells with a bright green fluorescence were used for patch clamp experiments.

### Electrophysiology

#### General patch clamp framework

Electrophysiological experiments were performed in whole-cell voltage clamp mode using an EPC10-USB amplifier (HEKA Elektronik GmbH, Lambrecht/Pfalz, Germany). Cells were covered in extracellular solution (ECS) containing 140 mM NaCl, 3 mM KCl, 1 mM MgCl_2_, 1 mM CaCl_2_, 10 mM HEPES and 20 mM Glucose. ECS was adjusted to pH 7.4 using NaOH, while osmolarity was between 305 and 312 mosm/l. Since ND7/23 cells display endogenous VGSC-mediated TTXs currents (Rogers et al., 2016; Lee et al., 2019), 500 nM TTX (Tocris, Bristol, United Kingdom) was added to the ECS for measurements of the TTXr channel hNa_v_1.8.

Glass pipettes from borosilicate glass tubes (Biomedical Instruments, Zöllnitz, Germany) were manufactured and fire-polished using a DMZ pipette puller (Zeitz Instruments GmbH, Martinsried, Germany). Pipette tip resistance (R_pip_) was between 0.8 and 2 MΩ for all measurements. The pipette was filled with intracellular solution (ICS) containing 10 mM NaCl, 140 mM cesium fluoride (CsF), 1 mM EGTA, 10 mM HEPES and 18 mM Sucrose. ICS was adjusted to pH 7.31 with CsOH, while osmolarity was 313 mosm/l.

Experiments were conducted at room temperature (22 ± 1°C). Cells were kept in ECS for a maximum of 90 minutes before being discarded. Cell membrane capacitance was between 5 and 33 pF. For all measurements, seal resistance (R_seal_) was above 0.8 GΩ. Only cells with an initial series resistance (R_s_) below 5 MΩ and a peak current amplitude above 500 pA during the voltage dependence of activation protocol were accepted. At all times, voltage error (R_s_ * I) was below 5 mV.

Data were gathered using Patchmaster Next version 1.2 (HEKA Elektronik GmbH, Lambrecht/Pfalz, Germany). Analog signals were digitized at a sampling rate of 100 kHz (except for AP clamping measurements, which were sampled at 20 kHz) with a 10 kHz Bessel low-pass filter. Leak currents were measured and subtracted online (for activation, steady-state fast inactivation (SSFI) and ramp current measurements) or offline (for AP clamping due to technical limitations) with a *P/n* procedure (*n* = 4) preceding the respective test pulse. Capacitive currents were cancelled, and R_s_ was compensated for by 77-83% for all cells. All measurements were corrected online for a liquid junction potential of 8.2 mV.

#### Patch clamp protocols

For each cell, stimulation protocols were only recorded once (no technical replicates). Protocols are indicated in the respective figures. For all protocols, the holding potential in between test pulses (V_hold_) was set to -120 mV.

After establishing whole-cell configuration, a current stabilization protocol was conducted for 5 minutes to ensure complete recovery, in which cells were held at holding potential and repeatedly stimulated with test pulses to 0 mV every 10 seconds.

To investigate the voltage dependence of activation, a series of 40 ms pulses from holding potential to a test pulse potential was elicited, with the test pulse potential increasing stepwise from -90 mV to +40 mV in 10 mV increments (-60 mV to +70 mV for most Na_v_1.8 measurements). The interpulse interval was 5 s. For assessment of the voltage dependence of activation (i.e., the fraction of channels activated to an open state depending on the membrane potential), we calculated the voltage dependent sodium conductance (*G_Na_*) for each test pulse, using the equation

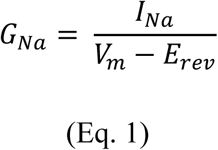

with *I_Na_* being the peak inward current at the respective test pulse voltage *V_m_*, and *E_rev_* being the sodium reversal potential individually determined for each cell. To exclude effects of varying cell size and current density, *G_Na_* values were normalized to the maximum conductance *G_Na, max_* of that cell, before being plotted against test pulse voltages.

During the voltage dependence of SSFI protocol, after being held at holding potential, cells were initially stimulated with a 500 ms prepulse, before being exposed to a 40 ms test pulse at 0 mV. From sweep to sweep, the prepulse voltage increased stepwise from -140 mV to -30 mV in 10 mV increments (-110 mV to 0 mV for most Na_v_1.8 measurements). The interpulse interval was 10 s. For investigation of the voltage dependence of SSFI (i.e., the fraction of non-inactivated channels available for activation depending on the preceding membrane potential), the peak inward current for each test pulse (*I_Na_*) was normalized to the maximum inward current during inactivation measurement (*I_Na,max_*). Normalized current values were plotted against the respective prepulse potential.

Voltage dependence of activation conductance-voltage curves and voltage dependence of SSFI current-voltage curves were fitted with non-linear regression using the Boltzmann equation

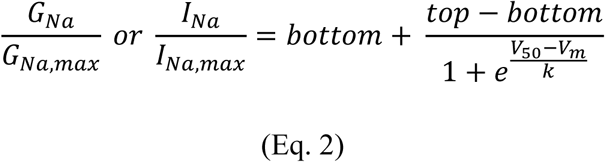

with *V_50_* being the membrane potential at half-maximal channel (in)activation in mV, *V_m_* being the membrane voltage in mV and *k* being the slope factor in 1/mV. The top value of the sigmoid curve was constrained to 1 for both activation and inactivation measurements. The bottom value was constrained to 0 for activation measurements but was not constrained for inactivation measurements to examine the fraction of channels remaining open after SSFI. The goodness of fit was *R²* > 0.9 for all Boltzmann fits.

We equated the window current for each cell as the area under the curve (AUC) of the superimposed activation and SSFI Boltzmann fit curves. Because SSFI Boltzmann fits do not reach 0 values when approaching infinity voltages, the AUC was calculated with an upper limit set 20 mV above the intersection of activation and SSFI Boltzmann fits for each cell.

For ramp current measurements, cells were stimulated with depolarizing voltage ramps from holding potential to +20 mV at different ramp rates, namely 0.1, 0.2, 0.4, 0.6, 1.2, 2, 4, and 6 mV/ms (for ramp durations see Table S2). For each ramp rate, the measured current response was averaged from three consecutive sweeps. The maximum inward current and the voltage at which it occurs were measured, as well as the area under the curve (AUC) of the current response. Because especially for Na_v_1.8, the ramp current bell curve did not return to 0 pA at the end of the ramp, the AUC was calculated only up to the maximum inward current. Maximum inward current and AUC were normalized to the respective maximum inward current from the activation protocol for each cell (*I_Na,max_*).

To measure the VGSC current response to APs, three different pre-recorded APs (AP1 – AP3) measured in iPSC-derived nociceptors were used as a voltage command (Blair and Bean, 2002). The AP commands differ in their characteristics (Table 1) with e.g., subthreshold voltage slope being between 0.2 and 2 mV/ms. These voltage stimuli included 50 ms of the respective resting membrane potential preceding each recorded AP (Table 1) to prevent transient currents caused by step depolarization from holding potential to initial AP voltage from interfering with the examined AP current responses. The respective AP current response was averaged out of three consecutive sweeps. For each stimulation, we calculated the maximum inward current and AUC of both the total AP and of the subthreshold phase of the AP. For the latter, the slope of the elicited current was calculated as well. Current responses for ramp and AP clamping were normalized to the maximum inward current during activation measurement (*I_Na,max_*).

**Table 1:**
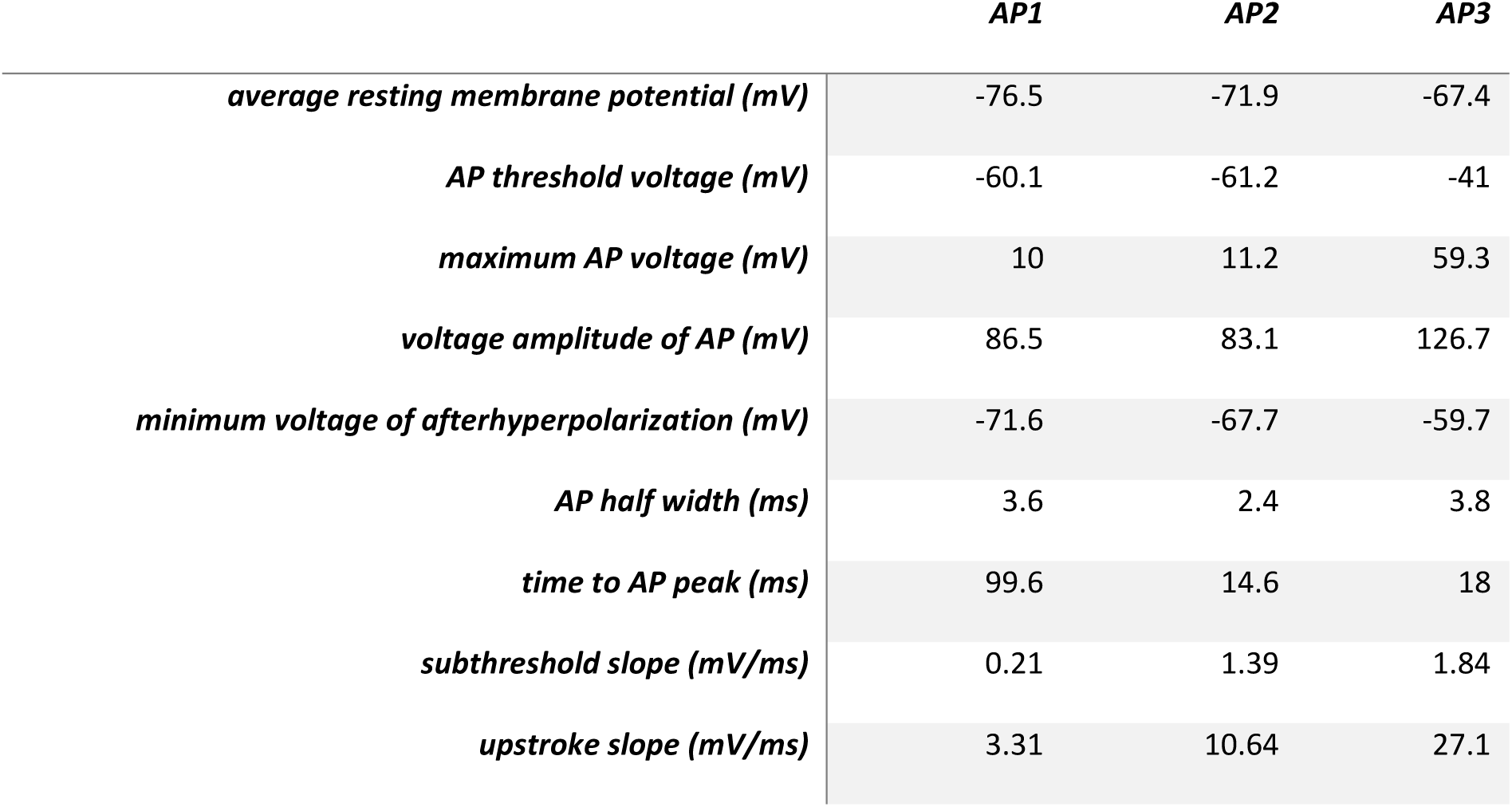
Properties of the Pre-Recorded Action Potentials Used as Voltage Commands. . Data were gathered from current clamp measurements of iPSC-derived nociceptors using the Igor Pro software.

### Sample sizes and data exclusion

We performed an a-priori analysis to estimate the number of cells needed for each experiment. In a small pre-study with Na_v_1.3 and Na_v_1.7, the voltage at which the maximum inward ramp current appeared was compared, assuming normal distribution. Using the software G*Power version 3.1.9.7 (Heinrich-Heine-Universität, Düsseldorf, Germany) we calculated the sample size for each VGSC to be *n* = 27 (*hedge’s g\** = 1.23 as a measure of the effect size *d*, two-tailed *α* = 0.05 and power (1-*β*) = 0.95).

We designed the study to have groups of equal size. Group size variation might occur due to cells being excluded from analysis a-posteriori, e.g., because of measurement artifacts impeding data analysis. Since the ramp current responses often showed a considerable drift, which made it difficult to calculate the AUC, the following exclusion criteria were defined for ramp current AUC measurements: cells with a drift larger than +100 pA or -500 pA as well as cells with a drift larger than 10% of the maximum inward current during activation measurement were excluded from analysis. For AP clamping measurements, we identified outliers using the ROUT method with *Q* = 0.1% (Motulsky and Brown, 2006). Outliers were excluded from further analysis. Beyond that, we performed no outlier analysis or elimination.

### Data analysis

Data analysis for electrophysiological experiments was blinded except for Na_v_1.5. Data gathered by electrophysiological experiments were analysed using FitMaster version 2x90.5 (HEKA Electronik GmbH, Lambrecht, Germany) and IgorPro version 6.37 (WaveMetrics Inc., Portland, Oregon, USA). IgorPro analysis procedures are available upon reasonable request. GraphPad Prism version 9.5.1 (GraphPad Software, San Diego, California, USA) was used for Boltzmann fits and statistical testing. Graphing of data was perfomed with IgorPro, GraphPad Prism and CorelDRAW 2017 version 19.0.0.328 (Alludo, Ottawa, Ontario, Canada).

### Statistics

Data are presented as mean ± standard deviation (SD) unless stated otherwise. Error bars denote the SD, while box plot whiskers show the 10^th^ and 90^th^ percentile. Threshold for statistical significance was set to *p* < 0.05. Given *p* values are shown with 4 decimal places and summarized as follows: ns = not significant, * = *p* < 0.05, ** = *p* < 0.01, *** = *p* < 0.001, ****= *p* < 0.0001.

Data were tested for normal distribution using the D’Agostino & Pearson omnibus K2 test. Normal distribution for an experiment was only assumed if each group passed the test individually. For statistical testing, groups were compared by ordinary one-way ANOVA with Tukey-Kramer’s multiple comparisons test and a single pooled variance for parametric testing or Kruskal-Wallis test with Dunn’s multiple comparison test for non-parametric testing. Values derived from ramp current measurements (*I_max_*, *U_location of Imax_* and AUC) were compared by ordinary two-way ANOVA with Tukey-Kramer’s multiple comparisons test and individual variances computed for each comparison, to compare between both VGSC isoforms and an additional factor of influence, i.e., the ramp rate. For correlations, we computed the nonparametric Spearman correlation with a two-tailed *p* value (data are given as Spearman’s *r* with 95% CI) and graphed it with simple linear regression. Data were neither matched nor paired.

The exact group *n* values are not shown in the figures for readability purposes but are indicated in the respective tables and supplementary tables.

### Computational modelling

To simulate the AP of a CMi-and an Aδ-fiber we use a modified version of the Hodgkin-Huxley model. Compared to the original Hodgkin-Huxley model (Hodgkin and Huxley, 1952) we introduced a delay for current decay (fast inactivation kinetics). This modification prevents premature inactivation of sodium channels and improves the fit of the model to sodium currents of Na_v_1.1 -Na_v_1.3 and Na_v_1.5 - Na_v_1.8 for activation voltages above 10 mV. The gating variables and the delay of inactivation were fitted using voltage clamp data for Na_v_1.1 - Na_v_1.3 and Na_v_1.5 - Na_v_1.8 acquired at room temperature (22 ± 1°C) as well as previously published voltage clamp data for Na_v_1.9 (Leipold et al., 2013; Leipold et al., 2015). To fit Na_v_1.9 kinetics, the original Hodgkin-Huxley equations were used because they accurately captured the data.

The fits were performed with *fmincon* from the Matlab software R2021b (The MathWorks, Inc., Natick, Massachusetts, USA) using a multistart approach with latin hypercube sampling and a weighted least squares cost functional. To simulate APs in a CMi-fiber and an Aδ-fiber the ratios between the maximal conductances for the considered VGSC isoforms were chosen to match recently published putative gene expressions quantified in human DRGs by spatial transcriptomics (Tavares-Ferreira et al., 2022). The simulations are based on the quantifications of gating variables at room temperature. The resting potential is -71 mV, and the model is at rest before different stimulation currents are applied. Details, model equations and parameters are provided in the supplementary information.

## Results

### Na_v_1.5 and Na_v_1.8 activate and fast inactivate at more hyper-and depolarized potentials than other subtypes, respectively

To gain a comparable oversight of the biophysics of VGSC isoforms involved in nociception, we examined Na_v_1.1, Na_v_1.2, Na_v_1.3, Na_v_1.5, Na_v_1.6, Na_v_1.7 and Na_v_1.8 for their voltage dependence of activation and steady-state fast inactivation (SSFI, Fig. 1 and 2). Of all channels examined two stood out: Na_v_1.5 activated and fast inactivated at more hyperpolarized potentials than all other VGSC isoforms (Fig. 1G and 2A/B, Table 2 and S3), whereas Na_v_1.8 was set apart by its more depolarized activation and SSFI (Fig. 1J and 2A/B, Table 2 and S3). The TTXs VGSCs examined activated at intermediate voltages with Na_v_1.6 and Na_v_1.7 being grouped at slightly more hyperpolarized voltages than Na_v_1.1, Na_v_1.2 and Na_v_1.3 (Fig. 1 and 2A/B, Table 2 and S3). Data obtained from fitting conductance-voltage or current-voltage curves to the Boltzmann equation are summarized in Fig. 2, Table 2, and Table S3-S6. Comparison of the Boltzmann equation fitting results showed differences between the VGSC isoforms in all obtained parameters (*p* < 0.0001 for all ANOVA or Kruskal-Wallis tests).

**Figure 1:**
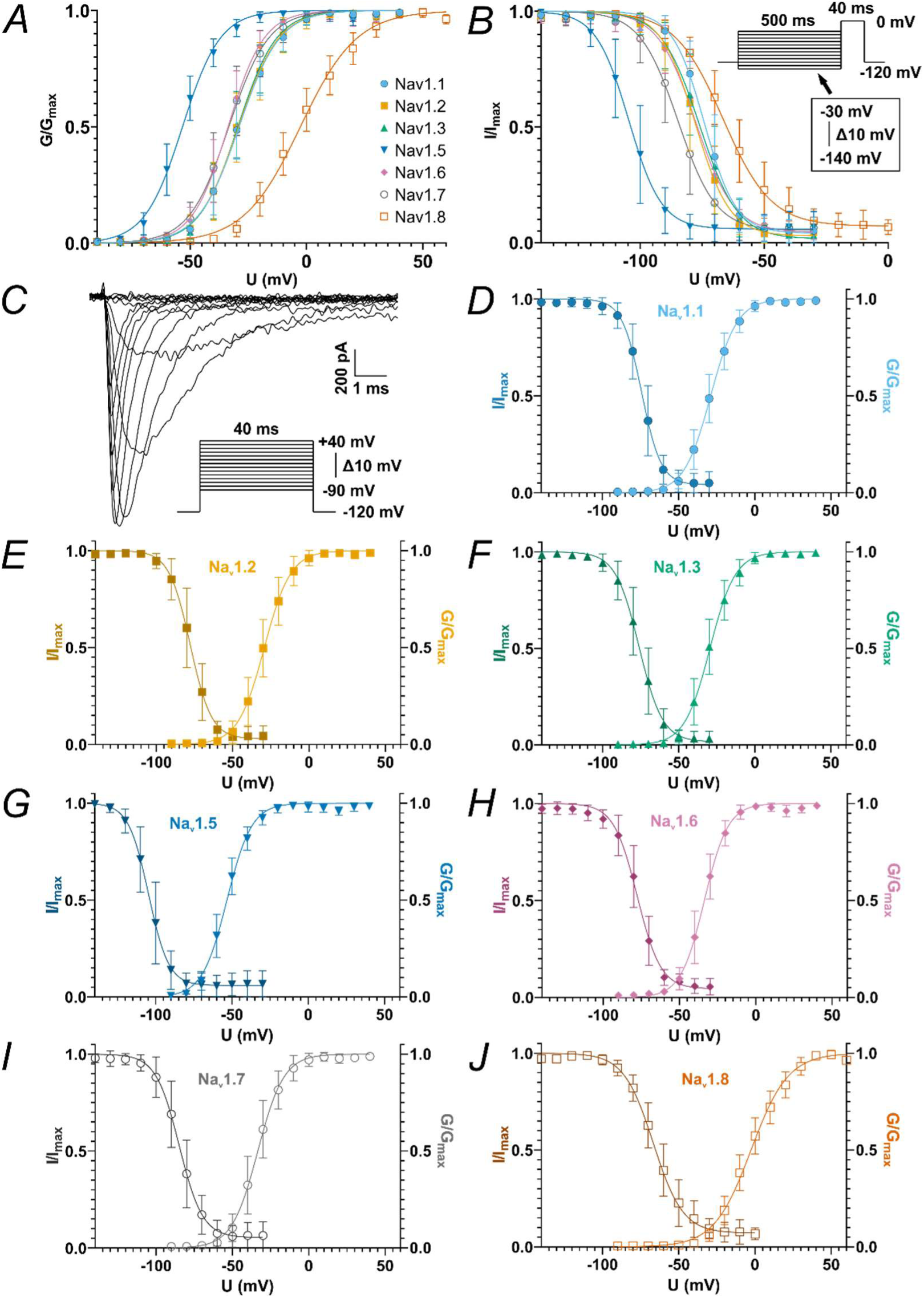
Conductance-Voltage or Current-Voltage Relationship of Activation and SSFI Protocols. *A* overview of all conductance-voltage curves derived from activation measurements. *B* overview of all current-voltage curves derived from SSFI measurements (voltage protocol of SSFI depicted in inlet. Note that for Na_v_1.8 measurements, most cells were stimulated with a -110 mV to 0 mV voltage range). *C* example current traces of Na_v_1.7 elicited by the activation protocol (voltage protocol depicted in inlet. Note that for Na_v_1.8 measurements, most cells were stimulated with a -60 mV to +70 mV voltage range). *D-J* overlay of activation and inactivation curves for Na_v_1.1 (*D*), Na_v_1.2 (*E*), Na_v_1.3 (*F*), Na_v_1.5 (*G*), Na_v_1.6 (*H*), Na_v_1.7 (*I*) and Na_v_1.8 (*J*). Data are shown as mean ± SD and superimposed Boltzmann fit curve.

**Figure 2:**
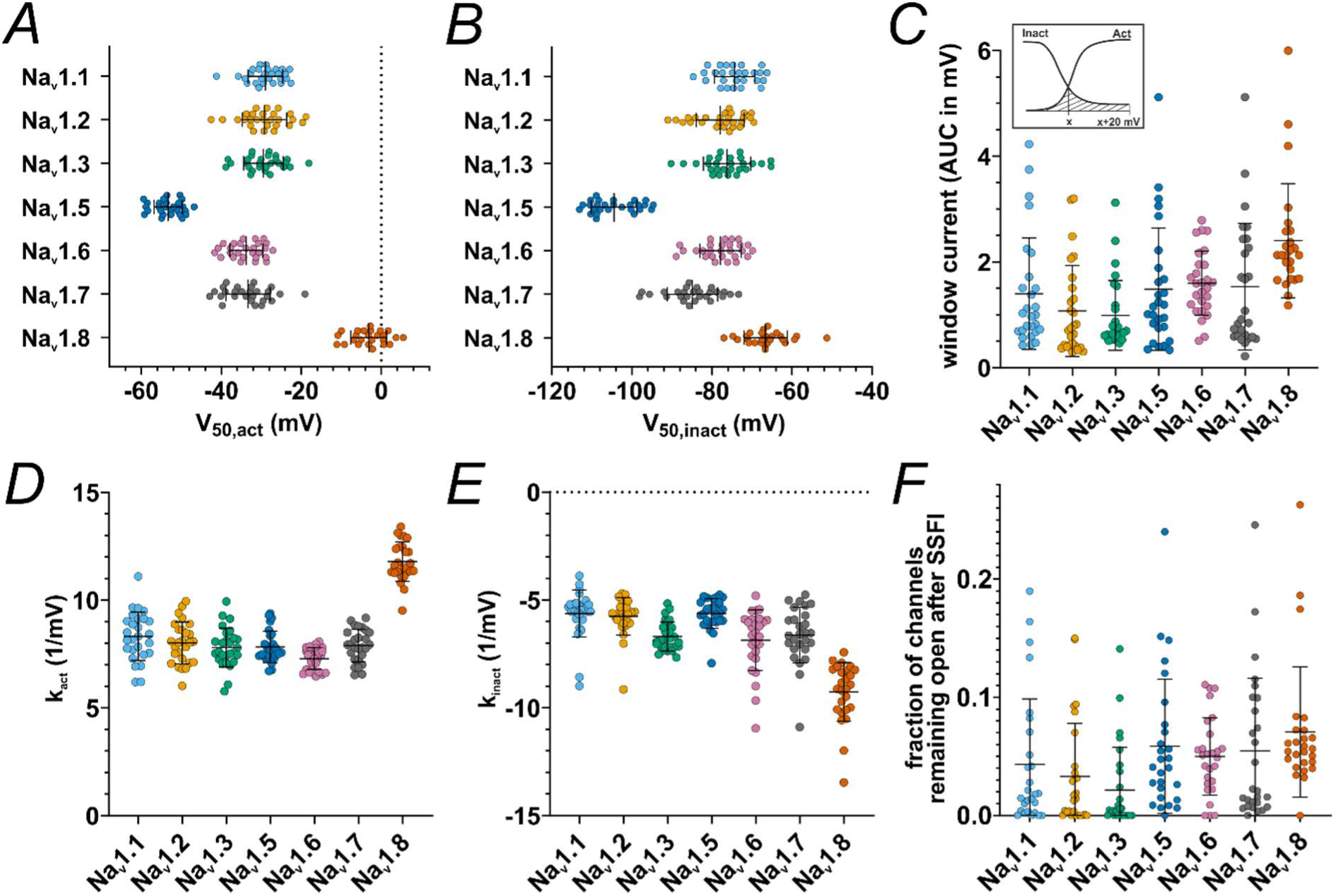
Activation and SSFI Parameters and Window Current. *A V_50_* values obtained from activation Boltzmann fits. *B V_50_* values obtained from SSFI Boltzmann fits. *C* AUC values underneath the superimposed activation and SSFI Boltzmann fit curves, i.e., the window current. *Inlet:* schematic depiction of AUC calculation, *x* being the membrane voltage of the activation and SSFI Boltzmann fit curve intersection. *D* slope values *k* obtained from activation Boltzmann fits. *E* slope values *k* gathered from SSFI Boltzmann fits. *F* bottom values gathered from SSFI Boltzmann fits, indicating the fraction of channels remaining open after complete SSFI. Data are shown as mean ± SD. Statistical significance from multiple comparisons has not been indicated in the figure panels for readability purposes but can be consulted in Table S3-S6.

**Table 2:**
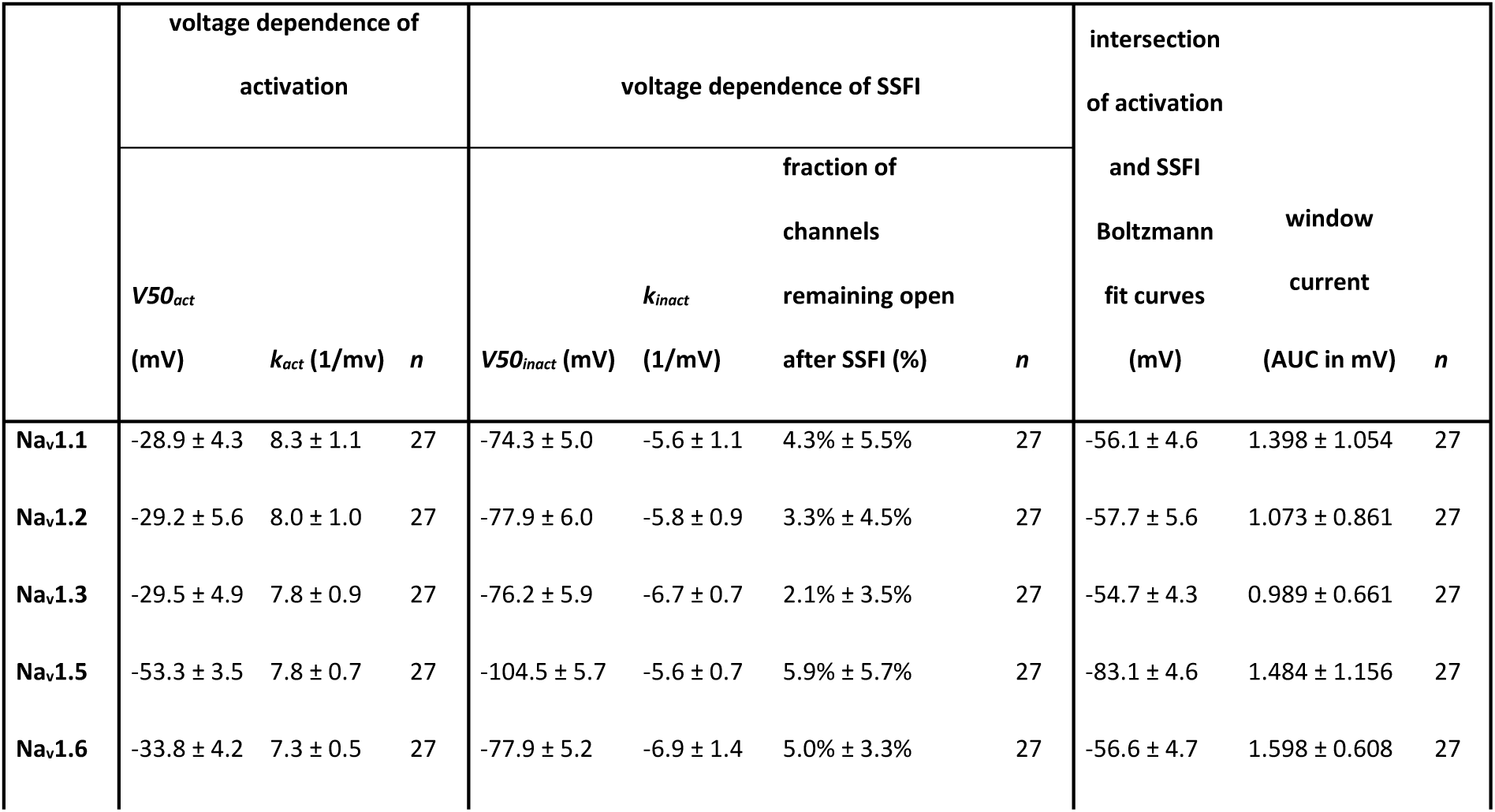
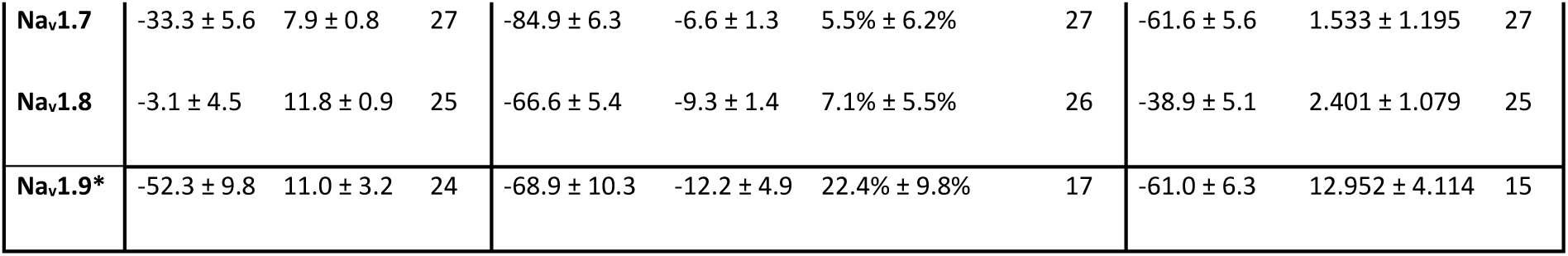
Descriptive Statistics of Parameters Obtained From Fitting Voltage Dependence of Activation and SSFI With the Boltzmann Equation. Data are shown as mean ± SD. *Na_v_1.9 Boltzmann values were derived from pooled previously published data from Leipold et al. 2013 and Leipold et al. 2015.

The overlap between activation and SSFI curves is often referred to as window current (Wang et al., 1996; Magistretti and Alonso, 1999; Abriel et al., 2001; Osteen et al., 2017). The intersection of activation and SSFI curves (i.e., the peak of the window current) of Na_v_1.7 was approximately 5 mV more hyperpolarized than that for all other TTXs channels tested except for Na_v_1.2 (Fig. 1, Table 2, Table S5). This indicates that in this heterologous expression system a higher fraction of Na_v_1.7 channels are in an open state (i.e., activated and not yet fast inactivated) at more hyperpolarized voltages than the other assumed subthreshold channels Na_v_1.1, Na_v_1.3 and Na_v_1.6, and thus could contribute to cell depolarization at subthreshold voltages.

Window current size was larger for Na_v_1.8 than for every other VGSC examined except for Na_v_1.6. Among TTXs channel isoforms, Na_v_1.6 elicited slightly higher window currents than Na_v_1.2 and Na_v_1.3 but did not differ from Na_v_1.1 and Na_v_1.7 (Fig. 2C, Table 2, Table S5), underlining their equal potential to support subthreshold depolarization. Concerning the fraction of channels remaining open after SSFI, we observed a trend for Na_v_1.2/3 and Na_v_1.8 to smaller or larger currents, respectively (Fig. 2F, Table 2, Table S6).

To complete our data set with the last remaining VGSC essential for nociceptors, we analyzed pooled Na_v_1.9 data from previously published recordings (Leipold et al., 2013; Leipold et al., 2015) (Fig. 3, Table 2). While activation of Na_v_1.9 occurred at hyperpolarized voltages comparable to Na_v_1.5, SSFI voltage dependence resembled the depolarized inactivation of Na_v_1.8 (Fig. 3B/C, Table 2). Na_v_1.9 displayed pronounced persistent currents (Fig. 3A) as described previously (Herzog et al., 2001; Maingret et al., 2008). These properties enable the formation of major window currents exceeding those of all other VGSC isoforms measured in this study (Fig. 3E, Table 2).

**Figure 3:**
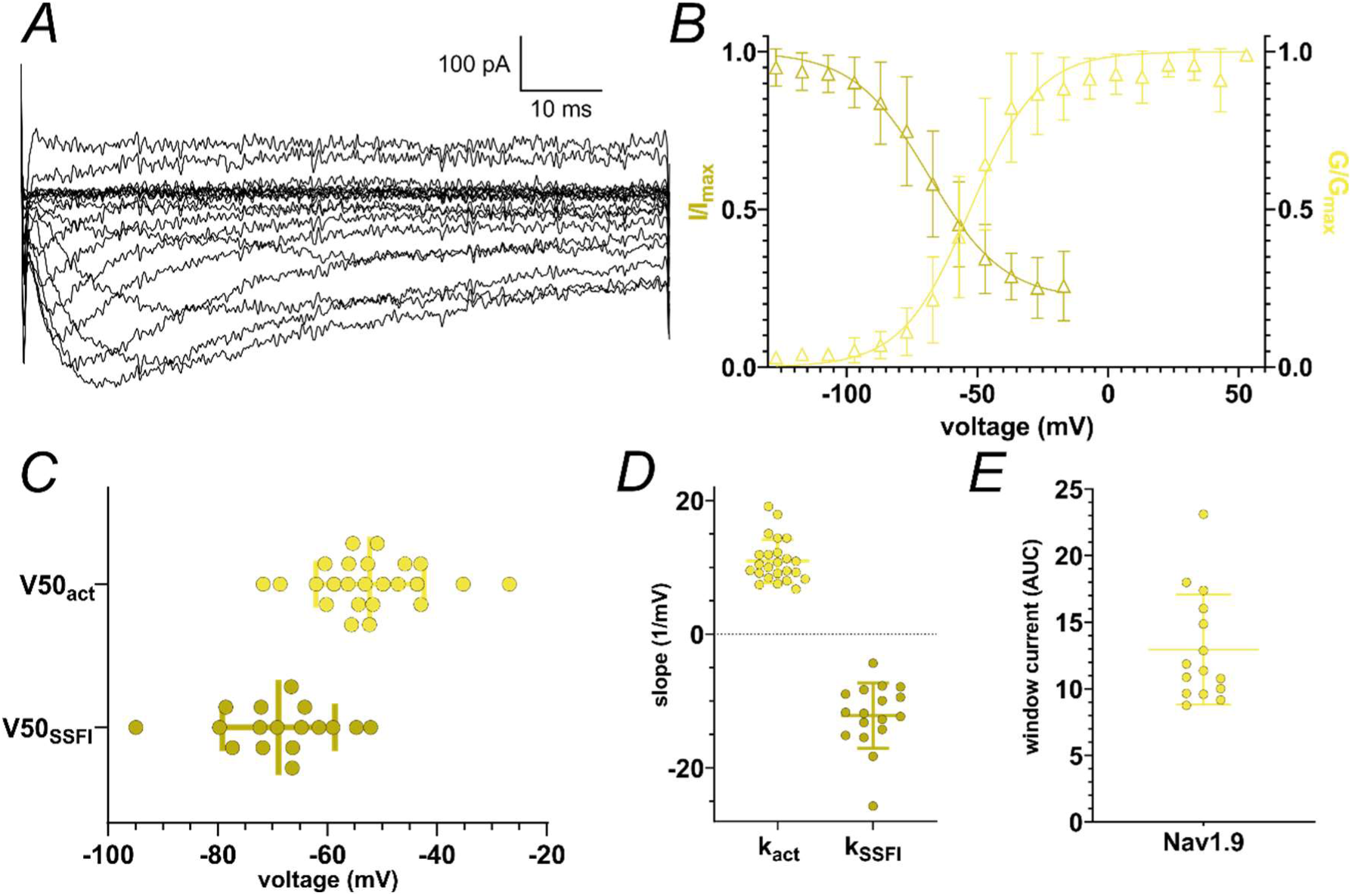
Summary of Nav1.9 Activation and SSFI Data. Data from Leipold et al. 2013 and Leipold et al. 2015 were pooled and analyzed concerning Na_v_1.9 gating and window currents. *A* example current traces of Na_v_1.9 elicited by the activation protocol. Note that cells were stimulated with a -127 mV to +63 mV voltage range. *B* overlay of activation conductance-voltage and SSFI current-voltage curves. *C V_50_* values obtained from activation and SSFI Boltzmann fits. *D* slope values *k* obtained from activation and SSFI Boltzmann fits. *E* AUC values underneath the superimposed activation and SSFI Boltzmann fit curves, i.e., the window current. Data are shown as mean ± SD.

### Ramp and window currents of only some VGSC isoforms may arise from a similar mechanism

Ramp stimuli evoked a bell-shaped inward current response for all tested VGSC isoforms (Fig. 4), as reported previously (Cummins et al., 1998; Abriel et al., 2001; Herzog et al., 2003; Power et al., 2012; Estacion and Waxman, 2013; DeCaen et al., 2014; Han et al., 2015a; Zhang et al., 2017). For cells expressing Na_v_1.3, we observed two peaked ramp currents for slowly depolarizing ramps (i.e., 0.1 mV/ms and partly 0.2 mV/ms) in six out of a total of 27 recordings, similar to what was reported previously (Estacion and Waxman, 2013) (see Fig. S1). For the remaining cells and higher ramp rates, a delayed current decay was observed, potentially corresponding to Na_v_1.3 persistent current.

**Figure 4:**
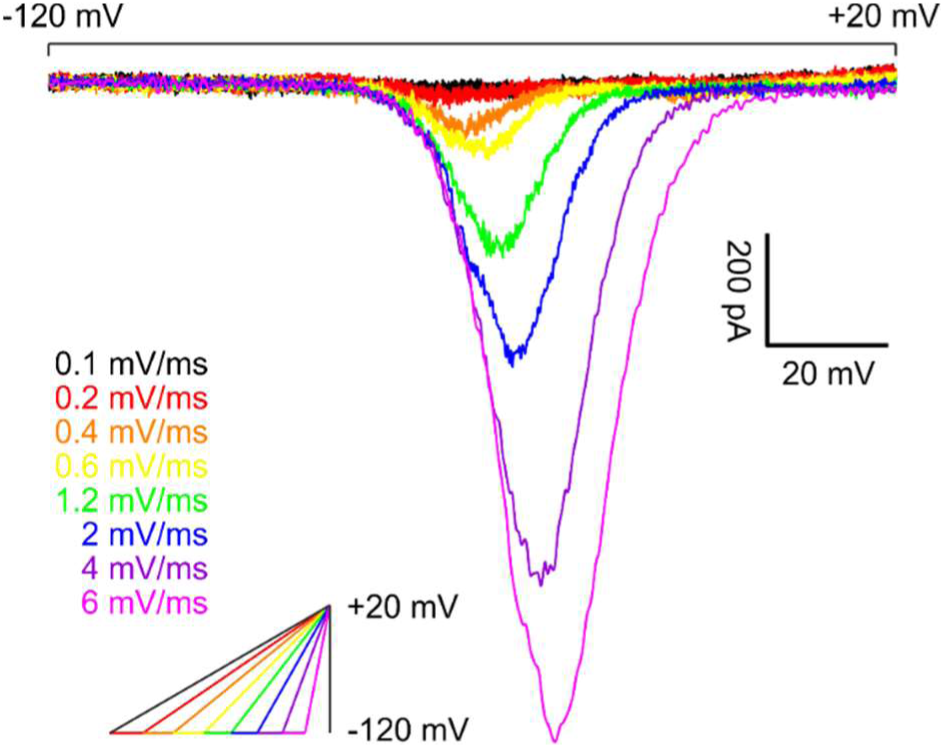
Example Traces of Na_v_1.7 Ramp Currents Elicited by Slowly Depolarizing Ramp Stimuli. Voltage protocol depicted in inlet. Scale bars apply to current traces only.

Ramp currents for all tested isoforms increased with steeper ramp stimuli, most likely due to more channels activating before the onset of fast inactivation. As determined in post-hoc testing, for the lowest ramp rates (i.e., ≤ 0.4 mV/ms), most VGSC isoforms did not differ in maximum inward ramp current; only Na_v_1.8 elicited higher maximum ramp currents than the other VGSC isoforms (Table S7). For medium ramp rates (0.6 – 1.2 mV/ms), Na_v_1.3 and Na_v_1.8 exceed the other VGSC isoforms, whereas for the highest ramp rates (2 – 6 mV/ms), Na_v_1.6 and Na_v_1.7 gradually catch up, leaving Na_v_1.1, Na_v_1.2, and Na_v_1.5 grouped at lower values (Fig. 5A, Table S7/S8).

**Figure 5:**
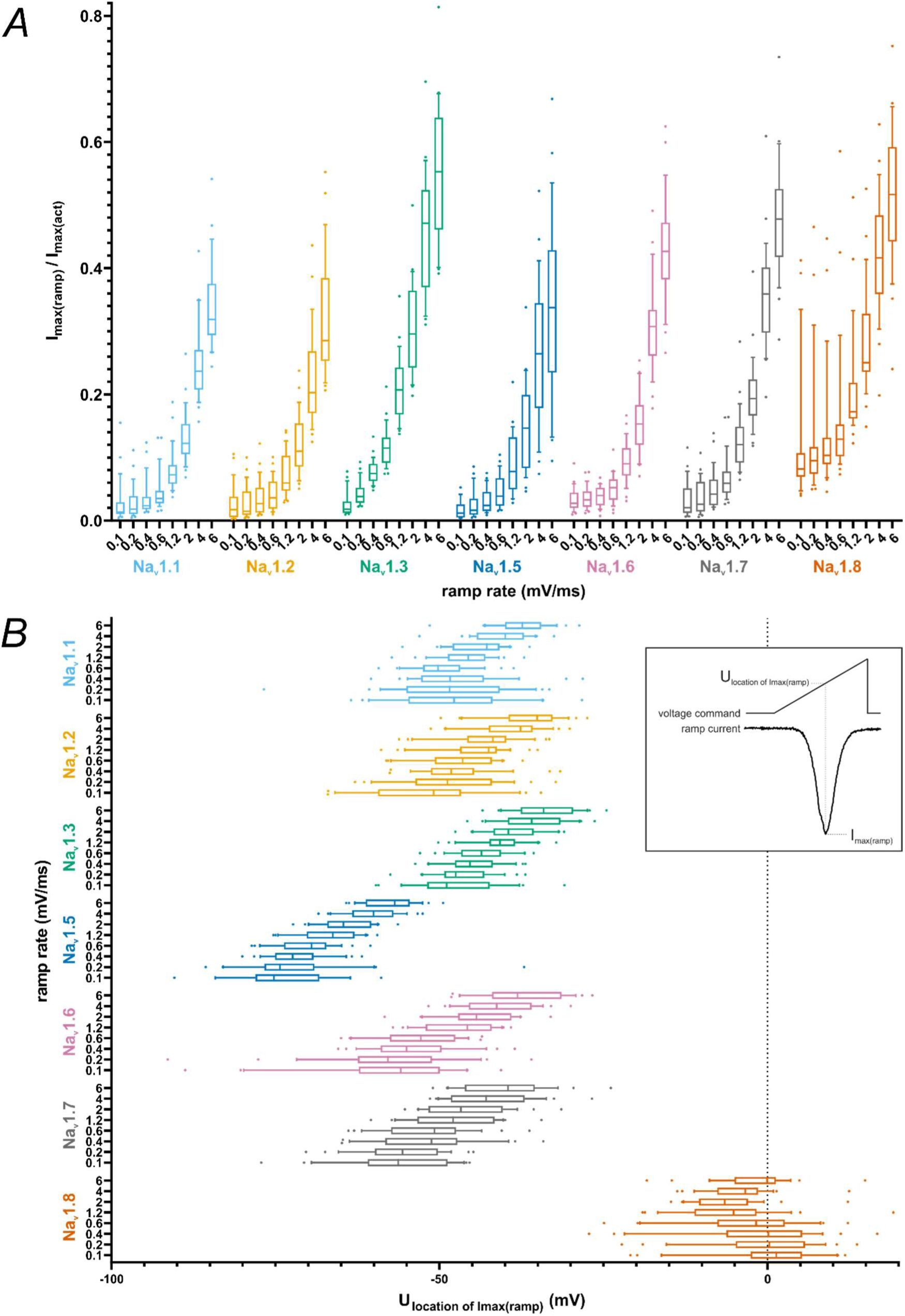
Current-Voltage Parameters Obtained From Ramp Current Measurements. *A* maximum inward current from ramp current measurement normalized to maximum inward current from activation measurements. *B* voltage at which the maximum inward current occurred during ramp current measurement. Data are shown as box plots with whiskers indicating the 10^th^ and 90^th^ percentile and dots for measurements below the 10^th^ and above the 90^th^ percentile.

We observed that the maximum inward current evoked by ramp stimulation was affected more by the ramp rate than by the VGSC isoform: the difference between VGSC isoforms (*F*(6, 1429) = 153.0, *p* < 0.0001) and ramp rate (*F*(7, 1429) = 974.8, *p* < 0.0001) made up for 9.7% and 71.8% of total variation, respectively (Fig. 5A, Table S7). Interaction between these terms accounted for 3.3% of total variation (*F*(42, 1429) = 7.528, *p* < 0.0001).

Ancillary to the size of the maximum inward ramp current, we saw differences in its voltage dependence (Fig. 5B, Table S7) between VGSC subtypes (*F*(6, 1429) = 1748, *p* < 0.0001; 82.5% of total variation), ramp rates (*F*(7, 1429) = 90.15, *p* < 0.0001; 5% of total variation) and their interaction (*F*(42, 1429) = 5.17, *p* < 0.0001; 1.7% of total variation). Using multiple comparison testing, VGSC isoforms showed a distribution matching the results of activation/SSFI *V_50_* results and Boltzmann fit intersections for the vast majority, with Na_v_1.5 and Na_v_1.8 ramp currents occurring at more hyperpolarized and depolarized voltages, respectively, than all other TTXs ramp currents tested (Table S7/S9). TTXs channels are grouped at intermediate values, with Na_v_1.6 and Na_v_1.7 shifted towards more hyperpolarized values than Na_v_1.1, Na_v_1.2, and Na_v_1.3. For steeper ramps, this effect intensified, especially for Na_v_1.7 and Na_v_1.3.

As a measure of charge added to a cell during slow depolarizations such as the subthreshold phase, we investigated the AUC underneath ramp currents (Fig. 6A, Table S7). Post-hoc testing evidenced that Na_v_1.8 had larger AUC values than the other VGSC isoforms for all ramp rates (Table S10). No distinct differences in the AUC between the remaining channel isoforms were observed for lower ramp rates (≤ 0.6 mV/ms). With rising ramp rates, the AUC of Na_v_1.3, Na_v_1.5, Na_v_1.6, and Na_v_1.7 gradually surpasses the AUC of Na_v_1.1 and Na_v_1.2. The AUC is affected by VGSC isoform (*F*(6, 1180) = 95.75, *p* < 0.0001; 15% of total variation), ramp rate (*F*(7, 1180) = 281.6, *p* < 0.0001; 51.4% of total variation) and their interaction (*F*(42, 1180) = 1.984, *p* = 0.0002; 2.2% of total variation).

**Figure 6:**
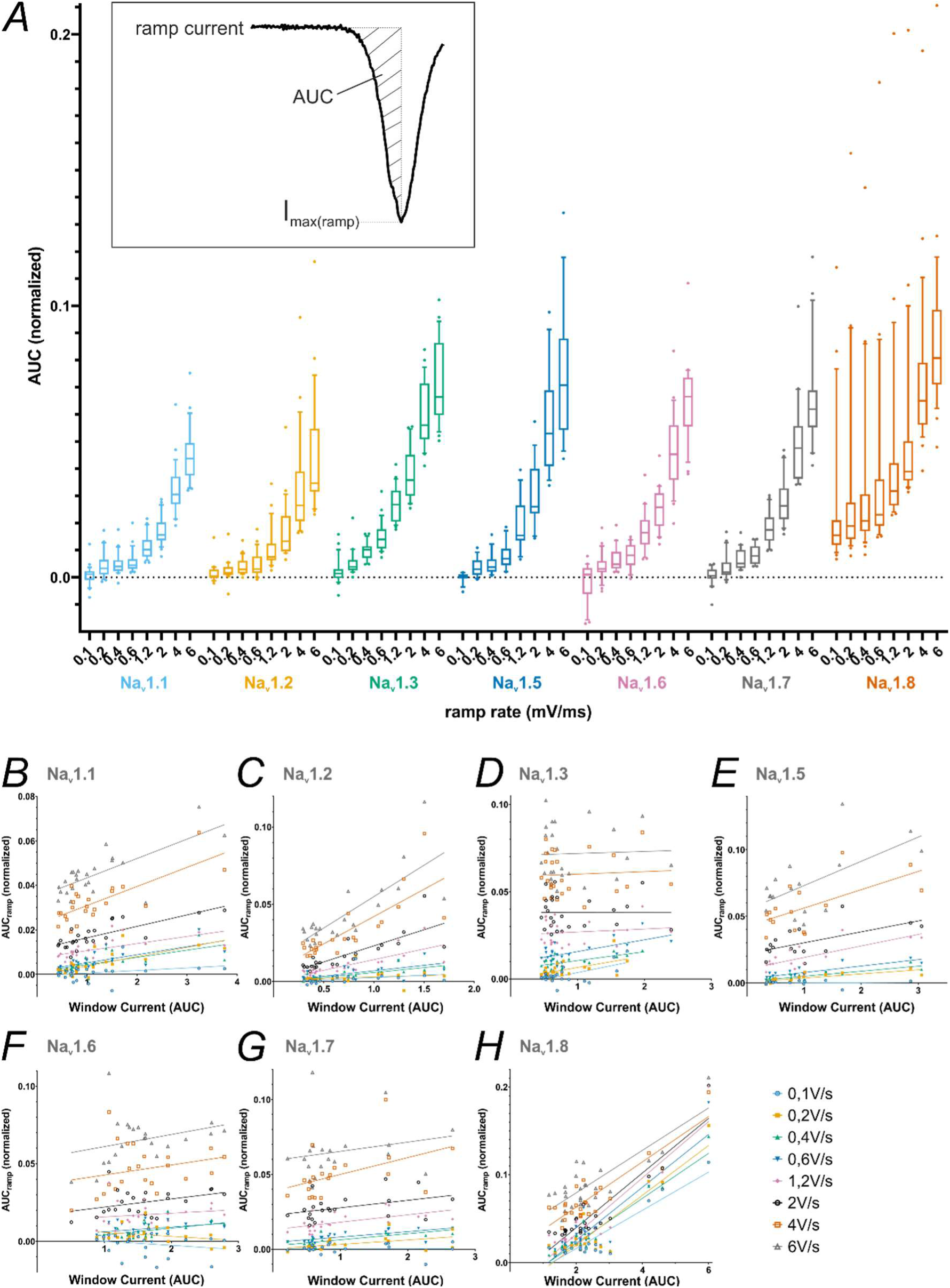
AUC Values From Ramp Current Measurements Correlate With Window Current Measurements for Some VGSC Isoforms. *A* AUC values obtained from ramp current measurements, normalized to the maximum inward current from activation measurements for each cell (inlet depicts a schematic example of an AUC underneath a ramp current curve. Note that the AUC was calculated only up to the point of maximum inward current to make up for Na_v_1.8 ramp currents being cut off at +20 mV). Data are shown as box plots with whiskers indicating the 10^th^ and 90^th^ percentile. *B-H* correlation between AUC values from ramp current measurements and window current values from activation/SSFI measurements. Data are shown as AUC values plotted against the window current values of the respective cell with a linear regression line for each ramp rate

Some authors refer to window current as corresponding to persistent or ramp current (Blair and Bean, 2002; Enomoto et al., 2006; Matsutomi et al., 2006; Vasylyev and Waxman, 2012; Estacion and Waxman, 2013), although so far we are unaware of a study investigating this potential link. We used Spearman’s rank correlation to test for a relationship between window current and ramp current AUC (Fig. 6B-H, Table S11). For Na_v_1.1 and Na_v_1.2, a positive correlation was identified for all ramp rates. While Na_v_1.5, Na_v_1.7 and Na_v_1.8 exhibited a positive correlation for most, but not all ramp rates, correlation was scarce for Na_v_1.3 and Na_v_1.6. Note that smaller Na_v_1.5 sample sizes due to the established exclusion criteria might limit its correlation accuracy. Our results indicate that ramp and window currents correlate only in specific cases and are thus likely to be generated by distinct mechanisms.

### All tested VGSCs activate during the AP subthreshold phase except for Na_v_1.8

We used single AP recordings from human iPSC-derived nociceptors (control cells from Meents et al. (2019)) as voltage command to determine the contribution of each VGSC to the phases of the AP. The three pre-recorded APs (AP1, AP2 and AP3) differ in their subthreshold properties, their width and size of overshoot (Fig. 7A/C/E; Table 1).

**Figure 7:**
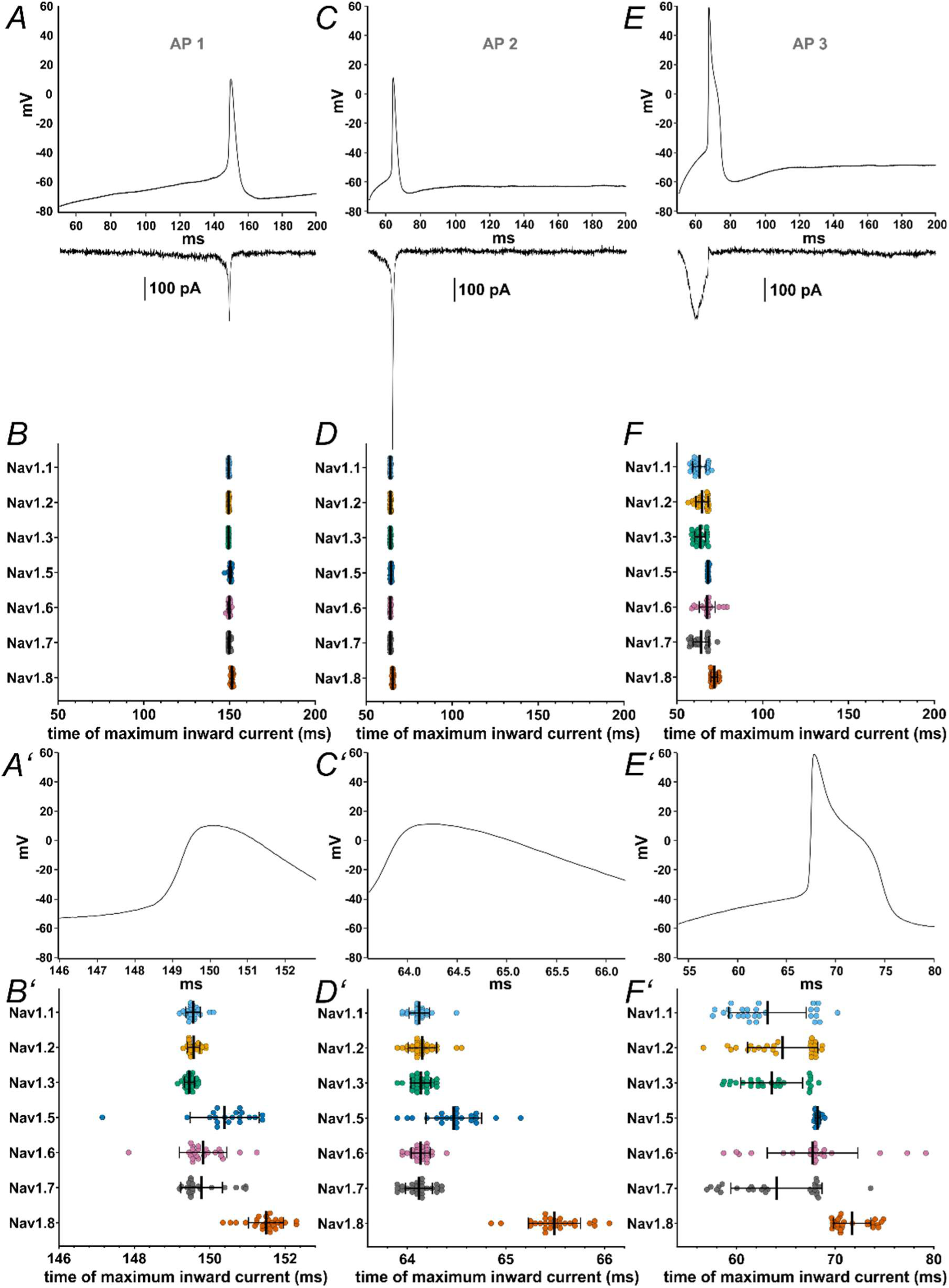
Maximum Inward Current Occurs at Different Timepoints During Action Potential Clamping for Each VGSC Isoform. *A/C/E* action potential waveforms (AP1 – AP3, respectively) recorded from iPSC-derived nociceptors used as voltage commands in AP clamping, with representative current traces, all from the same Na_v_1.7 cell, charted underneath. *B/D/F* timepoint at which the maximum inward current during the action potential (AP1 – AP3, respectively) occured. *A’-F’* zoomed in depictions of figures *A-F* (excluding the current example traces). Data are shown as mean ± SD. Statistical significance from multiple comparisons has not been indicated in the figure panels for readability purposes but can be consulted in Table S13.

The timepoint during the AP at which the maximum inward current is elicited differed between certain VGSC isoforms for all AP commands (Fig. 7B/D/F; Table S12 and S13; *p* < 0.0001 for all Kruskal-Wallis tests), as did the maximum inward current sizes (Fig. 8A-C, Table S12, *p* < 0.0001 for all Kruskal-Wallis tests) and the area under the current response curve (Fig. 8D-F, Table S12; *p* < 0.0001 for all Kruskal-Wallis tests).

**Figure 8:**
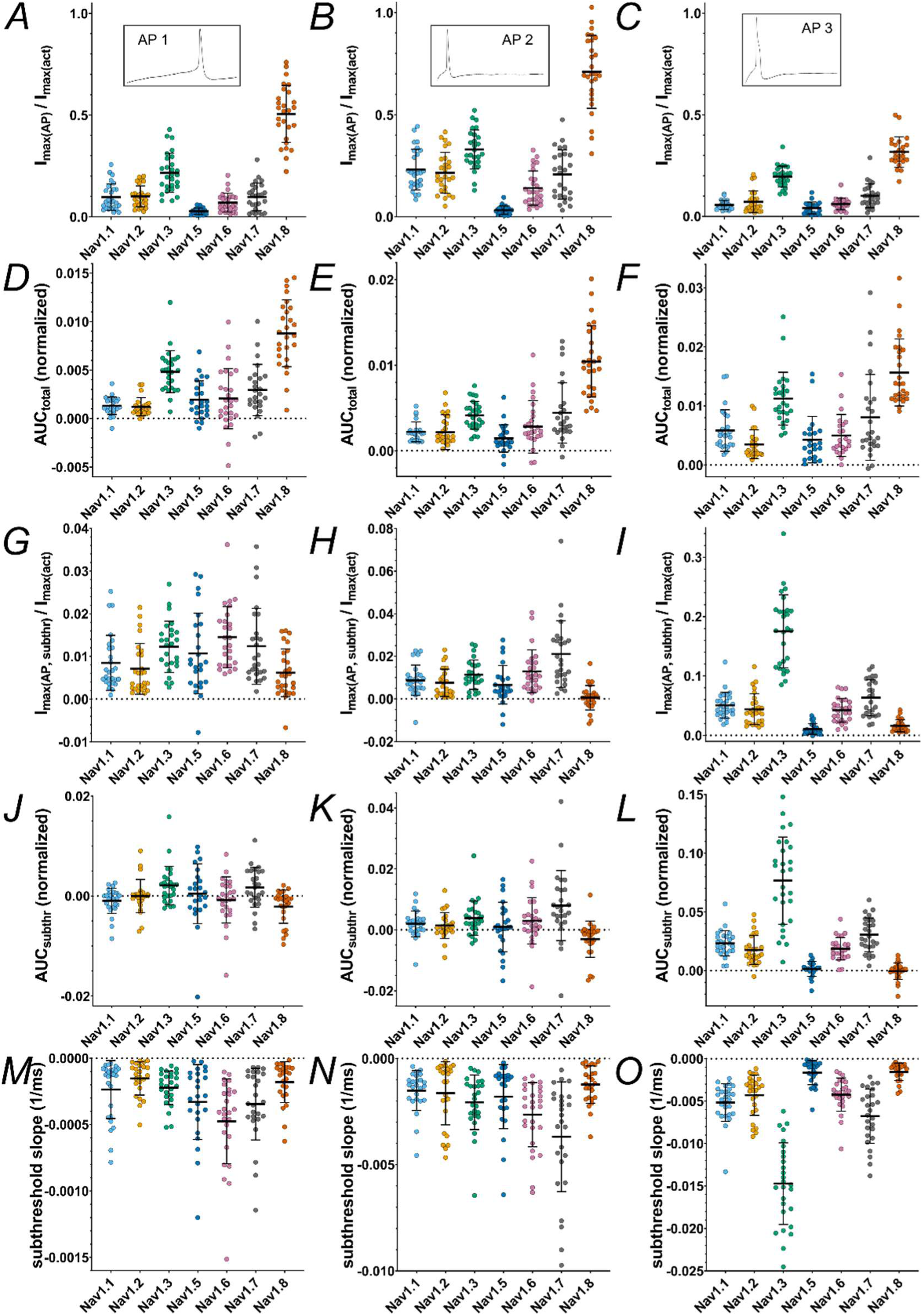
Current and AUC Parameters Obtained From Action Potential Clamping. *A-C* maximum inward current during the total command action potential (AP1-AP3, respectively) (inlets indicate the action potential command). *D-F* area under the current response curve to the total command action potential (AP1-AP3, respectively). *G-I* maximum inward current during the subthreshold phase of the command action potential (AP1-AP3, respectively). *J-L* area under the current response curve to the subthreshold phase of the command action potential (AP1-AP3, respectively). *M-O* slope of the inward current measured during the subthreshold phase of the command action potential (AP1-AP3, respectively). Data are normalized to the maximum inward current of activation measurement of the correspondent cell and shown as mean ± SD. Statistical significance from multiple comparisons has not been indicated in the figure panels for readability purposes but can be consulted in Table S13.

Multiple comparisons revealed that Na_v_1.8 currents were elicited at later timepoints than all other tested VGSC isoforms except for Na_v_1.5, namely after the AP peak (Table S13). Na_v_1.5 currents occurred slightly later within the AP time course as well, whereas Na_v_1.3 leaned towards earlier timepoints in the subthreshold phase (Fig. 7B’/D’F’, Table S12 and S13). Current sizes and AUC sizes were similarly distributed between the AP commands, with Na_v_1.8 and Na_v_1.3 exhibiting larger and Na_v_1.5 exhibiting slightly lower values for maximum inward current (Fig. 8A-F, Table S12 and S13).

The examination of the timepoint of maximum inward current during AP3 stood out since some VGSC isoforms displayed results with a bimodal distribution (Fig. 7F’). Post-hoc data revision revealed a transient current peak during the upstroke of the AP command in varying degrees for Na_v_1.1-1.3 and Na_v_1.7, most likely caused by insufficient pipette capacitance compensation. When this transient current exceeded the more or less evenly pronounced subthreshold influx, the measured timepoint was shifted, forming the second cluster depicted in Fig. 7F’. For Na_v_1.8, which also displayed a slightly clustered distribution, no similar effect could be observed. By excluding the measurements impeded by above mentioned transients, channel distribution became even more distinct: Na_v_1.1-1.3 and Na_v_1.7 had their maximum influx during the subthreshold phase of the AP command, considerably before Na_v_1.5, Na_v_1.6 and Na_v_1.8 (Fig. S2, Table S13).

During the subthreshold phase of the APs maximum inward current, AUC and inward current slope differed between the VGSC isoforms (*p* < 0.0001 for all Kruskal-Wallis tests except for AP1 subthreshold AUC (*p* = 0.0008)). On average, maximum inward currents increased with increasing subthreshold slope of the AP commands (Fig. 8G-I, Table S12). Post-hoc testing indicates that Na_v_1.8 and Na_v_1.5 elicited little to no current during the subthreshold phase, while maximum inward current was higher for Na_v_1.3 (AP1 and AP3), Na_v_1.7 (AP2 and AP3), and, for AP1, Na_v_1.6 (Fig. 8G-I, Table S13). Especially for AP3 with the steepest subthreshold slope, Na_v_1.3 currents increased drastically (Fig. 8I, Table S12 and S13). The remaining TTXs isoforms had intermediate maximum inward current responses.

A similar pattern was observed for the subthreshold phase AUC (Fig. 8J-L, Table S12 and S13). Na_v_1.5 and Na_v_1.8 AUC values remained close to zero for all AP commands (note that negative AUC values occur due to undulation of the current base rate and/or drift). Na_v_1.7 and Na_v_1.3 displayed larger subthreshold AUC values (the latter particularly for AP3), thereby adding a larger charge to a cell during subthreshold depolarization and potentially alleviating initiation of an AP.

As another measure of channel contribution to subthreshold depolarization, we determined the slope of currents elicited during the subthreshold phase of AP clamping by each VGSC isoform (Fig. 8M-O, Table S12 and S13). Steeper subthreshold slopes were documented for Na_v_1.7 (AP2 and AP3), Na_v_1.6 (AP1 and AP2), and, with emphasis on AP3, Na_v_1.3 compared to the other tested channels. In contrast, Na_v_1.5 and Na_v_1.8 subthreshold slopes were minor. The data are in line with the assumption that Na_v_1.8 is mainly activated during the upstroke of a nociceptor AP, while TTXs channels (predominantly Na_v_1.7, Na_v_1.6, and – especially for steeper subthreshold depolarizations – Na_v_1.3) contribute in earlier, subthreshold phases.

### *In silico* simulation suggests significant contribution of Na_v_1.9 to the fast upstroke of the AP and its shoulder

With the large data set in hand, which contains patch-clamp recordings of all VGSCs relevant in nociceptors, we now aimed to obtain insights into the contribution of each isoform to the AP by *in silico* experiments. Our computational model comprises the sodium currents of Na_v_1.1 – Na_v_1.3, Na_v_1.5 – Na_v_1.8 (derived from the current trace measurements of this study), Na_v_1.9 (derived from current trace measurements of Leipold et al. (2013) and Leipold et al. (2015)) and a generic potassium and leak current (see Methods and Supplements).

In order to generate an adequate fit of the recorded current traces of the different VGSCs, we implemented a modification to the original Hodgkin-Huxley model: we introduced a waiting state after pulse onset due to which the fast inactivation gate (denoted as *h* in the Hodgkin-Huxley framework) is delayed (see Methods and Supplements). This waiting state could mimic the time during which VGSC may open and are not ready to inactivate yet. This modification led to improved trace fitting for Na_v_1.1-1.3 and Na_v_1.5-1.8, as can be seen in the reduced deviation from the original trace (see Supplements). The Hodgkin-Huxley-like models of each sodium channel subtype were integrated into a non-spatial model of an Aδ-fiber and a CMi-fiber, which were used to simulate APs in response to current injections (see Methods and Supplements).

To model CMi-fibers (Fig. 9/S3) and Aδ-fibers (Fig. 10/S4), we accounted for the fiber-specific expression patterns of VGSCs quantified in Tavares-Ferreira et al. (2022), as these sequencing results provide well accessible, reliable human expression data of all voltage-gated sodium channels in sensory neurons.

**Figure 9:**
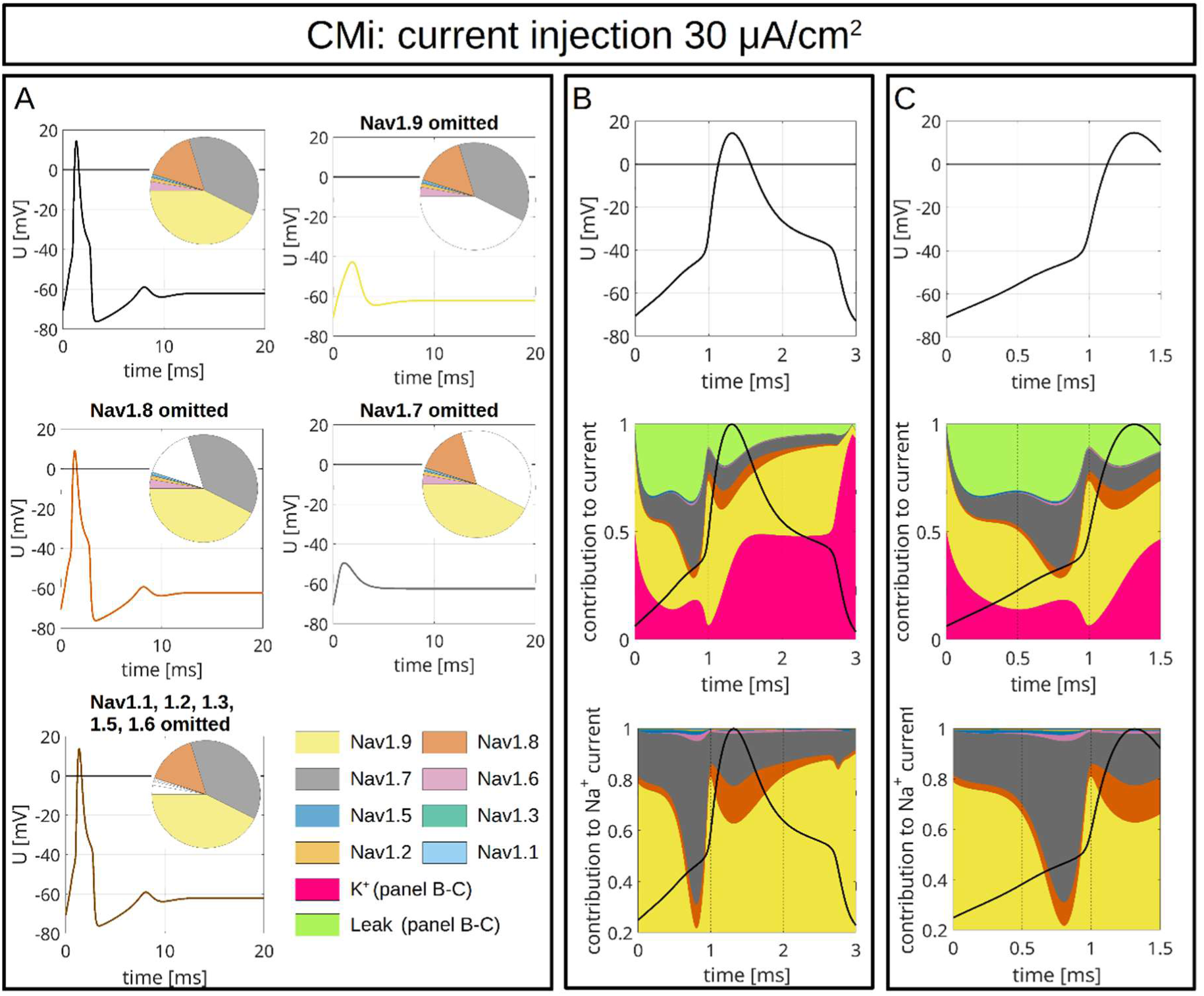
Computer Simulations of APs in a CMi-fiber. The pie charts indicate the abundance of different VGSC isoforms. APs are triggered by an injection of 30 μA/cm² which persists until the end of the simulation. *A* The upper left panel shows the AP for the relative abundance of VGSC isoforms quantified in spatial gene expression. The initial condition of the simulation is the resting state of the fiber. The other panels show changes of the AP resulting from removal of Na_v_1.9 (upper right), Na_v_1.8 (middle left), Na_v_1.7 (middle right) and Na_v_1.1-1.3, 1.5 & 1.6 (bottom left). The VGSC isoforms are coded by the colors specified in the bottom right panel. *B/C* VGSC isoform contributions to the simulated AP depicted in the upper left panel of *A.* The upper panels of *B* and *C* show the membrane potential during the initial 3 ms *(B)* and 1.5 ms *(C)* of the simulated AP. The middle panels show the relative contribution of the different sodium currents, the potassium current and the leak current over the course of the AP, plotted as stacked individual currents normalized to total current at each time point. The bottom panels show VGSC isoform contributions to the total sodium current, plotted as stacked individual VGSC isoform currents normalized to the total sodium current at each time point. The black line visualizes the shape of the AP. The color coding is identical to that in *A*.

**Figure 10:**
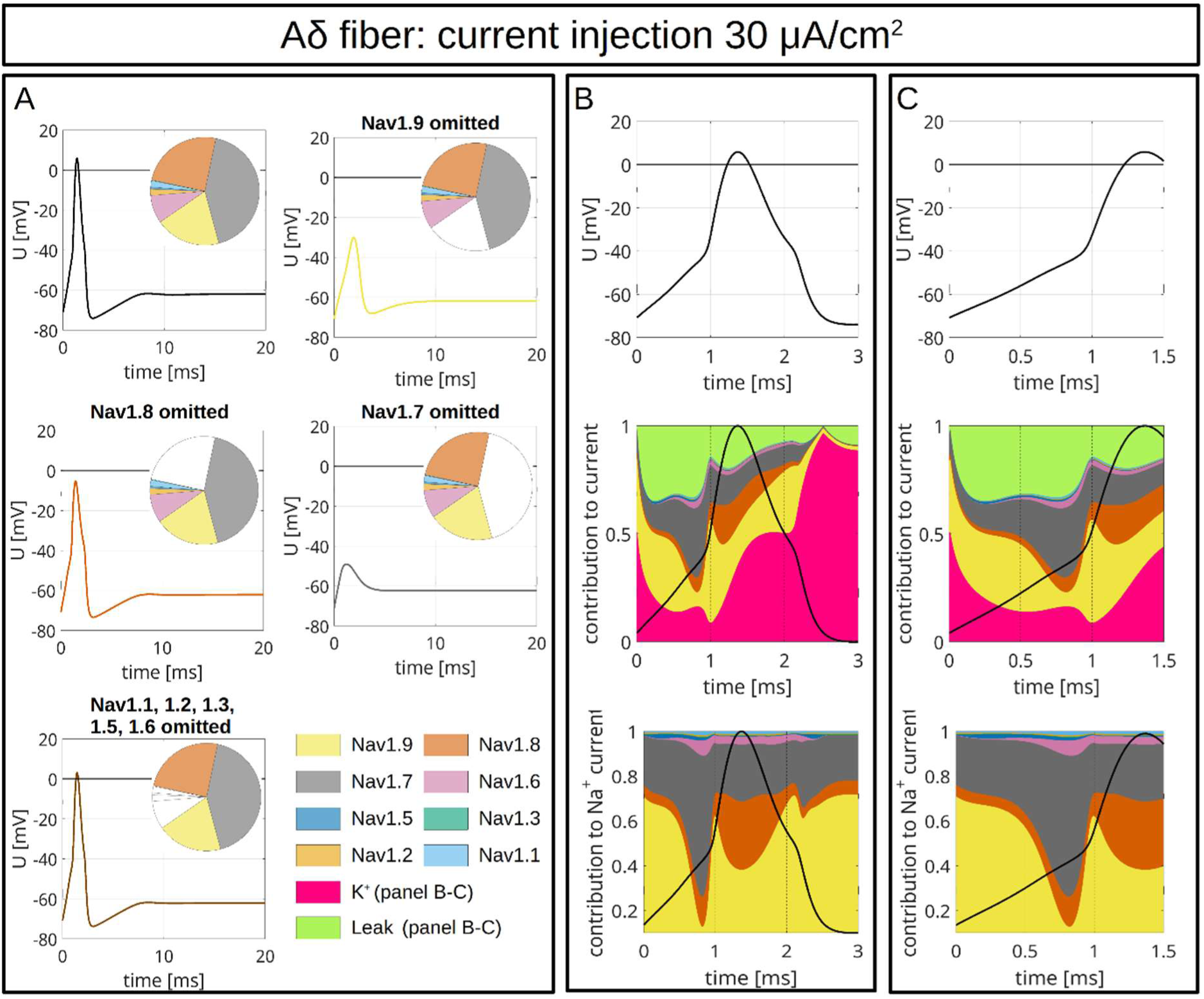
Computer Simulations of APs in a Aδ-fiber. The pie charts indicate the abundance of different VGSC isoforms. APs are triggered by an injection of 30 μA/cm² which persists until the end of the simulation. *A* The upper left panel shows the AP for the relative abundance of VGSC isoforms quantified in spatial gene expression. The initial condition of the simulation is the resting state of the fiber. The other panels show changes of the AP resulting from removal of Na_v_1.9 (upper right), Na_v_1.8 (middle left), Na_v_1.7 (middle right) and Na_v_1.1-1.3, 1.5 & 1.6 (bottom left). The VGSC isoforms are coded by the colors specified in the bottom right panel. *B/C* VGSC isoform contributions to the simulated AP depicted in the upper left panel of *A*. The upper panels of *B* and *C* show the membrane potential during the initial 3 ms (*B*) and 1.5 ms (*C*) of the simulated AP. The middle panels show the relative contribution of the different sodium currents, the potassium current and the leak current over the course of the AP, plotted as stacked individual currents normalized to total current at each time point. The bottom panels show VGSC isoform contributions to the total sodium current, plotted as stacked individual VGSC isoform currents normalized to the total sodium current at each time point. The black line visualizes the shape of the AP. The color coding is identical to that in *A*.bottom left). The VGSC isoforms are coded by the colors specified in the bottom right panel. *B/C* VGSC isoform contributions to the simulated AP depicted in the upper left panel of *A*. The upper panels of *B* and *C* show the membrane potential during the initial 3 ms (*B*) and 1.5 ms (*C*) of the simulated AP. The middle panels show the relative contribution of the different sodium currents, the potassium current and the leak current over the course of the AP, plotted as stacked individual currents normalized to total current at each time point. The bottom panels show VGSC isoform contributions to the total sodium current, plotted as stacked individual VGSC isoform currents normalized to the total sodium current at each time point. The black line visualizes the shape of the AP. The color coding is identical to that in *A*.

The maximal total voltage-gated sodium, potassium and leak conductivities were chosen to recapitulate the experimentally observed AP morphology. In the simulations, the APs were triggered by current injection of 30 µA/cm² (Fig. 9/10) or 40 µA/cm² (Fig. S3/S4) throughout the duration of the experiment. Both fiber types revealed an AP with overshoot and a shoulder (stronger for CMi-fibers) upon stimulation with 30 µA/cm² (Fig. 9A/10A) and 40 µA/cm² (Fig. S3A/S4A).

To assess the role of a specific VGSC isoform, we removed it from the model and observed the resulting AP changes (Fig. 9A/10A/S3A/S4A/S5). Notably, the removal of an isoform resulted in a reduction of the total sodium conductance.

The most striking finding of our simulations is the strong contribution of Na_v_1.9 to the fast upstroke of the AP as well as to the formation of a shoulder in both fiber types (Fig. 9B/C, 10 B/C, S3B/C, S4B/C, S5, S6). Omission of Na_v_1.9 at low stimulation intensities aborted AP generation, very similar to omission of Na_v_1.7 (Fig. 9A/10A). The latter simulates *loss-of-function* as it may occur for Na_v_1.7 in patients suffering from chronic insensitivity to pain (Lampert et al., 2014). While removal of Na_v_1.8 from the model resulted in AP waveforms that failed to overshoot, this intervention did not affect formation of the AP shoulder, suggesting that other subtypes such as Na_v_1.9 may contribute to shoulder formation (Fig. 9A/10A).

To investigate this further, we reduced the conductance of Na_v_1.9 gradually in our CMi-model, while increasing that of the other sodium channels, to keep the overall VGSC conductance constant (Fig. S6). Reducing Na_v_1.9 gradually flattened the shoulder; the fast upstroke of the AP turned less steep and subthreshold depolarizations following the AP disappeared. Additionally, the afterhyperpolarization was reduced, which may reflect a weaker activation of voltage-gated potassium channels due to the vanishing of the shoulder. Removal of Na_v_1.1 – 1.3, Na_v_1.5 and Na_v_1.6 (at the same time) had no significant impact on the AP of the CMi-fiber but reduced the overshoot in the Aδ-fiber (Fig. 9A/10A/S3A/S4A/S5).

Fig. 9B/C and 10B/C show the relative contributions of the different VGSC isoforms to the simulated APs of the CMi-fiber and the Aδ-fiber, respectively. In both fiber types the relative contribution of Na_v_1.7 is maximal right before or at the beginning of the fast upstroke of the AP (Fig. 9C, 10C, S3C, S4C), in line with reports on Na_v_1.7 as subthreshold or threshold channel (Rush et al., 2007; Vasylyev and Waxman, 2012; Meents et al., 2019). The largest relative (Fig. 9B/C, 10B/C), but also absolute (Fig. S7) conduction by far in our two models is mediated by Na_v_1.9. Most of its relative contribution occurs during subthreshold depolarizations, then during the fast upstroke of the AP (relative and absolute), and, with a prominent second peak (relative and absolute), during the falling phase of the AP coinciding with the shoulder.

Na_v_1.8, on the other hand, shows largest conductivity during the upstroke, the peak and the falling phase of the AP (Fig. 9B/C, 10B/C, S3B/C, S4B/C, S7), but when the AP shoulder occurs, it seems to be already partially inactivated. The relative and absolute contribution of Na_v_1.8 and Na_v_1.7 to the AP, especially during the peak, seems to be flipped between the two fiber models investigated here: while in the CMi-fiber model Na_v_1.7 contributes more, it is Na_v_1.8 in the Aδ-fiber model. The other channels show only minor contributions during the course of the AP: Na_v_1.5 shows some activity during subthreshold depolarizations and a small peak during the shoulder in the CMi-fiber model, while Na_v_1.6 is active at the AP threshold in both models. Na_v_1.6 is continuously active throughout the AP in the Aδ model (Fig. 9B/C, 10B/C, S3B/C, S4B/C, S7).

*Gain-of-function* mutations of Na_v_1.7 linked to inherited pain syndromes often modify channel gating such that during an AP more channels are activated. Erythromelalgia mutations in Na_v_1.7 lead to hyperpolarized activation of the channel, inducing hyperexcitability when overexpressed in rodent sensory neurons (Körner and Lampert, 2020) and enhance spontaneous activity in human stem cell derived nociceptors (Mis et al., 2019). We used our computational model to simulate *gain-of-function* in two ways: 1) by increasing the maximal conductance of Na_v_1.7 and comparing it to an increased Na_v_1.8 conductance and 2) by shifting activation of Na_v_1.7 to more hyperpolarized potentials.

When we increased Na_v_1.7 conduction by 5 times in our computer model, both, the CMi-fiber model and the Aδ-fiber model showed persistent firing in response to an ongoing current injection of 1 µA/cm² (Fig. 11A/12A). Interestingly, enhancing Na_v_1.8 conduction by 11 times (revealing similar overall conductance as for a five-fold increased Na_v_1.7 conductance), was unable to induce ongoing AP firing (Fig. 11B/12B). This finding supports the concept that the specific properties of Na_v_1.7 and not the increase of the total sodium conductance trigger the repetitive activity observed in Fig. 11A and 12A.

**Figure 11:**
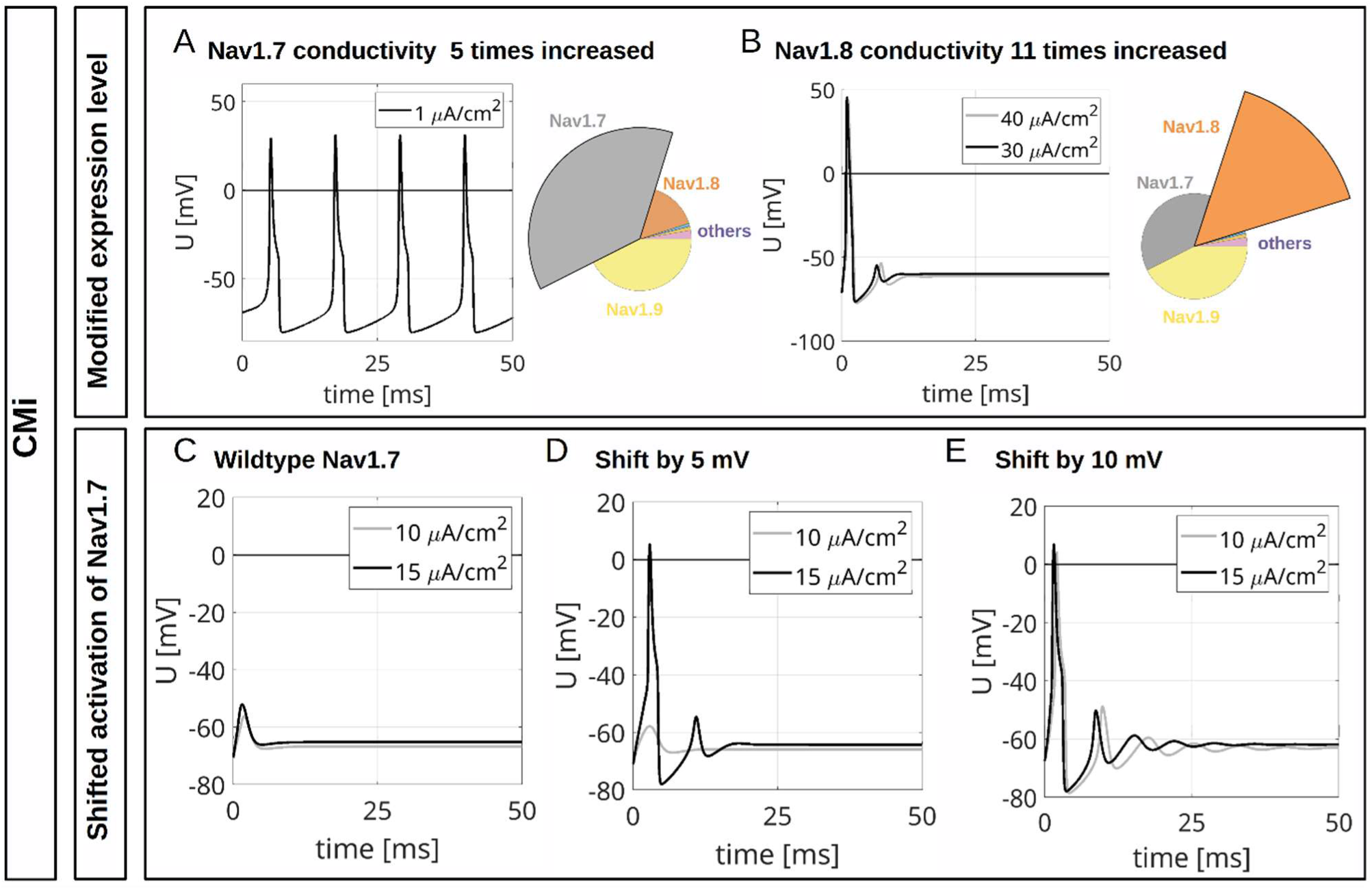
Computational Disease Modelling in a CMi-fiber. *A* Repetitive depolarizations resulting from increasing the maximal conductance for Na_v_1.7 to 500% of the wildtype value. The current injection of 1 μA/cm² persists until the end of the simulation. *B* An increase of the maximal conductance for Na_v_1.8 to 1100% of the wildtype value results in an isolated AP. The total sodium conductances in *A* and *B* are approximately the same. The current injections of 30 μA/cm² and 40 μA/cm² persist until the end of the simulations. The pie charts indicate the abundance of different VGSC isoforms. *C* Current injections of 10 μA/cm² and 15 μA/cm² trigger no APs in the simulated wildtype CMi (isoform abundances as in Fig 9A, upper left panel). *D* If the activation of Na_v_1.7 is shifted by 5 mV to hyperpolarized potentials, a current injection of 15 μA/cm² but not of 10 μA/cm² triggers an AP. *E* If the activation of Na_v_1.7 is shifted by 10 mV to hyperpolarized potentials, current injections of 10 μA/cm² and 15 μA/cm² trigger an AP. The current injections in *C-E* persist until the

**Figure 12:**
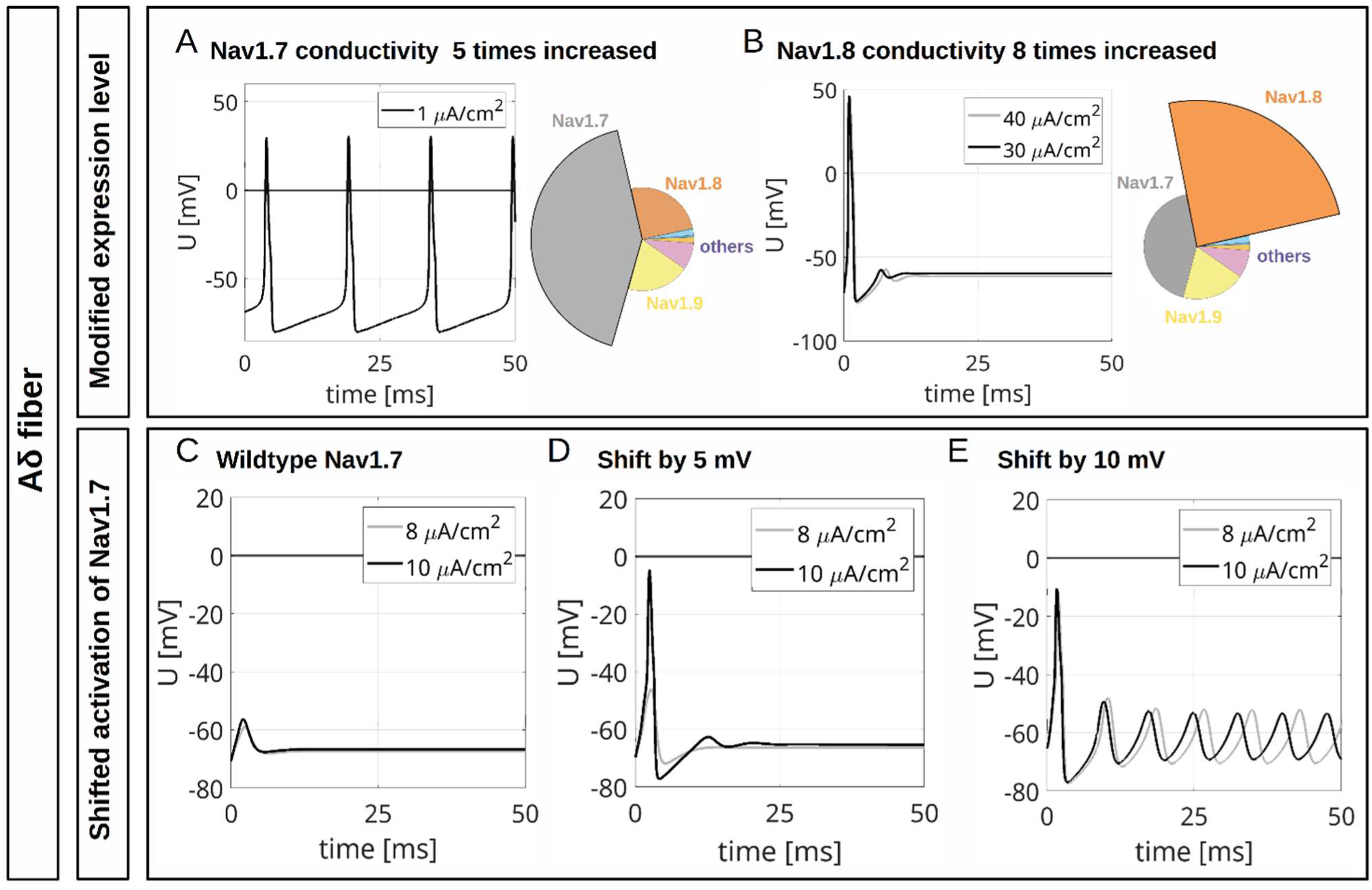
Computational Disease Modelling in an Aδ fiber. *A* Repetitive depolarizations resulting from increasing the maximal conductance for Na_v_1.7 to 500% of the wildtype value. The current injection of 1 μA/cm² persists until the end of the simulation. *B* An increase of the maximal conductance for Na_v_1.8 to 800% of the wildtype value results in an isolated AP. The total sodium conductances in *A* and *B* are approximately the same. The current injections of 30 μA/cm² and 40 μA/cm² persist until the end of the simulations. The pie charts indicate the abundance of different VGSC isoforms. *C* Current injections of 10 μA/cm² and 15 μA/cm² trigger no APs in the simulated wildtype CMi (isoform abundances as in Fig 10A, upper left panel). *D* If the activation of Na_v_1.7 is shifted by 5 mV to hyperpolarized potentials, a current injection of 15 μA/cm² but not of 10 μA/cm² triggers an AP. *E* If the activation of Na_v_1.7 is shifted by 10 mV to hyperpolarized potentials, current injections of 10 μA/cm² and 15 μA/cm² trigger an AP. The current injections in *C-E* persist until the

With unshifted Na_v_1.7 activation, subthreshold stimulations of 10 or 15 µA/cm² did not evoke an AP in the CMi-fiber model (Fig. 11C), and neither did subthreshold stimulations of 8 or 10 µA/cm² in the Aδ-fiber model (Fig. 12C). When Na_v_1.7 activation was shifted to hyperpolarized potentials by only 5 mV – which is often observed in erythromelagia mutations (e.g., Lampert et al. (2009)) – an AP was initiated by small stimulations, showing a reduced firing threshold (Fig. 11D/12D). When shifted by 10 mV, weak stimulations even induced an unstable membrane potential leading to several strong, repetitive depolarizations (Fig. 11E/12E).

Taken together, our simulations including Hodgkin-Huxley-like models of all nociceptive sodium channels suggest a contribution of Na_v_1.9 to the fast upstroke of the AP and its shoulder formation and proved useful in modelling excitability changes associated with Na_v_1.7-linked chronic pain syndromes such as erythromelalgia.

## Discussion

In this study, we provide a detailed, comparative investigation of the gating properties of all VGSCs relevant in nociceptors with focus on subthreshold depolarizations and propose computational models to predict their contributions to AP electrogenesis in two types of sensory neurons: CMi-fibers and Aδ-fibers. The models suggest that Na_v_1.9 may actively contribute to AP generation and formation of its shoulder in these neurons.

### VGSC isoforms are characterized by their distinct gating behavior *in vitro* and *in silico*

The TTXr channels measured in this study, i.e., Na_v_1.5 and Na_v_1.8, showed divergent gating properties with shifted voltage dependence of activation, SSFI and ramp currents toward hyperpolarized and depolarized voltages, respectively. While Na_v_1.9 activation occurred at very hyperpolarized voltages, voltage dependence of Na_v_1.9 SSFI resembled the depolarized inactivation of Na_v_1.8. Out of the remaining TTXs channels, which all had their voltage dependence of activation, SSFI and ramp currents at subthreshold voltages, especially Na_v_1.3 and Na_v_1.7 elicited prominent currents in ramp and AP clamping. During the subthreshold phases of AP clamping, both displayed large AUC values and steeper current slopes compared to the other VGSCs tested. For Na_v_1.7, we observed slightly more hyperpolarized voltage dependences for both, SSFI and ramp currents, than for other TTXs channel isoforms, potentially enabling Na_v_1.7 to charge a cell at subthreshold or threshold voltages.

Our experiments emphasized the importance of Na_v_1.7 as AP initiator, also since knocking out or enhancing Na_v_1.7 *in silico* led to omitted or multiple APs, respectively. This resembles putative *loss-* and *gain-of-function* pathophysiology that is attributed to Na_v_1.7 channelopathies (Dib-Hajj et al., 2005; Cox et al., 2006; Faber et al., 2012; Bennett et al., 2019; McDermott et al., 2019). Shifting activation of Na_v_1.7 to more hyperpolarized potentials in models for CMi-or Aδ-fibers revealed the experimentally observed threshold reduction, suggesting the high potential of our simulations for disease modelling.

Our findings confirmed Na_v_1.8 contribution to the AP upstroke (Renganathan et al., 2001; Matsutomi et al., 2006; Rush et al., 2007; Vasylyev and Waxman, 2012), as it activates and fast-inactivates at more depolarized potentials, elicits large ramp currents at more depolarized potentials and showed little to no subthreshold activity in AP clamping compared to the other VGSCs tested. Interestingly, in our simulations including Na_v_1.9, the contribution of Na_v_1.8 to the shoulder was minor: knocking out Na_v_1.8 in the *in silico* experiments left APs in the CMi-model almost unchanged, while in the Aδ-model the overshoot was reduced. This is in line with findings from Na_v_1.8 knockout DRGs, where maximum AP voltages were lower compared to wildtype DRGs (Renganathan et al., 2001; Harty and Waxman, 2007). Yet, the initiation of the AP was not impeded when Na_v_1.8 was omitted from the established model.

The strongest difference between the two fibers modelled in this study is the inverse relative contribution of Na_v_1.7 compared to Na_v_1.8 during the AP: for the CMi-fiber model, Na_v_1.7 has the larger current and conductivity, whereas for the Aδ-fiber model it is Na_v_1.8. In an earlier modelling study, we investigated the contribution of Na_v_1.7, Na_v_1.8 and Na_v_1.9 to the excitability of a peripheral C-fiber, to reproduce axonal spike propagation speed and activity-dependent slowing as recorded in microneurography experiments (Tigerholm et al., 2014; Tigerholm et al., 2015). We showed that the speed of two APs following each other depends on the influx of sodium through Na_v_1.7 relative to Na_v_1.8 current. Thus, fiber specific biophysical properties may result from the specific contribution of the sodium channels expressed in their membrane.

Our simulations suggest that Na_v_1.9, in addition to its activation during rest, strongly contributes to the fast upstroke and is a major determinant of the shoulder of the AP. It represents by far the largest current and conductivity in our model of all integrated channels.

The findings are in line with current-clamp recordings on enteric neurons from Na_v_1.9 knockout mice which show no shoulder in APs from mice lacking Na_v_1.9 (Osorio et al., 2014). An early report on the sensory neuron AP properties of those knockout mice revealed no differences in their AP width (Priest et al., 2005). The latter study did not differentiate between sensory neuron subtypes, yet in the realm of recent transcriptomic advances we know that Na_v_1.9 expression differs strongly between subgroups (Tavares-Ferreira et al., 2022). It is also known that not all sensory neuron subtypes display a shoulder in their AP (Körner and Lampert, 2022). Thus, pooling the groups may have hidden the effect of Na_v_1.9 on the AP shoulder in this study. As Na_v_1.9 channelopathies are linked to both gain-and loss-of-pain syndromes (Leipold et al., 2013; Huang et al., 2017), clarification of the fiber type-specific role of Na_v_1.9 during AP generation could help to understand their complex pathophysiology.

We documented large Na_v_1.3 currents during clamping of an AP with steep subthreshold voltage slope (AP3), and – consistent with literature – during steeper voltage ramps (Cummins et al., 2001). This might play a role in neuropathic pain, as – though only marginally expressed in C-and Aδ-fibers – some studies suggest that Na_v_1.3 is upregulated in spinal cord injuries or peripheral nerve injuries in rodents and therefore increases excitability (Hains et al., 2003; Hains et al., 2004; Lampert et al., 2006). Other studies present more ambiguous or even contradictory results, e.g., Na_v_1.3 knockout mice did not show lowered pain thresholds after nerve injury than wildtype mice (Lindia et al., 2005; Nassar et al., 2006).

Na_v_1.6 stood out to some extent, as it showed higher maximum inward currents and steeper current slopes during the subthreshold phase of AP clamping than other TTXs VGSC isoforms. Furthermore, Na_v_1.6 current was larger in the Aδ-fiber than in the CMi-fiber during the course of the AP. Yet, omitting Na_v_1.1, Na_v_1.2 and Na_v_1.6 during the *in silico* testing had no noticeable effect on AP morphology, likely because of their minor share in the spatial transcriptomics-derived expression patterns used in our model (Tavares-Ferreira et al., 2022). Na_v_1.6 has been linked to resurgent currents (O’Brien and Meisler, 2013), and these currents seem to be enhanced in A-fibers compared to C-fibers (Klinger et al., 2012). This is suggested as a reason why sea-anemone toxin ATX-II led to painful sensations by enhancing resurgent currents in A-fibers, but not in C-fibers (Klinger et al., 2012).

Na_v_1.5 played no major role during subthreshold AP clamping and in contributing to the simulated AP, while it is also only marginally detected in C-and Aδ-fibers (Tavares-Ferreira et al., 2022). Due to its hyperpolarized activation and SSFI, it may be responding in cells with more hyperpolarized membrane potential. During development, Na_v_1.5 is the third TTXr conductance present and may have a more prominent role than observed in the two fiber types described here (reflecting the transcriptomic expression pattern of adult DRG donors). The role of Na_v_1.5 during development, injury and regeneration needs to be investigated in more detail in follow-up studies (Renganathan et al., 2002; Tseng et al., 2010; Sousounis et al., 2020).

### Correlation between ramp current and window current is channel subtype-specific

Slowly depolarizing voltage ramps induce ramp currents, mimicking the subthreshold phase of an AP. Ramp currents of wild-type channels have been examined for all VGSC isoforms individually (Cummins et al., 1998; Abriel et al., 2001; Herzog et al., 2003; El-Bizri et al., 2011; Power et al., 2012; Estacion and Waxman, 2013; DeCaen et al., 2014; Han et al., 2015a; Han et al., 2015b; Zhang et al., 2017), but not yet in direct comparison, establishing experimenter-to-experimenter or lab-to-lab variations. Hereby, we provide an extensive and highly comparable data set of VGSC ramp currents.

While maximum ramp current size and AUC are predominantly influenced by the steepness of the inducing voltage ramp stimulus, VGSC subtypes rather influence the voltage dependence of ramp current responses and hence co-determine in which AP phase they contribute to its formation.

All tested TTXs channel isoforms elicited their peak ramp current at subthreshold voltages, enabling them to contribute to cell discharge prior to the AP threshold being exceeded. Of these subtypes, Na_v_1.3, Na_v_1.6 and Na_v_1.7 elicited the highest maximum ramp currents, corroborating their role as key players in subthreshold depolarization.

In VGSC literature, ramp and window currents were often assumed to describe a similar gating mechanism and were sometimes even used interchangeably (Wang et al., 1996; Magistretti and Alonso, 1999; Abriel et al., 2001; Blair and Bean, 2002; Enomoto et al., 2006; Matsutomi et al., 2006; Vasylyev and Waxman, 2012; Estacion and Waxman, 2013; Osteen et al., 2017). We here tested this hypothesis and showed that Na_v_1.1 and Na_v_1.2 (partly also Na_v_1.5, Na_v_1.7 and Na_v_1.8) showed a positive correlation between ramp current and window currents, suggesting a common underlying mechanism. Yet, because for Na_v_1.3 and Na_v_1.6 correlation was mostly non-existent, we suppose that additional factors such as persistent current play into ramp current formation, as suggested previously (Estacion and Waxman, 2013).

### Expression systems affect *in vitro* VGSC measurements

Heterologous expression systems bear their own interpretation pitfalls, as they do not resemble the complex environment of primary neurons, e.g., regarding cell morphology, intercellular contacts, or membrane protein interactions. VGSC isoforms were shown to have different gating properties when expressed in different expression systems (Cummins et al., 2001; Rush et al., 2006a; Tan and Soderlund, 2009; Zhang et al., 2017).

In this study, we used both HEK and ND7/23 cells for electrophysiological VGSC examination. Multiple differences in their respective cell characteristics lead to subtype gating differences among each other having to be analyzed carefully. While endogenous Na_v_1.6 and Na_v_1.7 currents in ND7/23 can easily be suppressed by adding TTX to the extracellular solution while measuring TTXr isoforms (Rogers et al., 2016; Lee et al., 2019), altered VGSC gating properties by endogenous expression of β1 and β3 subunits in ND7/23 cells cannot be eliminated (John et al., 2004; Lee et al., 2019). However, also HEK cells themselves express β1A subunits endogenously (Moran et al., 2000). Therefore, comparisons between VGSC α subunit measurements have to be treated carefully nonetheless and additional examination of α subunits in presence of varying β subunits is encouraged. Furthermore, other cellular proteins such as fibroblast growth factor homologous factors, microtubule-associated proteins or aquaporins have been shown to modify VGSC gating and/or current density (Wittmack et al., 2004; Rush et al., 2006b; Zhang and Verkman, 2010; O’Brien et al., 2012; Vanoye et al., 2013). Recent comparative studies of Na_v_1.2 and Na_v_1.6 variants also highlight the influence of alternative splicing on channel gating (Thompson et al., 2020; Vanoye et al., 2024).

Na_v_1.8 expression in HEK is challenging and resulting currents are low (Gladwell et al., 1998; John et al., 2004). As in our hands, expression of Na_v_1.8 in HEK293T cells yielded little to no current, we used ND7/23 cells, equivalently to the Na_v_1.9 measurements integrated into this study (Leipold et al., 2013; Leipold et al., 2015). Biophysical characteristics of Na_v_1.8 currents were shown to be similar between when expressed in ND7/23 cells or recorded from native DRG TTXr currents (John et al., 2004).

Even non-native neuronal cell lines such as ND7/23 do not resemble a comprehensive primary neuronal environment (Yin et al., 2016). Studies in expression systems closer to human peripheral sensory neurons such as rodent or human DRG neurons could give an even better insight into the channels biophysics *in vivo*. Moreover, nociceptors derived from human induced pluripotent stem cells offer the possibility to investigate ion channels in an adaption of their native environment (Meents et al., 2019; Mis et al., 2019; Namer et al., 2019). This could also make up for the fact that we included neither co-expression of β-subunits into our measurements nor did we check for channel dimerization, both of which have shown to alter biophysical properties of VGSCs (Brackenbury and Isom, 2011; Chahine and O’Leary, 2011; Clatot et al., 2017; Rühlmann et al., 2020).

Albeit our observations allow unprecedented comparison between isoforms as the experiments were conducted under the so far best possible comparable conditions, it may be that differing expression systems do not model certain isoform specificities.

Biophysical differences such as larger persistent or ramp currents can occur between VGSC isoforms from different mammalian species (Browne et al., 2009; Tan and Soderlund, 2009; Han et al., 2015a). For Na_v_1.3 and Na_v_1.6 measurements in this study, we used rodent channel isoforms. Hence, complementary measurements with human channel isoforms are desirable.

In this study as well as the studies providing the implemented Na_v_1.9 data, intracellular solution contained CsF which is widely used in patch clamp experiments to enhance high seal resistances for stable and long-lasting patch clamp configurations (Kostyuk et al., 1975; Fernandez et al., 1984). Yet, usage of CsF has been shown to affect the gating of VGSCs in a subtype-specific manner. For example, the voltage dependence of activation and inactivation of Na_v_1.9 is shifted by about 15-20 mV to more hyperpolarized potentials in presence of CsF (Rugiero et al., 2003; Coste et al., 2004). The gating of Na_v_1.3 and Na_v_1.7 is also affected by intracellular fluoride, albeit to a lesser extent (Chen et al., 2000; Meadows et al., 2002; Jarecki et al., 2008). While this does not necessarily limit subtype comparability within our *in vitro* measurements, which all included CsF, comparison to CsF-free experiments such as measurements in primary neurons have to be performed cautiously. Even minor shifts in channel biophysics can profoundly impact complex and therefore vulnerable computational models. Although the model established in this study provides valuable insights into the AP contribution of each channel, reissuing the model with data from CsF-free measurements would be desirable to increase its predictive power.

### Limitations of *in silico* modelling

Computational models are simplifications of the real world and therefore must be handled cautiously, especially when model complexity suggests the opposite. We estimated the distribution of the maximal sodium conductances per VGSC subunit by using mRNA expression data from a human DRG spatial transcriptomics study (Tavares-Ferreira et al., 2022). While this approach is closest to a quantitative measure currently accessible from human DRGs, mRNA expression levels do not always indicate that the respective channel is also translated to protein, trafficked to the cell surface and fully functional. Even though correlation between mRNA expression, presence of VGSCs in the cell membrane and biophysical properties has been suggested (Thériault and Chahine, 2014), alternative splicing, correct folding and association with additional proteins or membrane turnover are just some factors that may impact channel translation from RNA to functional protein in the membrane (Buccitelli and Selbach, 2020). It is possible that mRNA levels do not directly relate to the amount of protein produced in the cells (Lindhout et al., 2020), and that we are overestimating or underestimating the conductance of the VGSCs modelled in this study. As proteomics of human DRGs, especially for VGSCs, is not yet available as a basis for our modelling, we used the published transcriptomics data to estimate the conductivity of the VGSC subtypes integrated in our simulations.

A promising solution to these problems are single-cell transcriptomics combined with functional analysis, such as Patch-seq, which offer the ability to link expression data directly to cell physiology (Lipovsek et al., 2021; Körner et al., 2022) or single cell proteomics, which are currently developed also for human nociceptors. Furthermore, since expression data are gathered at the cell soma, only limited assumptions can be made for small nerve fibers, e.g., in the human skin.

Several studies suggested the possibility of neuronal channel composition changing in compensation for *loss-* or *gain-of-function* mutations in VGSC isoforms, e.g., TTXs currents being upregulated in DRG neurons of Na_v_1.8 knockout mice (Akopian et al., 1999; Renganathan et al., 2000). Compensatory changes could also involve the expression of other ion channel types, e.g., calcium channels, or pathways indirectly affecting excitability. When modelling the effect of VGSC *loss-* or *gain-of-function* mutations *in silico*, these compensatory changes should be considered, as soon as their extent is determined in precedent studies.

VGSC data are usually gathered at room temperature, as it was done in this study. However, temperature dependence of VGSCs varies for each channel isoform and recorded voltages (Kriegeskorte et al., 2023). Thus, most computational models published in the recent years including the one established in this study do not accurately account for temperature, since this would require additional experimental data and re-parametrization of the models (Almog and Korngreen, 2016).

For fitting the gating variables of each channel subtype, we used current traces recorded at several potentials. At voltages below -40 mV it often occurred that multiple parameter combinations described the currents successfully, creating some uncertainty (unidentifiability). Particularly in the case of Na_v_1.9 this may impact on simulation results.

Our computational model describes ion channels based on modified Hodgkin-Huxley dynamics and neglects spatial aspects. To further study the VGSC influence on AP generation, the VGSC isoforms should be implemented in a morphologically detailed C-fiber or Aδ-fiber model (Tigerholm et al., 2014; Tigerholm et al., 2019).

### VGSC isoform characterization can be the basis for assessing potential drug targets

In this study we provide a broadly based groundwork for future comparative studies of VGSC isoforms, their biophysics and interaction with each other. By feeding our measurements into a computational model, we generated an *in silico* tool to test hypotheses, investigate the contribution of each nociceptive sodium channel to AP genesis and model pharmacology and effect of disease-related mutations. This may also prove useful when developing new potential analgesics such as specific channel blockers or optimizing therapy schemes for chronic pain.

## Supporting information

in silico supplements

The following abbreviations are used:

## Abbreviations

*AP*: action potential
*AUC*: area under the curve
*CMi*: mechano-insensitive C-fiber
*CNS*: central nervous system
*CsF*: cesium fluoride
*DMEM*: Dulbecco’s modified Eagle medium
*DRG*: dorsal root ganglion
*ECS*: extracellular solution
*E_rev_*: reversal potential
*FBS*: fetal bovine serum
*G_Na_*: sodium conductance
*G_Na,max_*: maximum sodium conductance
*G418*: Geneticin disulphate
*ICS*: intracellular solution
*I_Na_*: inward sodium current
*I_Na,max_*: maximum inward sodium current
*iPSC*: induced pluripotent stem cells
*k*: slope factor
*PNS*: peripheral nervous system
*R_pip_*: pipette tip resistance
*R_s_*: series resistance
*R_seal_*: seal resistance
*SD*: standard deviation
*SSFI*: steady-state fast inactivation
*TTX*: tetrodotoxin
*TTXr*: TTX-resistant
*TTXs*: TTX-sensitive
*VGSC*: voltage-gated sodium channel
*V_hold_*: holding potential
*V_m_*: membrane potential
*V_50_*: membrane potential at half-maximal channel (in)activation.

## Additional information

### Data availability

All data gathered in this study and all analysis procedures used are available from the corresponding authors upon request.

### Disclosures/Competing interests

The authors declare no competing financial interests and no conflicts of interest with the contents of this article. A.L. had an unrelated research agreement with Grunenthal. A.L. receives counseling fees from Grunenthal.

## Acknowledgements

We thank Brigitte Hoch, Petra Hautvast, and Raya A. Bott for their excellent technical support (cell culture, molecular biology, electrophysiology). We thank Jannis Körner, Sophia Kriegeskorte and Jannis E. Meents for their thoughtful discussions as well as Jannis E. Meents for providing the AP stimuli used in this study.

## CRediT author contributions

P. A. K.: Conceptualization, Methodology, Investigation, Software, Validation, Formal analysis, Data curation, Visualization, Writing -Original draft. E. L.: Resources, Data curation, Writing - Review & Editing. J. T.: Writing - Review & Editing. A. M.: Writing - Review & Editing. B. N.: Writing - Review & Editing. T. S.: Resources, Methodology, Investigation, Software, Validation, Formal analysis, Data curation, Visualization, Writing - Review & Editing. A. L.: Supervision, Project administration, Funding acquisition, Resources, Conceptualization, Methodology, Validation, Data curation, Writing - Review & Editing.

## Funding/Grants

This work was funded by the Deutsche Forschungsgemeinschaft (DFG, German Research Foundation 363055819/GRK2415 Mechanobiology of 3D epithelial tissues (ME3T) to A.L.; 368482240/GRK2416, MultiSenses-MultiScales to A.L., LA2740/6-1 to A.L.; NA 970/5-1 to B.N.). A.L. (IZKF TN1-1/IA 532001) and B.N. were supported by a grant from the Interdisciplinary Center for Clinical Research within the Faculty of Medicine at the RWTH Aachen University.

## Notes

http://www.thomas-stiehl.de/Preprints.html

### Summary of Updates

We provide a revised version of our manuscript now including manual patch clamp data of Nav1.9 in both our biophysical examination of voltage-gated sodium channels as well as the computational Hodgkin-Huxley-like models. The new findings reveal surprising results, e.g., a contribution of Nav1.9 to the shoulder formation of an action potential is suggested by our in silico results. Furthermore, another disease modelling approach was added to the computational models by shifting Nav1.7 activation to more hyperpolarized potentials. The introduction and discussion section have been revised thoroughly.

