## Supplementary material for "Sodium channels expressed in nociceptors contribute distinctly to action potential subthreshold phase, upstroke and shoulder": in silico supplements

### Supplementary Information: An electrophysiological and *in silico* examination of voltage-gated sodium channels contributing to subthreshold depolarization in nociceptor action potentials

**5 Simulation** **17**

### 1 Data Preparation for Modeling

Conductance was calculated as in equation (1) of the main text. Measurements fulfilling $|V_m - E_{rev}| < 10mV$ were excluded to avoid error amplification resulting from division by a small number. For each individual cell the mean conductance at -120mV was subtracted as background conductance. The maximal conductance of each cell was normalized to 1. Due to capacitive artifacts the calculated conductance can be negative after changes of the clamping voltage. When this occurred we used the earliest timepoint with non-negative conductance as the starting point of the voltage-induced sodium current.

### 2 Fitting

Fitting was performed using a weighted least square approach. Variances of the calculated conductance were used as weights. To simplify the fitting we assumed that at $-120mV$ the amount of inactivated Nav channels is negligible. Furthermore we assumed that the sodium current through the Nav channels is negligible at

$-120mV$, i.e. all Nav channels are closed at $-120mV$ but can be activated upon depolarization. We fitted the gating variables for each measured voltage between $-70mV$ and $40mV$. The fitted values were continuously extended using the function

$$\chi(x) = \frac{a + b \cdot x}{c + d \cdot \exp\left(\frac{x+f}{g}\right)} + q, \tag{1}$$

where $x$ is the membrane potential in volts. For low voltages some parameters could not be determined. The reason for this is that e.g., if channels do not open the closing rate cannot be estimated. Such parameters were omitted when interpolating the fitted values by the function $\chi$. To avoid artifacts when $\chi(x)$ is close to zero, we replace $\chi(x)$ by $\max(\chi(x), 0)$ in the simulations. For Nav1.9 a correction for the liquid junction potential has to be applied to the fitted voltage dependencies. The respective rate constants for Nav1.9 are given by $x \mapsto \max(\chi(x+\phi), 0)$, where $\phi = 0.007\,V$.

### 3 Model Equations

#### 3.1 Delayed sodium channel inactivation

We have noticed that for multiple Nav-subtypes (i.e., Nav1.1-Nav1.3, Nav1.5-Nav1.8) the original Hodgkin-Huxley model could not accurately reproduce the ion current for clamping voltages above $0V$. For this reason we introduced a delayed inactivation kinetics. The simplest way to do this is to replace the equation

$$\frac{d}{dt}h = \alpha_h(V_m)(1-h) - \beta_h(V_m)h$$

by

$$\begin{aligned}
\frac{d}{dt}h_1 &= \alpha_h(V_m)h_0 - \beta_h(V_m)h_1 \\
\frac{d}{dt}h_w &= \beta_h(V_m)h_1 - \gamma_h(V_m)h_w \\
\frac{d}{dt}h_0 &= \gamma_h(V_m)h_w - \alpha_h(V_m)h_0,
\end{aligned}$$

where the state $h_1$ corresponds to the activated state, $h_0$ corresponds to the inactivated state and $h_w$ corresponds to a state in which the channel transits towards inactivation, but is still activated, see Supplemental Figure 1.

The $h$ in the original Hodgkin-Huxley model from [1] would then be replaced by $h_1 + h_w$. Mathematically, the waiting time in the state $h_w$ in the above equations is exponentially distributed if $V_m$ is constant. It turned out that such an exponentially distributed waiting state led to only minor improvements. Therefore, we introduced a transition state with non-exponential waiting time by sequentially concatenating multiple

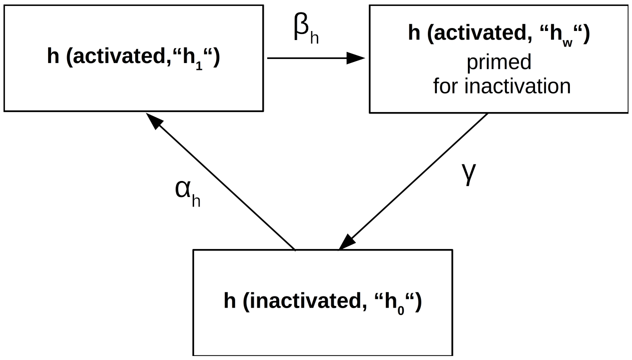

Supplementary Figure 1: Extension of the Hodgkin-Huxley model. We assume that inactivation occurs with a delay. The activated channels enter a waiting state $h_w$, in which they are still activated but primed for inactivation which occurs at rate $\gamma$.

states with exponential waiting time given by

$$\begin{aligned}
\frac{d}{dt}h_1 &= \alpha_h(V_m)h_0 - \beta_h(V_m)h_1 \\
\frac{d}{dt}h_{w,1} &= \beta_h(V_m)h_1 - \gamma_h(V_m)h_{w,1} \\
\frac{d}{dt}h_{w,i} &= \gamma_h(V_m)h_{w,i-1} - \gamma_h(V_m)h_{w,i} \qquad (2 \le i \le k) \\
\frac{d}{dt}h_0 &= \gamma_h(V_m)h_{w,k} - \alpha_h(V_m)h_0,
\end{aligned}$$

where the variable $h$ in the original model is replaced by

$$\tilde{h} := h_1 + \sum_{i=1}^{k} h_{w,i} = 1 - h_0. \tag{2}$$

This leads to a more plateau-like waiting time distribution, see Supplemental Figure 2. For $k = 5$ we found the best agreement between model and data. Supplemental Figures 3-9 compare the original model to the modified model.

For Nav1.9 the original Hodgkin-Huxley framework yielded accurate fits. Therefore, we use the standard Hodgkin-Huxley equations for Nav1.9.

#### 3.2 Rationale of the modified Hodgkin-Huxley model

In the original Hodgkin-Huxley model $\frac{d}{dt}h$ is a linear function. Therefore, in the case of voltage clamping experiments, $h(t)$ is given by a function of the type $h_0 - \Delta_h(1 - e^{-t/\tau_h})$ with $\tau_h > 0$. This has two immediate consequences: (i) Whenever the clamping voltage changes, the conductance of the ion channel changes immediately, i.e., the time required for conformational changes of the subunits is neglected, (ii) the change of $h$, i.e., $\frac{d}{dt}h$, is maximal at the moment when the clamping voltage changes. Therefore, a significant fraction

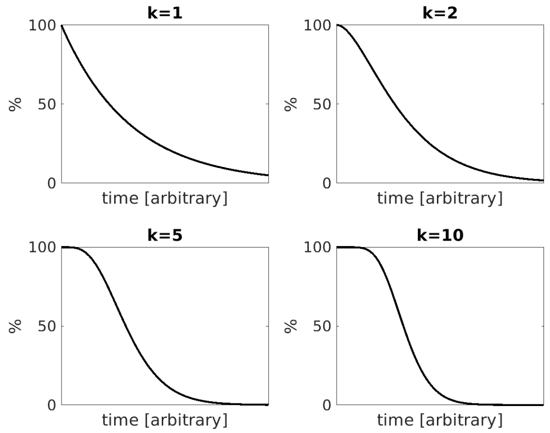

Supplementary Figure 2: Illustration of the time spent in the waiting state. The figure shows which percentage of the channels which are initially in the waiting state remain there as time progresses. In the illustration there is no influx to the waiting state, since it is meant to visualize after which period of time channels exit the state. The shape of this function depends on $k$. If we wait sufficiently long, practically all channels have left the waiting state. For $k = 1$ the decline of the channels in the waiting state is exponential for larger $k$ the shape of the function changes.

of the sodium channels inactivate prematurely, i.e., deactivate before they have been activated. This implies that the peak sodium conductance occurs shortly after the change of the clamping voltage and that at the time of the peak conductance a significant proportion of the channels is already inactivated.

In the modified model, the inactivation term $h$ accounts for the fact that the conformational changes leading to channel inactivation take some time. The change of $h$ is relatively small in the immediate aftermath of the voltage change, then it increases before it finally decreases again, see Supplementary Figure 2. Therefore, the peak conductance can occur later after the voltage change compared to the original model and a smaller fraction of channels is inactivated at the time of the peak current. The specific properties of the delay (distribution of waiting time before inactivation) is fine-tuned by the parameters $\gamma_h$ and $k \in \mathbb{N}$, where $\gamma_h$ can be voltage dependent. The parameter $k$ determines the shape of the distribution, the parameter $\gamma_h$ the time scale. The smaller the value of $\gamma_h$, the longer the delay before inactivation. The following table compares the modified to the original model.

| | **Original model** | **Modified Model** |
|---|---|---|
| time required for conformational changes leading to inactivation | neglected | considered |
| speed of inactivation ($\frac{d}{dt}h$) | fastest immediately after the change of clamping voltage | steadily increases after voltage change then decreases |
| onset of inactivation | immediately after voltage change | delayed |
| fraction of inactivated channels at the time of peak current | higher compared to modified model | lower compared to original model |
| equation for $h$ | $\frac{d}{dt}h = \alpha_h(V_m)(1-h) - \beta_h(V_m)h$ | $h = 1 - h_0$, where<br>$\frac{d}{dt}h_1 = \alpha_h(V_m)h_0 - \beta_h(V_m)h_1$<br>$\frac{d}{dt}h_{w,1} = \beta_h(V_m)h_1 - \gamma_h(V_m)h_{w,1}$<br>$\frac{d}{dt}h_{w,i} = \gamma_h(V_m)h_{w,i-1} - \gamma_h(V_m)$,<br>where $(2 \leq i \leq k)$<br>$\frac{d}{dt}h_0 = \gamma_h(V_m)h_{w,k} - \alpha_h(V_m)h_0$, |
| number of voltage-dependent parameters to describe inactivation | 2 $(\alpha_h, \beta_h)$ | 3 $(\alpha_h,\ \beta_h,\ \gamma_h)$ |
| total number of parameters to describe inactivation | 2 voltage dependent $(\alpha_h, \beta_h)$ | 3 voltage dependent $(\alpha_h,\ \beta_h,\ \gamma_h)$,<br>1 voltage independent $(k \in \mathbb{N})$ |

##### 3.3 Full model

For the change of the membrane potential $V_m$ we use the following equation in analogy to Hodgkin and Huxley:

$$
\begin{aligned}
\frac{d}{dt}V_m = \frac{I}{C_m} &- \frac{g_K}{C_m}n^4(V_m - V_K) - \frac{g_l}{C_m}(V_m - V_l) - \frac{g_{1.1}}{C_m}m_{1.1}^3\tilde{h}_{1.1}(V_m - V_{Na}) \\
&- \frac{g_{1.2}}{C_m}m_{1.2}^3\tilde{h}_{1.2}(V_m - V_{Na}) - \frac{g_{1.3}}{C_m}m_{1.3}^3\tilde{h}_{1.3}(V_m - V_{Na}) - \frac{g_{1.5}}{C_m}m_{1.5}^3\tilde{h}_{1.5}(V_m - V_{Na}) \\
&- \frac{g_{1.6}}{C_m}m_{1.6}^3\tilde{h}_{1.6}(V_m - V_{Na}) - \frac{g_{1.7}}{C_m}m_{1.7}^3\tilde{h}_{1.7}(V_m - V_{Na}) - \frac{g_{1.8}}{C_m}m_{1.8}^3\tilde{h}_{1.8}(V_m - V_{Na}) \\
&- \frac{g_{1.9}}{C_m}m_{1.9}^3 h_{1.9}(V_m - V_{Na}),
\end{aligned}
$$

where $C_m$ denotes the membrane capacitance, $g_K$ the maximal potassium conductance, $g_{1.x}$ the maximal conductance of Nav1.x ($x \in \{1, 2, 3, 5, 6, 7, 9\}$), $g_l$ the leak conductance, $V_m$ the membrane potential, $V_l$ the leak reversal potential, $V_K$ the potassium reversal potential and $V_{Na}$ the sodium reversal potential and $I$ is a current which can be applied.

The gating variables are subjected to the following ordinary differential equations, where the index 1.x refers to Nav1.x.

$$\frac{d}{dt}n = \alpha_n(V_m)(1-n) - \beta_n(V_m)n \tag{3}$$

$$\frac{d}{dt}m_{1.9} = \alpha_{m,1.9}(V_m)(1-m_{1.9}) - \beta_{m,1.9}(V_m)m_{1.9} \tag{4}$$

$$\frac{d}{dt}h_{1.9} = \alpha_{h,1.9}(V_m)(1-h_{1.9}) - \beta_{h,1.9}(V_m)h_{1.9} \tag{5}$$

$$\frac{d}{dt}m_{1.x} = \alpha_{m,1.x}(V_m)(1-m_{1.x}) - \beta_{m,1.x}(V_m)m_{1.x} \qquad x \in \{1,2,3,5,6,7,8\} \tag{6}$$

and $\tilde{h}_{1.x} = 1 - h_{0,1.x}$, fulfilling

$$\frac{d}{dt}h_{1,1.x} = \alpha_{h,1.x}(V_m)h_{0,1.x} - \beta_{h,1.x}(V_m)h_{1,1.x} \tag{7}$$

$$\frac{d}{dt}h_{w,1,1.x} = \beta_{h,1.x}(V_m)h_{1,1.x} - \gamma_{h,1.x}(V_m)h_{w,1,1.x} \tag{8}$$

$$\frac{d}{dt}h_{w,i,1.x} = \gamma_{h,1.x}(V_m)h_{w,i-1,1.x} - \gamma_{h,1.x}(V_m)h_{w,i,1.x} \qquad (2 \leq i \leq 5) \tag{9}$$

$$\frac{d}{dt}h_{0,1.x} = \gamma_{h,1.x}(V_m)h_{w,k,1.x} - \alpha_{h,1.x}(V_m)h_{0,1.x} \tag{10}$$

for each $x \in \{1,2,3,6,7,8\}$. We set

$$\tilde{h}_{1.x}(V_m) := h_{1,1.x} + \sum_{i=1}^{k} h_{w,i,1.x} = 1 - h_{0,1.x}. \tag{11}$$

### 4 Parameters

#### 4.1 Electrical Properties and Gating

The membrane capacitance was set to $C_m = 1uF/cm^2$ in agreement with [1, 3]. The reversal potentials were set to $V_{Na} = 68.97mV$ and $V_K = -81.56mV$, corresponding to the Nernst potentials for the sodium and potassium concentrations used in [3]. The conductivities and the leak potential were tuned by hand to obtain sufficient agreement between simulations and observations. We set $g_{Na} := g_{1.1} + g_{1.2} + g_{1.3} + g_{1.6} + g_{1.7} + g_{1.8} + g_{1.9} = 46mS/cm^2$, $g_K = 72mS/cm^2$, $V_l = -70.61mV$ and $g_l = 1mS/cm^2$.

The gating variables for the potassium channel were taken from [1], taking into account that potentials reported in the original work by Hodgkin and Huxley have to be transformed to contemporary units using the transformation $p \mapsto -p - 60mV$. We shifted the voltage by additional $5mV$ to obtain better agreement

**Original Hodgkin-Huxley** **Modified Hodgkin-Huxley**

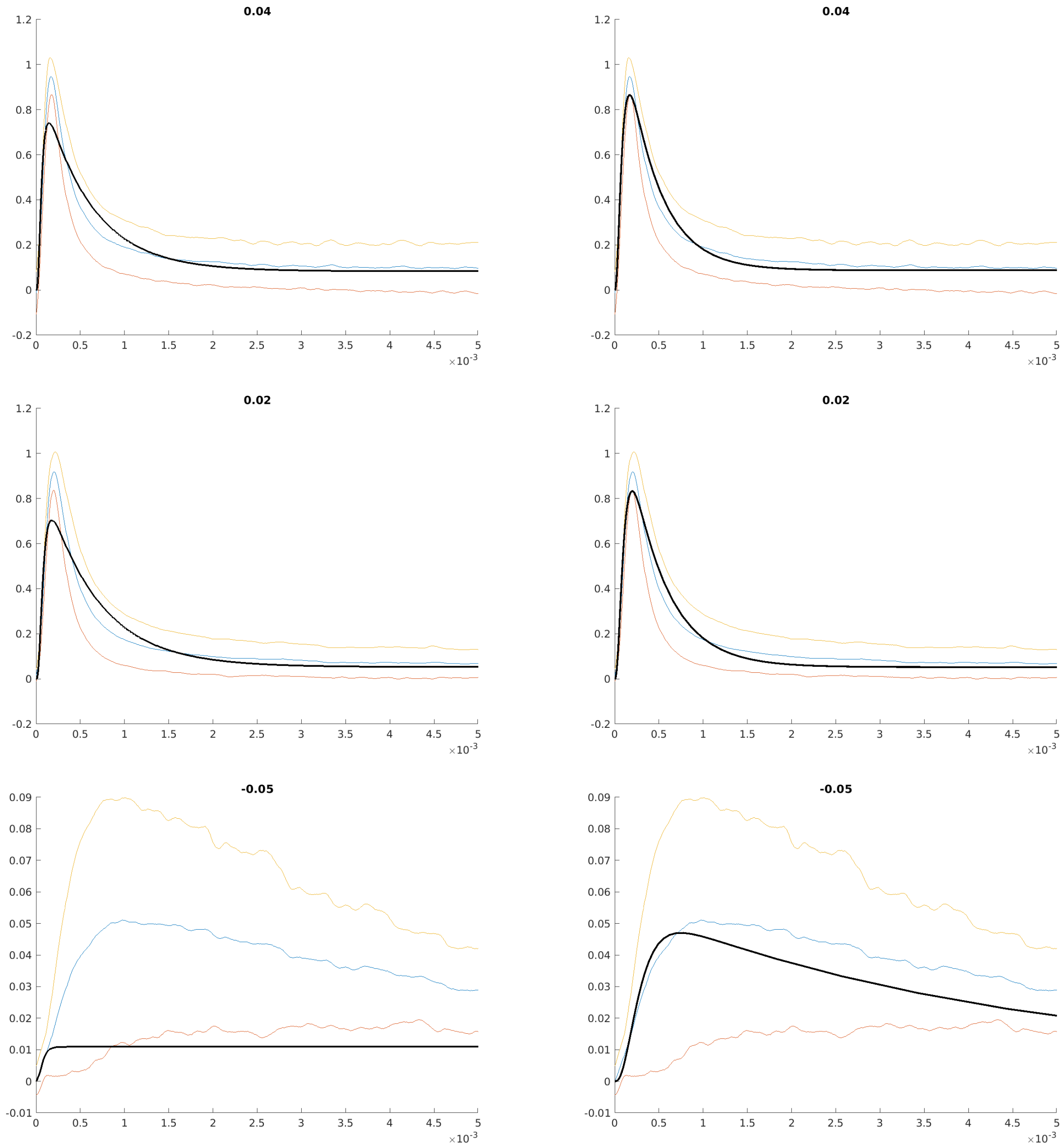

Supplementary Figure 3: Comparison of original and modified model for Nav1.1. The horizontal axis shows the time in $s$, the vertical axis the normalized sodium current. Panel titles show the clamping voltage in $volts$. Blue: average of measured cells, yellow and orange: average ± standard deviation. The plots show the current during the initial $5ms$, however, the fit was performed for the full duration of the measurement (approx. $40ms$).

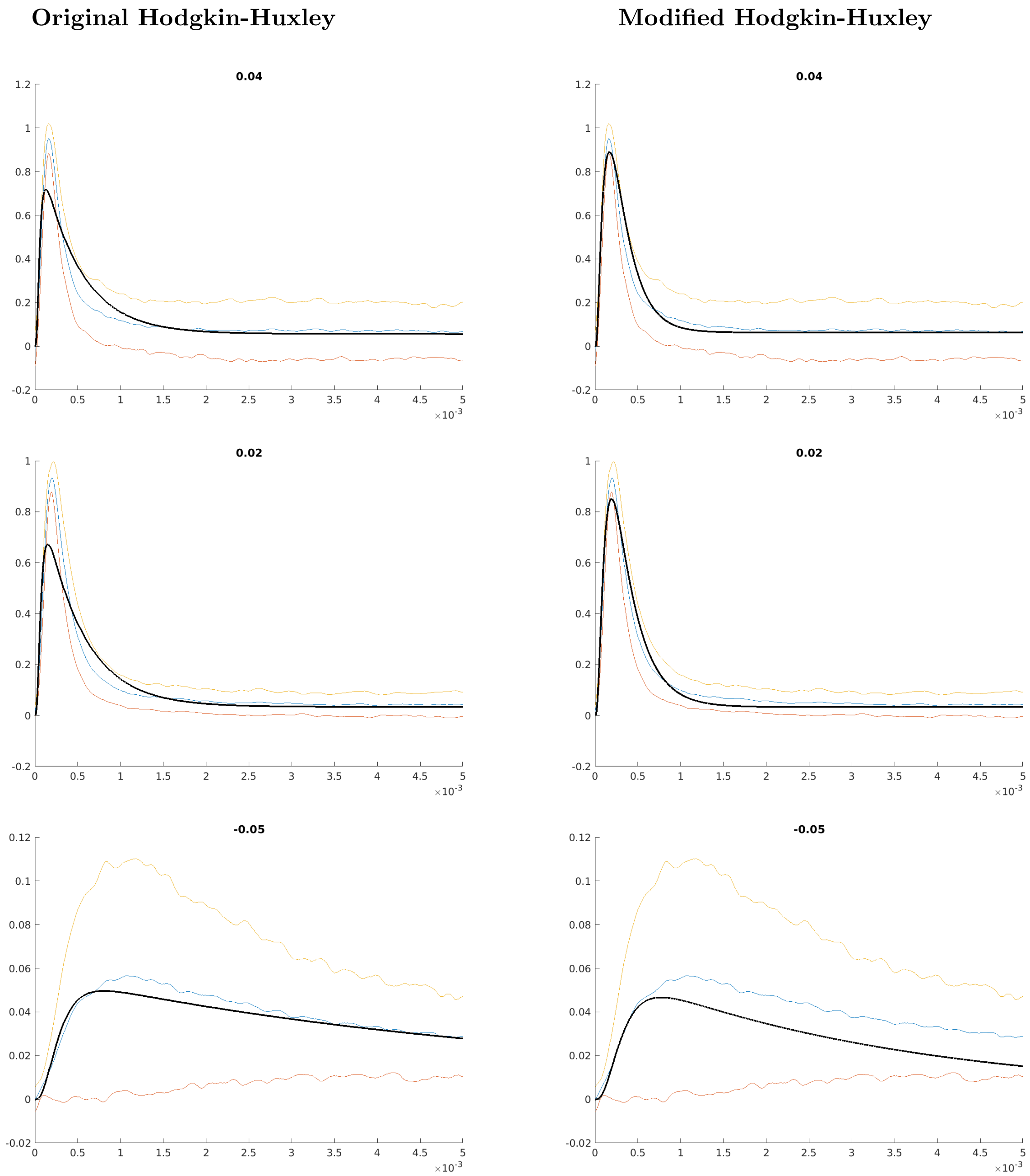

Supplementary Figure 4: Comparison of original and modified model for Nav1.2. The horizontal axis shows the time in $s$, the vertical axis the normalized sodium current. Panel titles show the clamping voltage in $volts$. Blue: average of measured cells, yellow and orange: average $\pm$ standard deviation. The plots show the current during the initial $5ms$, however, the fit was performed for the full duration of the measurement (approx. $40ms$).

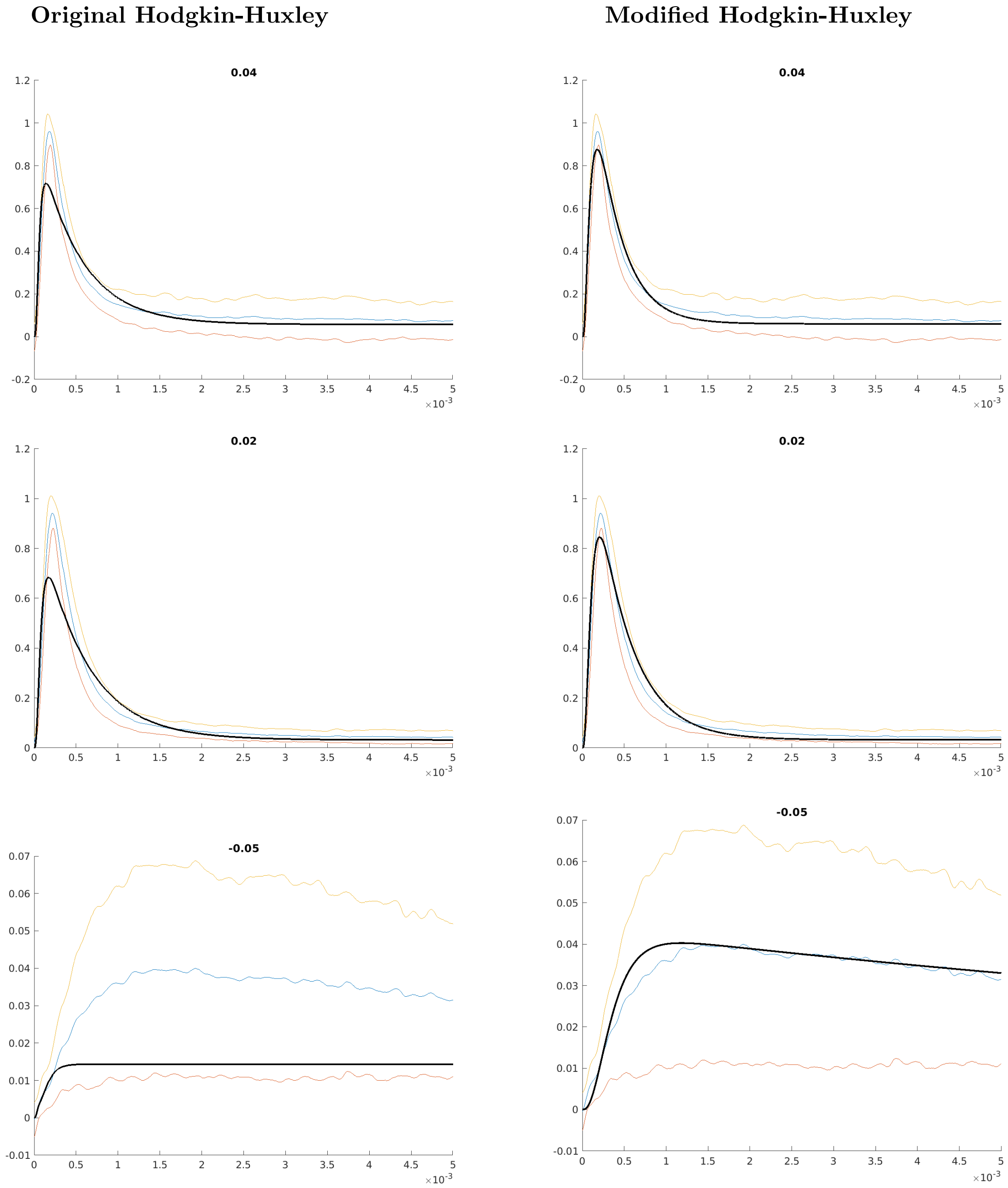

Supplementary Figure 5: Comparison of original and modified model for Nav1.3. The horizontal axis shows the time in $s$, the vertical axis the normalized sodium current. Panel titles show the clamping voltage in $volts$. Blue: average of measured cells, yellow and orange: average $\pm$ standard deviation. The plots show the current during the initial $5ms$, however, the fit was performed for the full duration of the measurement (approx. $40ms$).

**Original Hodgkin-Huxley** **Modified Hodgkin-Huxley**

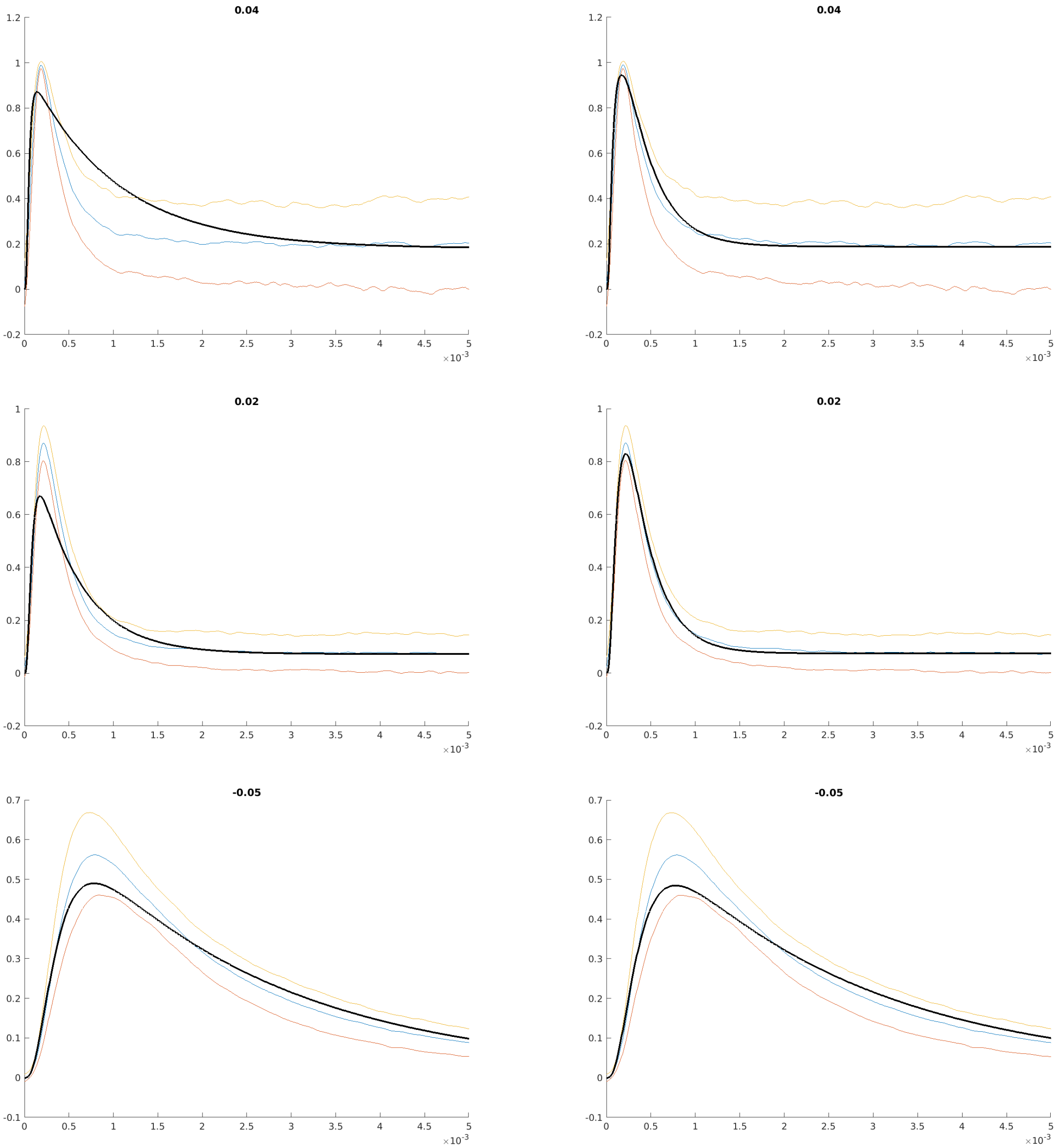

Supplementary Figure 6: Comparison of original and modified model for Nav1.5. The horizontal axis shows the time in $s$, the vertical axis the normalized sodium current. Panel titles show the clamping voltage in $volts$. Blue: average of measured cells, yellow and orange: average $\pm$ standard deviation. The plots show the current during the initial $5ms$, however, the fit was performed for the full duration of the measurement (approx. $40ms$).

**Original Hodgkin-Huxley** **Modified Hodgkin-Huxley**

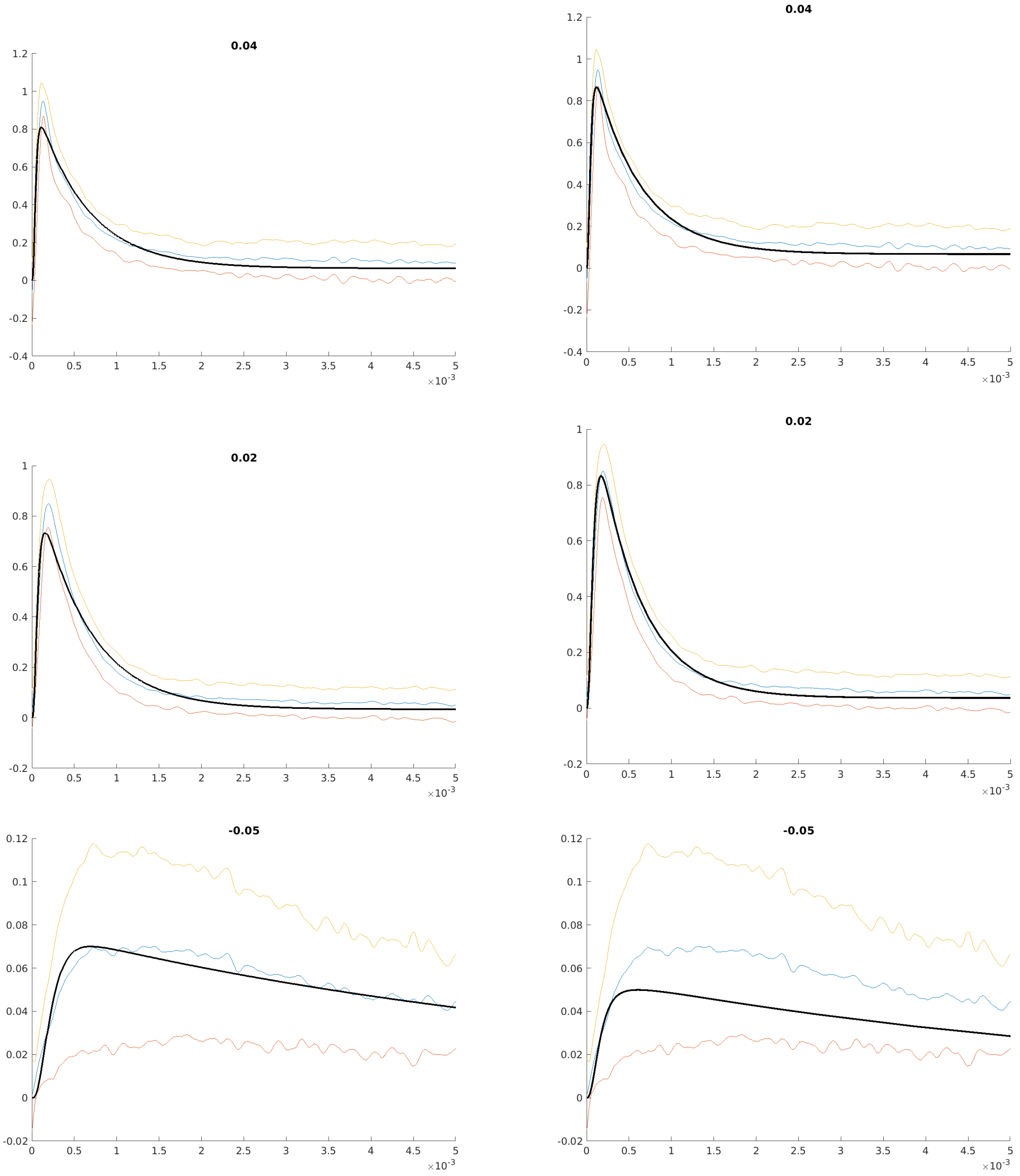

Supplementary Figure 7: Comparison of original and modified model for Nav1.6. The horizontal axis shows the time in $s$, the vertical axis the normalized sodium current. Panel titles show the clamping voltage in $volts$. Blue: average of measured cells, yellow and orange: average ± standard deviation. The plots show the current during the initial $5ms$, however, the fit was performed for the full duration of the measurement (approx. $40ms$).

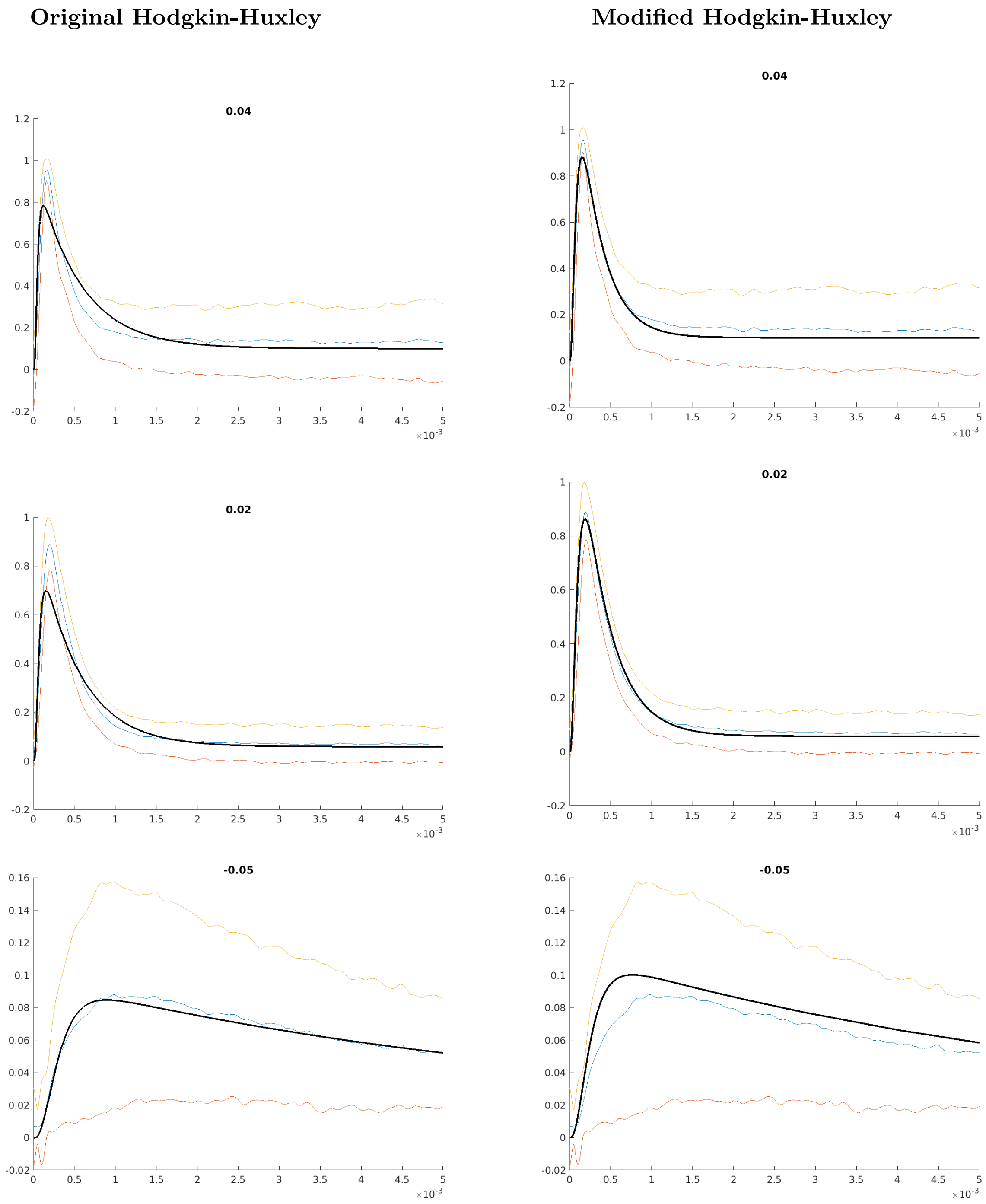

Supplementary Figure 8: Comparison of original and modified model for Nav1.7. The horizontal axis shows the time in $s$, the vertical axis the normalized sodium current. Panel titles show the clamping voltage in $volts$. Blue: average of measured cells, yellow and orange: average $\pm$ standard deviation. The plots show the current during the initial $5ms$, however, the fit was performed for the full duration of the measurement (approx. $40ms$).

**Original Hodgkin-Huxley** **Modified Hodgkin-Huxley**

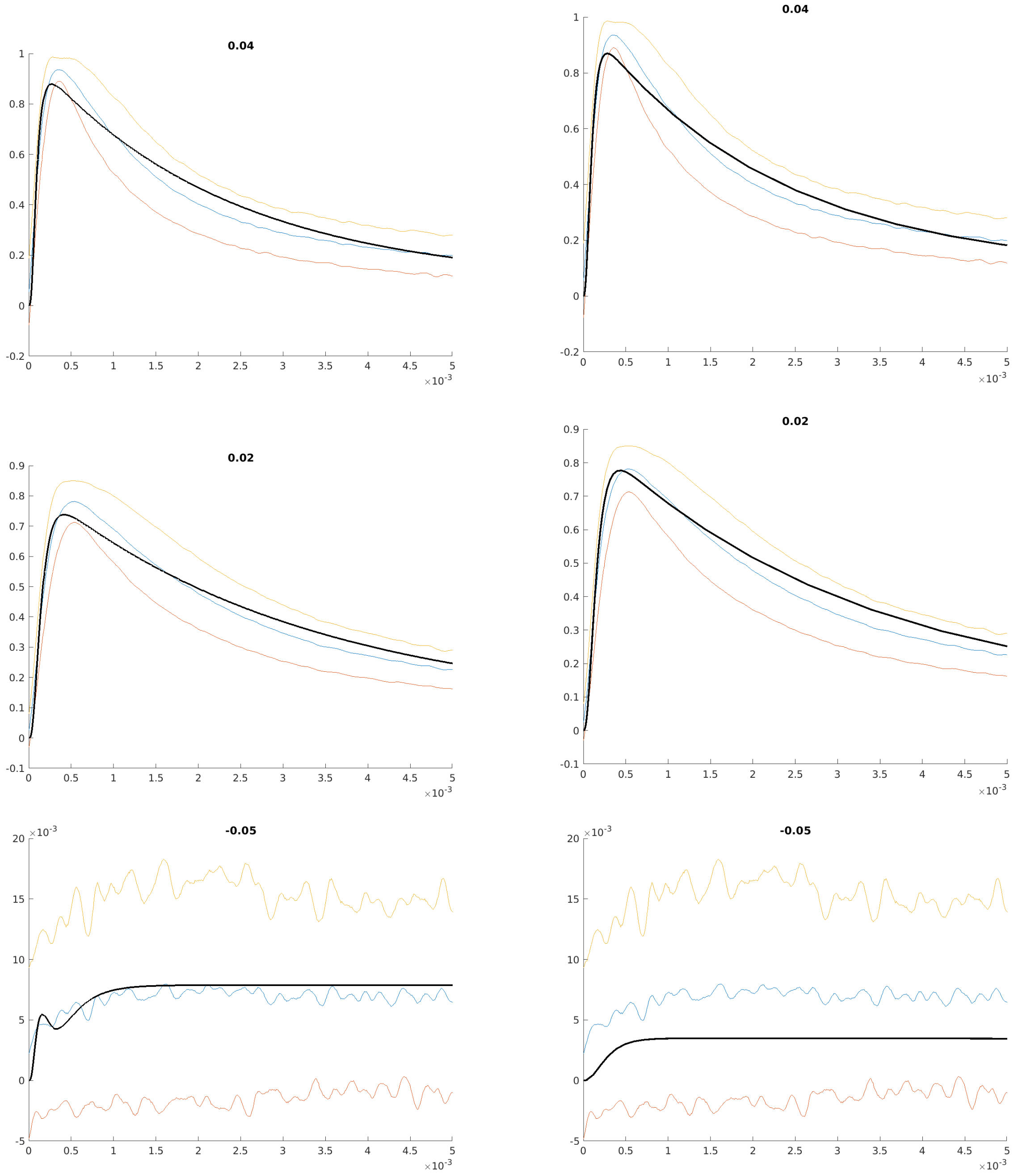

Supplementary Figure 9: Comparison of original and modified model for Nav1.8. The horizontal axis shows the time in $s$, the vertical axis the normalized sodium current. Panel titles show the clamping voltage in $volts$. Blue: average of measured cells, yellow and orange: average $\pm$ standard deviation. The plots show the current during the initial $5ms$, however, the fit was performed for the full duration of the measurement (approx. $40ms$).

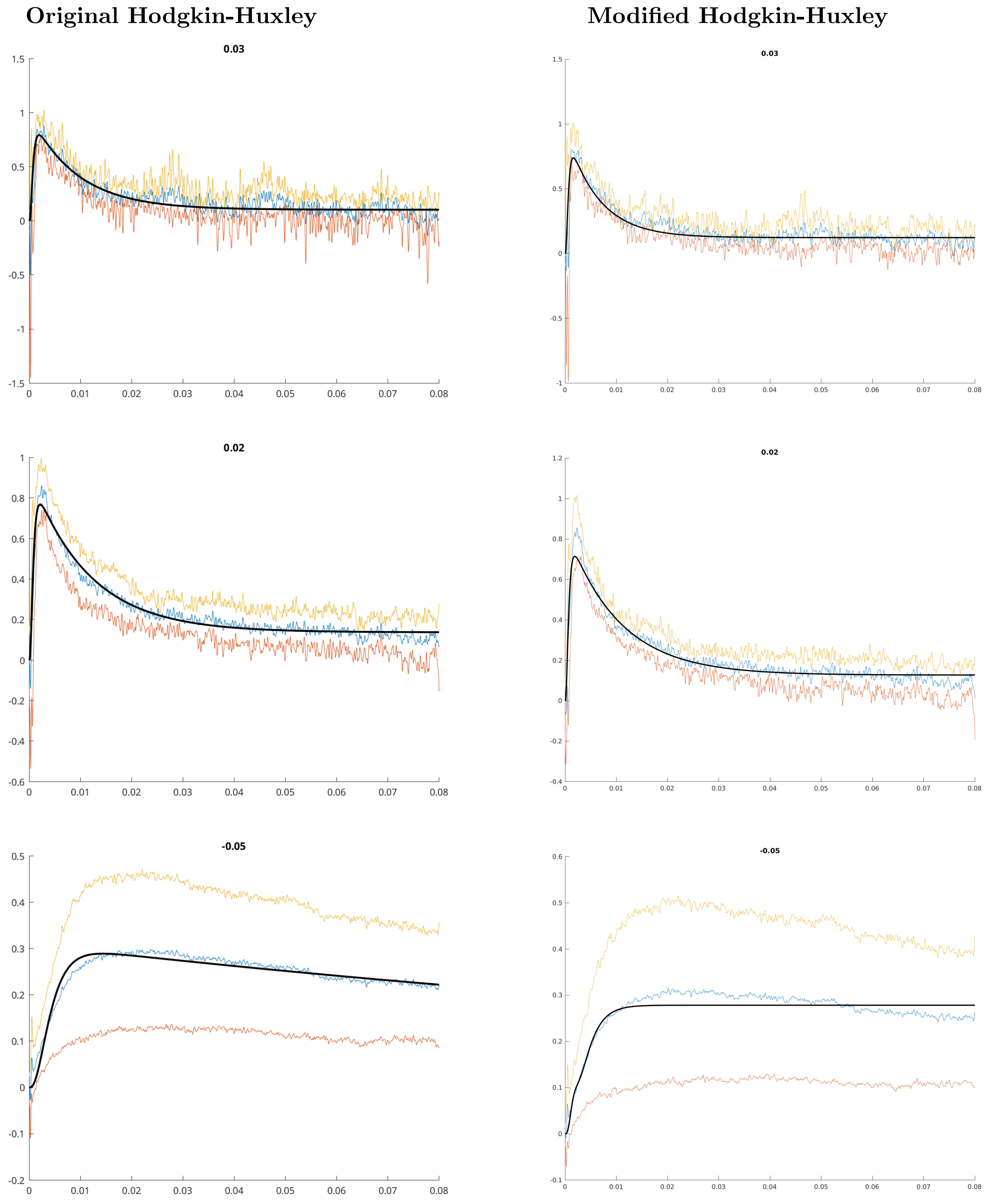

Supplementary Figure 10: Comparison of original and modified model for Nav1.9. The horizontal axis shows the time in $s$, the vertical axis the normalized sodium current. Panel titles show the clamping voltage in *volts*. Blue: average of measured cells, yellow and orange: average ± standard deviation.

between simulations and observations, i.e., we use the mapping $p \mapsto -p - 65mV$. This resulted in

$$\alpha_h(V_m[mV]) = \frac{0.01 \cdot (10.0 - (V_m + 65))}{\exp(1.0 - (0.1 \cdot (Vm + 65))) - 1.0}, \tag{12}$$

$$\beta_h(V_m[mV]) = 0.125 \cdot exp(-(V_m + 65)/80.0). \tag{13}$$

where $V_m$ is measured in *millivolts*. The gating variables for the sodium channels were parameterized based on our own data and are give by the function $\chi$ from equation (1), where $V_m$ is given in *volts*. The respective parameters of $\chi$ are provided in the following table.

| function | $a$ | $b$ | $c$ | $d$ | $f$ | $g$ | $q$ |
|---|---|---|---|---|---|---|---|
| $\alpha_{m,1.9}$ | 0.012809 | -0.19035 | 0 | 0.088411 | 0.23539 | -0.02587 | 0 |
| $\beta_{m,1.9}$ | -57.461 | -194.4368 | -0.036556 | -0.016775 | 0.073834 | 0.0081423 | 0 |
| $\alpha_{h,1.9}$ | 25.4063 | 344.8971 | 2.5413 | 47.625 | 0.037017 | -0.001453 | 0 |
| $\beta_{h,1.9}$ | 17.5241 | 298.0407 | 0.28349 | 0.037926 | 0.02984 | -0.0057412 | 0 |
| $\alpha_{m,1.8}$ | 109.3789 | -999.8963 | 0.0010931 | 0.0020029 | -0.048843 | -0.022244 | 0 |
| $\beta_{m,1.8}$ | -604.8165 | 0 | 0 | -0.51201 | 0 | 0.033752 | 0 |
| $\alpha_{h,1.8}$ | 206.0724 | -3.371 | 294.4345 | -328.0076 | -0.50016 | 3.6785 | 0 |
| $\beta_{h,1.8}$ | 17.4508 | 292.8407 | 0.058965 | 0.0057358 | -0.061377 | -0.030721 | 0 |
| $\gamma_{h,1.8}$ | -37.8993 | -15.3631 | -0.00061406 | -15.2606 | 0.18321 | -0.021847 | 0.0014255 |
| $\alpha_{m,1.7}$ | -88.3029 | -990.0132 | -0.0043814 | -0.19759 | 0.16128 | -0.034044 | 0 |
| $\beta_{m,1.7}$ | 11.6281 | -999.6937 | 0.0050351 | 0.096088 | 0.0040657 | 0.021937 | 0 |
| $\alpha_{h,1.7}$ | -63.4701 | 13.2776 | 0.054846 | -0.53512 | -0.025142 | -0.029311 | 0 |
| $\beta_{h,1.7}$ | -31.3657 | -165.6202 | -0.011608 | -0.016994 | 0.019332 | -0.013811 | 0 |
| $\gamma_{h,1.7}$ | 21242.0855 | 252919.9057 | 1 | 0 | 0 | 1 | 0 |
| $\alpha_{m,1.6}$ | 81.6161 | 972.7233 | 0.067819 | -0.12118 | -0.72364 | 1.083 | 0 |
| $\beta_{m,1.6}$ | 187.1987 | 0 | 0 | 1 | 0 | 0.014033 | 0 |
| $\alpha_{h,1.6}$ | 47.9342 | 601.0399 | 206.8624 | -224.0465 | -0.97327 | 11.251 | 0 |
| $\beta_{h,1.6}$ | -142.9141 | 999.3988 | -0.059411 | -0.26692 | 0.029413 | -0.012771 | 0 |
| $\gamma_{h,1.6}$ | -0.64764 | 76.3411 | -0.019961 | 147.6915 | 115.844 | -13.0107 | 31220.7915 |
| $\alpha_{m,1.5}$ | -69.7996 | -958.4105 | -0.005315 | 0.00060298 | -0.036619 | 0.0036496 | 0 |
| $\beta_{m,1.5}$ | 3.8215 | 193.355 | -0.0046047 | -0.0025651 | 0.058862 | 0.011492 | 0 |
| $\alpha_{h,1.5}$ | 46.9122 | 7.3413 | -0.0035953 | 0.32049 | -0.019881 | -0.022745 | 0 |
| $\beta_{h,1.5}$ | 401.2706 | 0 | 0.13312 | 0.086147 | 0.015675 | -0.013992 | 0 |
| $\gamma_{h,1.5}$ | 74.4554 | -974.757 | 0.005155 | 1.0409 | 0.039974 | -0.010188 | 15583.837 |
| $\alpha_{m,1.3}$ | -9.2779 | -87.9028 | -0.00054947 | -0.2961 | 0.13948 | -0.018614 | 0 |
| $\beta_{m,1.3}$ | -905.6435 | 0 | -0.23704 | -1.1001 | 0.024079 | 0.0090022 | 0 |
| $\alpha_{h,1.3}$ | -96.8351 | -140.6073 | 2.1407 | -4.0967 | -0.017261 | -0.069927 | 0 |
| $\beta_{h,1.3}$ | -36.6121 | -447.9485 | -0.017397 | -0.016165 | 0.010331 | -0.01491 | 0 |
| $\gamma_{h,1.3}$ | 1000 | -999.9749 | 0.0044704 | 5.4067e-07 | 0.062717 | 0.00477 | 26328.3751 |
| $\alpha_{m,1.2}$ | 88.5466 | 999.4955 | 0.0051046 | 0.33423 | 0.11721 | -0.021128 | 0 |
| $\beta_{m,1.2}$ | -73.5515 | 31.5788 | -0.023769 | -0.038974 | 0.031392 | 0.0036247 | 0 |
| $\alpha_{h,1.2}$ | -21.0803 | 192.7521 | 0.069443 | -0.26399 | -0.0245 | -0.029253 | 0 |
| $\beta_{h,1.2}$ | 811.1732 | 0 | 0.16495 | 0.26902 | 0.0088452 | -0.018089 | 0 |
| $\gamma_{h,1.2}$ | 999.992 | 0 | 0.014099 | 0.0023665 | 0.032441 | 0.0033897 | 23395.3259 |
| $\alpha_{m,1.1}$ | 91.6102 | 954.9235 | 0.0053553 | 0.019027 | 0.056761 | -0.023056 | 0 |
| $\beta_{m,1.1}$ | 2.76 | -972.6234 | 0.011493 | 0.0031945 | 0.04556 | 0.012647 | 0 |
| $\alpha_{h,1.1}$ | 47.453 | 53.5162 | -0.26038 | 0.7007 | -0.026895 | -0.054566 | 0 |
| $\beta_{h,1.1}$ | 997.1768 | 0 | 0.39278 | 0.38984 | 0.013832 | -0.015155 | 0 |
| $\gamma_{h,1.1}$ | -98.6873 | 0 | 0.0046454 | 2.2398 | -0.092769 | 0.015696 | 33569.9469 |

##### 4.2 Sodium Channel Abundance

The maximal conductance assigned to each sodium channel subtype was chosen to agree with the quantitative gene expression profiles from [2] (rounded to two digits). This resulted in the following choices for the CMi:

| | $g_{1.1}$ | $g_{1.2}$ | $g_{1.3}$ | $g_{1.5}$ | $g_{1.6}$ | $g_{1.7}$ | $g_{1.8}$ | $g_{1.9}$ |
|---|---|---|---|---|---|---|---|---|
| conductance $[mS/cm^2]$ | 0.22222 | 0.51852 | 0.07407 | 0.22222 | 1.25926 | 17.1111 | 7.03704 | 19.5556 |
| % of total Na-conductance | 0.48309 | 1.12721 | 0.16103 | 0.48309 | 2.73752 | 37.1981 | 15.2979 | 42.5121 |

The sodium conductances for the A$\delta$ fiber were set to:

| | $g_{1.1}$ | $g_{1.2}$ | $g_{1.3}$ | $g_{1.5}$ | $g_{1.6}$ | $g_{1.7}$ | $g_{1.8}$ | $g_{1.9}$ |
|---|---|---|---|---|---|---|---|---|
| conductance $[mS/cm^2]$ | 0.87342 | 0.87342 | 0.09705 | 0.29114 | 3.88186 | 19.6034 | 11.4515 | 8.92827 |
| % of total Na-conductance | 1.89873 | 1.89873 | 0.21097 | 0.63291 | 8.43882 | 42.616 | 24.8945 | 19.4093 |

### 5 Simulation

The ODE system was numerically solved using the function ode23s from Matlab R2024a, which is suitable for stiff systems. The initial condition of the nociceptor and A$\delta$ fiber for the simulations shown in Figs. 9, 10, S3, S4 and S5 corresponds to the resting state of the respective neuron with the parameters specified in the previous section. To quantify the contribution of a given isoform to the action potential, the conductance attributed to the respective isoform was set to zero. This results in a reduction of the total sodium conductance. The current injection of $30\mu A/cm^2$ or $30\mu A/cm^2$ persisted for the whole simulation.

For the simulations in Figs. 11A and 12A the conductance attributed to Nav1.7 was multiplied by 5. This results in an increase of the total sodium conductance. We numerically identified the resting state of both fiber types for the modified Nav1.7 conductance. These resting states were used as initial conditions for the respective simulations. The current injection of $1\mu A/cm^2$ persisted during the whole simulation. The simulations in Figs. 11B and 12B were obtained analogously. The fold increase of the Nav1.8 conductance was chosen such that the total sodium conductance for the simulation shown in Fig 11B was approximately equal to that in Fig 11A. Analogously, the total sodium conductances in Figs. 12A and 12B were approximately equal. For the CMi the Nav1.8 conductance was multiplied by 11, for the A$\delta$ fiber by 8. As initial condition we used the resting state of the respective fiber type for the modified Nav1.8 conductance.

For the simulations in Fig 11D-E and 12D-E the activation of Nav1.7 was shifted to more hyperpolarized potentials. This was achieved by replacing the voltage-dependent functions $\alpha_{m,1.7}(x)$ and $\beta_{m,1.7}(x)$ of the wildtype Nav1.7 by $\alpha_{m,1.7}(x+\Delta)$ and $\beta_{m,1.7}(x+\Delta)$, where $\Delta \in \{5mV, 10mV\}$. We numerically approximated the resting state for the modified Nav1.7 gating, which seemed to be unstable (up to numerical precision) for the A$\delta$ fiber in case of $\Delta = 10mV$. The respective resting states were used as initial conditions for the simulations. The current injections persisted during the whole simulated time interval.
