## supplemental figures for "Sodium channels expressed in nociceptors contribute distinctly to action potential subthreshold phase, upstroke and shoulder"

### Supplementary Figures

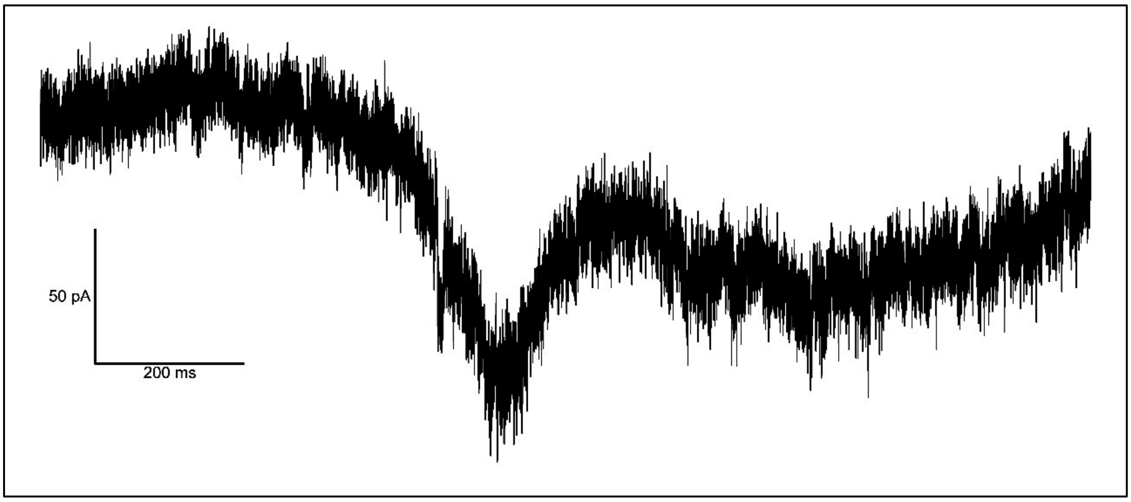


**Figure S1**: example trace of a $Na_v1.3$ ramp current elicited by a 0.1 mV/ms voltage ramp exhibiting two peaks



Parts of this study were published as a preprint: Köster et al. 2023. Biophysics of sodium channels during subthreshold depolarization *in vitro* and *in silico*. bioRxiv. doi.org/10.1101/2023.09.03.556095 (Preprint posted September 6, 2023)

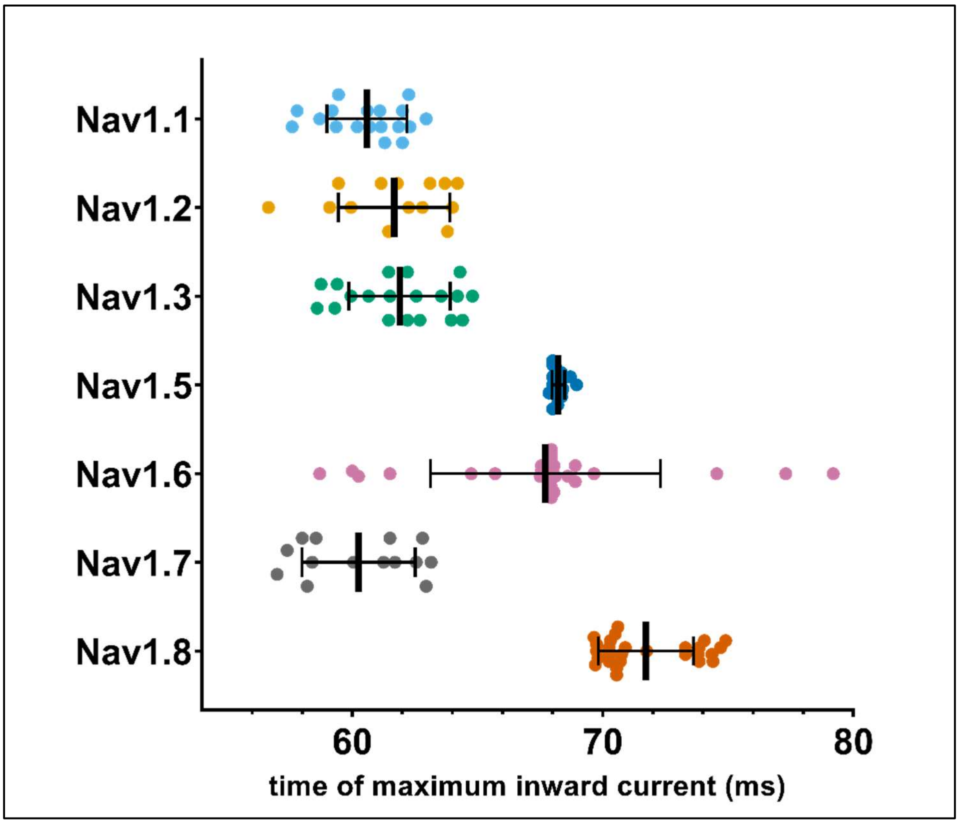


**Figure S2**: timepoint at which the maximum inward current during AP3 after exclusion of measurements impeded by transient current artifacts.



Parts of this study were published as a preprint: Köster et al. 2023. Biophysics of sodium channels during subthreshold depolarization *in vitro* and *in silico*. bioRxiv. doi.org/10.1101/2023.09.03.556095 (Preprint posted September 6, 2023)

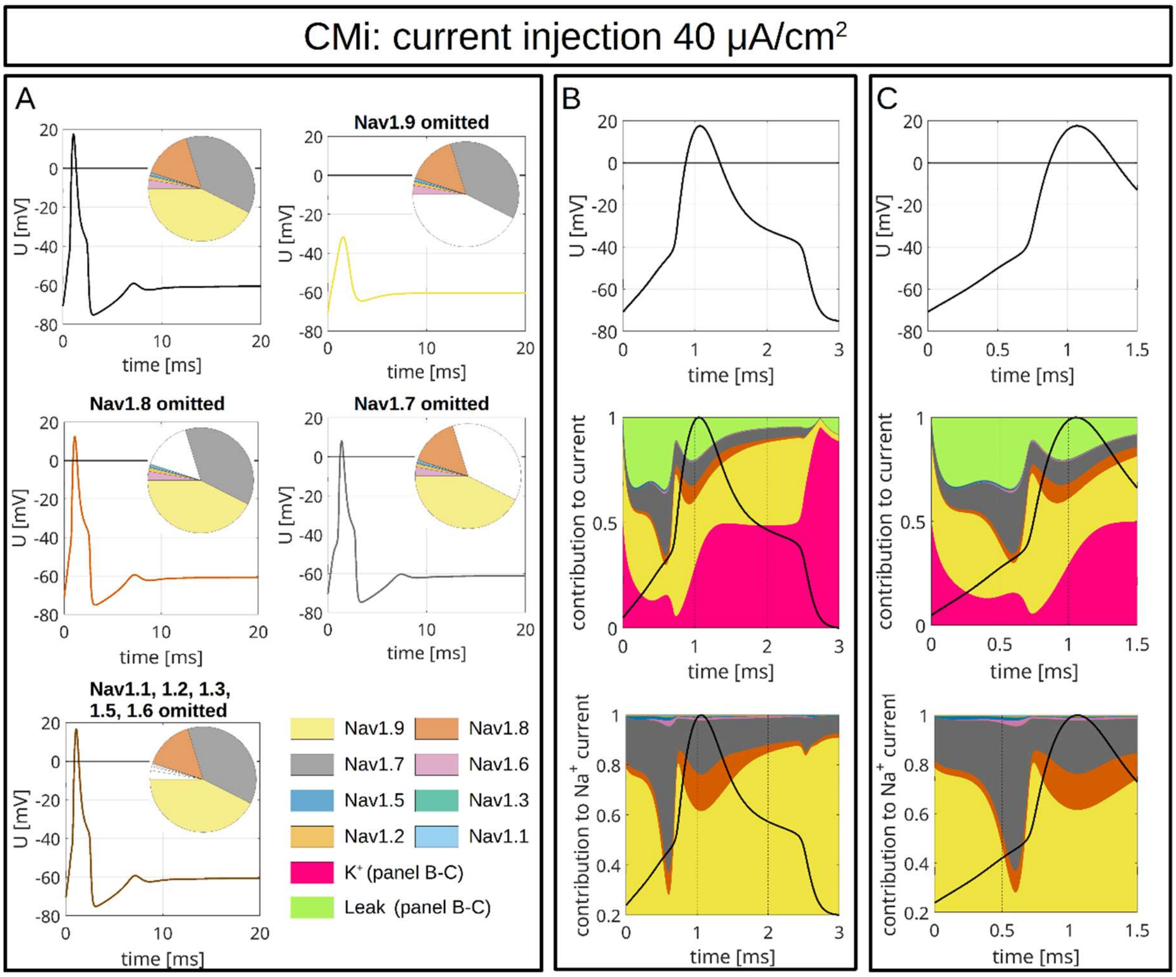


**Figure S3: Computer Simulations of APs in a CMi-fiber.** The pie charts indicate the abundance of different VGSC isoforms. APs are triggered by an injection of 40 µA/cm² which persists until the end of the simulation. *A:* The upper left panel shows the APs for the relative abundance of VGSC isoforms quantified in spatial gene expression. The initial condition of the simulation is the resting state of the CMi-fiber. The other panels show changes of the AP resulting from removal of $Na_v$1.9 (upper right), $Na_v$1.8 (middle left), $Na_v$1.7 (middle right) and $Na_v$1.1-1.3, 1.5 & 1.6 (bottom left). The VGSC isoforms are coded by the colors specified in the bottom right panel. B/C: VGSC isoform contributions to the simulated AP depicted in the upper left panel of A. The upper panels of B and C show the membrane potential during the initial 3 ms (B) and 1.5 ms (C) of the simulated AP. The middle panels show the relative contribution of the different sodium currents, the potassium current and the leak current over



Parts of this study were published as a preprint: Köster et al. 2023. Biophysics of sodium channels during subthreshold depolarization *in vitro* and *in silico*. bioRxiv. doi.org/10.1101/2023.09.03.556095 (Preprint posted September 6, 2023)

the course of the AP, plotted as stacked individual currents normalized to total current at each time point. The bottom panels show VGSC isoform contributions to the total sodium current, plotted as stacked individual VGSC isoform currents normalized to the total sodium current at each time point. The black line visualizes the shape of the AP. The color coding is identical to that in A.



Parts of this study were published as a preprint: Köster et al. 2023. Biophysics of sodium channels during subthreshold depolarization *in vitro* and *in silico*. bioRxiv. doi.org/10.1101/2023.09.03.556095 (Preprint posted September 6, 2023)

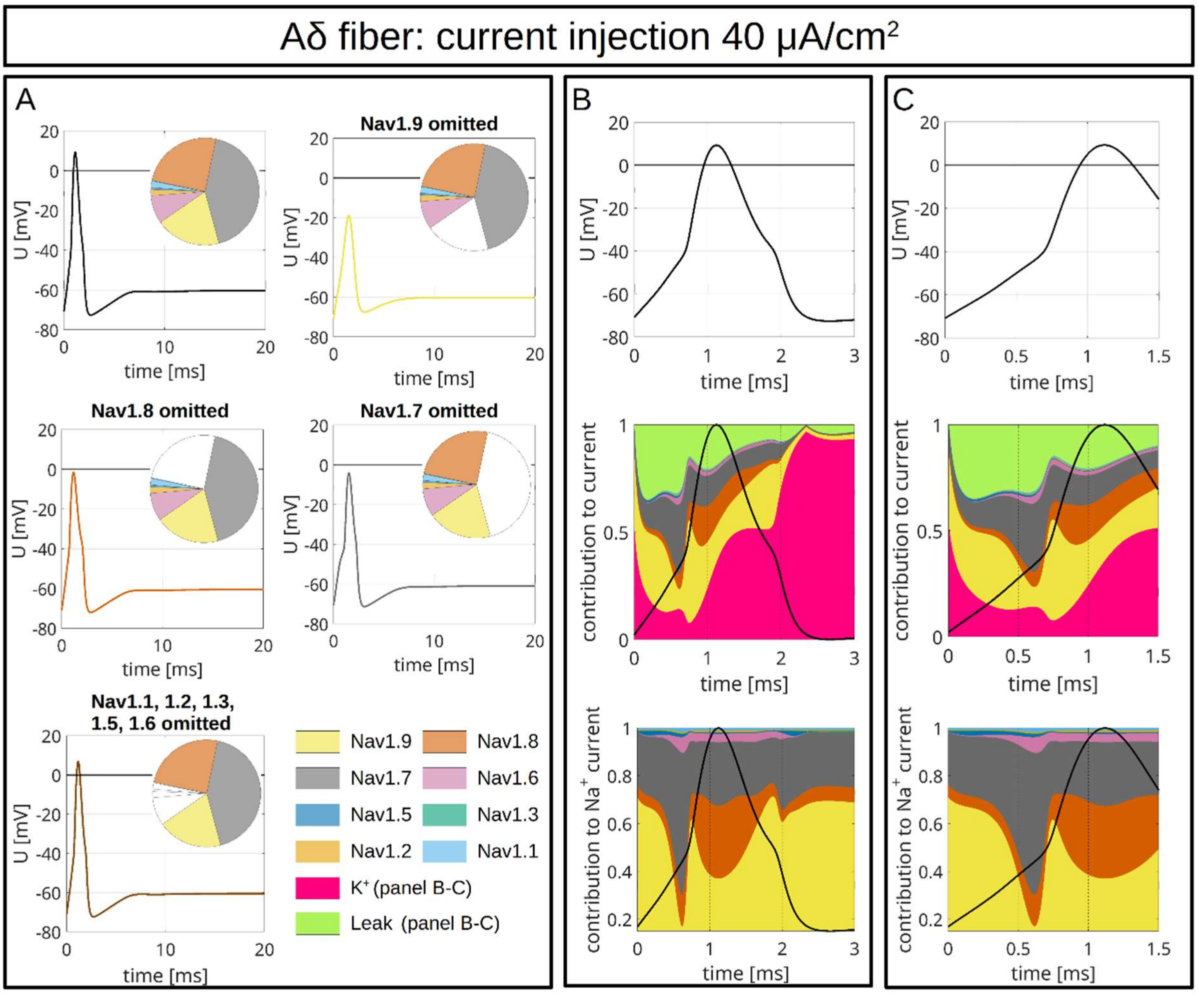


**Figure S4: Computer Simulations of AP in an Aδ-fiber.** The pie charts indicate the abundance of different VGSC isoforms. APs are triggered by an injection of 40 $\mu A/cm^2$ which persists until the end of the simulation. A: The upper left panel shows the APs for the relative abundance of VGSC isoforms quantified in spatial gene expression. The initial condition of the simulation is the resting state of the Aδ-fiber. The other panels show changes of the AP resulting from removal of $Na_v1.9$ (upper right), $Na_v1.8$ (middle left), $Na_v1.7$ (middle right) and $Na_v1.1$-1.3, 1.5 & 1.6 (bottom left). The VGSC isoforms are coded by the colors specified in the bottom right panel. B/C: VGSC isoform contributions to the simulated AP depicted in the upper left panel of A. The upper panels of B and C show the membrane potential during the initial 3 ms (B) and 1.5 ms (C) of the simulated AP. The middle panels show the relative contribution of the different sodium currents, the potassium current and the leak current over



Parts of this study were published as a preprint: Köster et al. 2023. Biophysics of sodium channels during subthreshold depolarization *in vitro* and *in silico*. bioRxiv. doi.org/10.1101/2023.09.03.556095 (Preprint posted September 6, 2023)

the course of the AP, plotted as stacked individual currents normalized to total current at each time point. The bottom panels show VGSC isoform contributions to the total sodium current, plotted as stacked individual VGSC isoform currents normalized to the total sodium current at each time point. The black line visualizes the shape of the AP. The color coding is identical to that in A.



Parts of this study were published as a preprint: Köster et al. 2023. Biophysics of sodium channels during subthreshold depolarization *in vitro* and *in silico*. bioRxiv. doi.org/10.1101/2023.09.03.556095 (Preprint posted September 6, 2023)

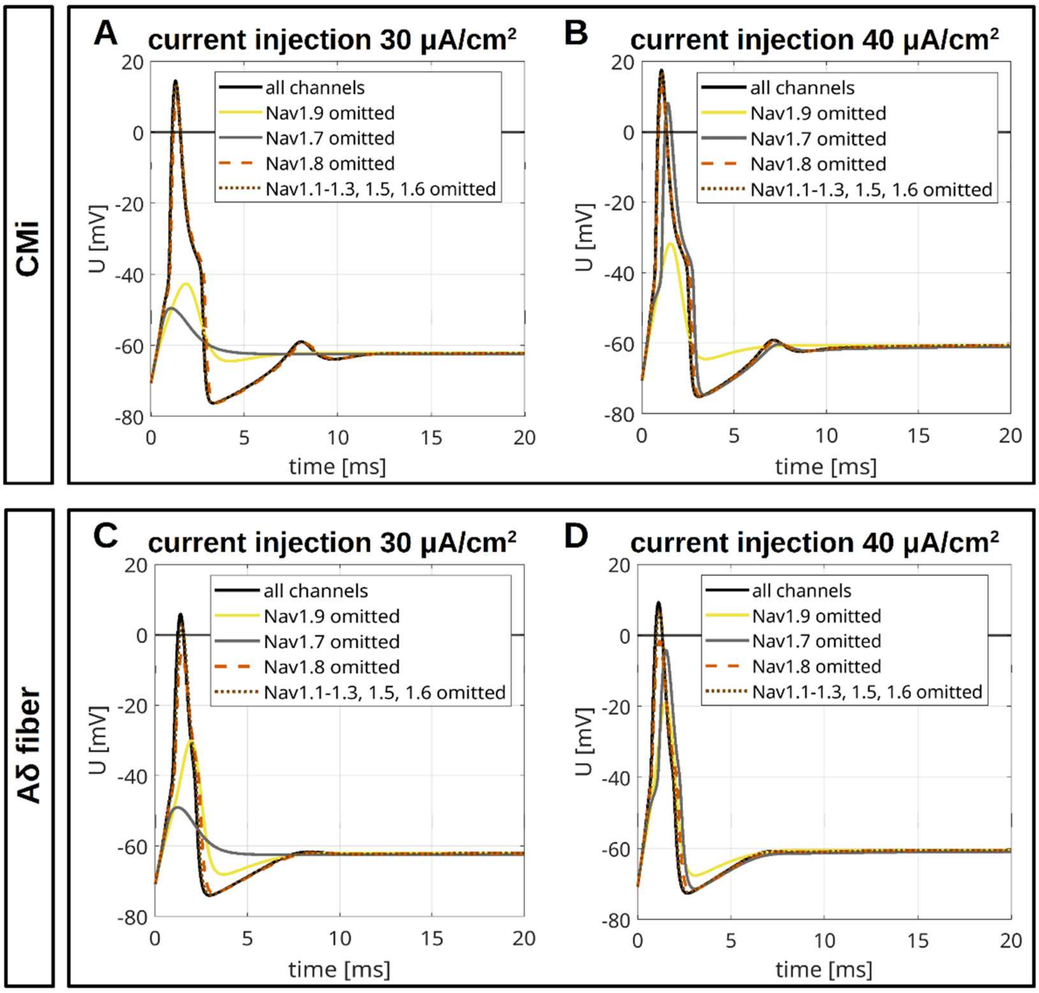


**Figure S5: Overlay of Simulated APs in a CMi- and in an Aδ-fiber.** A/B: AP for the CMi-fiber with the relative abundance of VGSC isoforms quantified in spatial gene expression (black, isoform abundances as in Fig. 9A, upper left panel) and changes of the AP resulting from removal of $Na_v1.9$ (yellow), $Na_v1.8$ (orange), $Na_v1.7$ (gray) and $Na_v1.1$-1.3, 1.5 & 1.6 (brown). The current injections of 30 μA/cm² (A) and 40 μA/cm² (B) persist until the end of the simulation. C/D: AP for the Aδ-fiber with the relative abundance of VGSC isoforms quantified in spatial gene expression (black, isoform abundances as in Fig. 10A, upper left panel) and changes of the AP resulting from removal of $Na_v1.9$ (yellow), $Na_v1.8$ (orange), $Na_v1.7$ (gray) and $Na_v1.1$-1.3, 1.5 & 1.6 (brown). The current injections of 30 μA/cm² (C) and 40 μA/cm² (D) persist until the end of the simulation.



Parts of this study were published as a preprint: Köster et al. 2023. Biophysics of sodium channels during subthreshold depolarization *in vitro* and *in silico*. bioRxiv. doi.org/10.1101/2023.09.03.556095 (Preprint posted September 6, 2023)

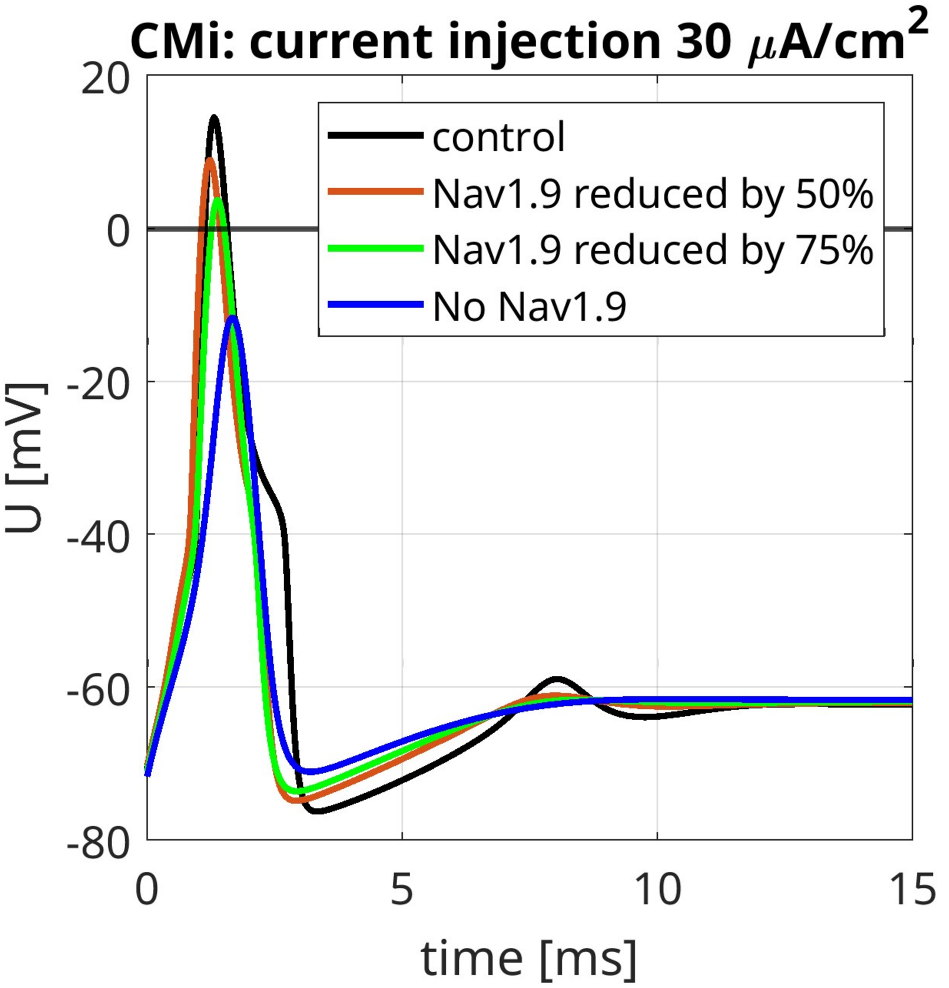


**Figure S6: Simulated Contribution of $Na_v$1.9 to APs in a CMi-fiber.** The black line (control) shows the AP in a CMi-fiber with the relative abundance of VGSC isoforms quantified in spatial gene expression (isoform abundances as in Fig. 9A, upper left panel). The maximal conductance of $Na_v$1.9 is stepwise reduced to 50% (red line), 25% (green line) and 0% (blue line) compared to the control scenario. The maximal conductances of the other isoforms are increased proportionally to their expression, such that the total maximal sodium conductance remains unchanged. This is different to Fig. 9A, where the total maximal sodium conductance decreased upon removal of a VGSC isoform. The current injections of 30 μA/cm² persist until the end of the simulation.



Parts of this study were published as a preprint: Köster et al. 2023. Biophysics of sodium channels during subthreshold depolarization *in vitro* and *in silico*. bioRxiv. doi.org/10.1101/2023.09.03.556095 (Preprint posted September 6, 2023)

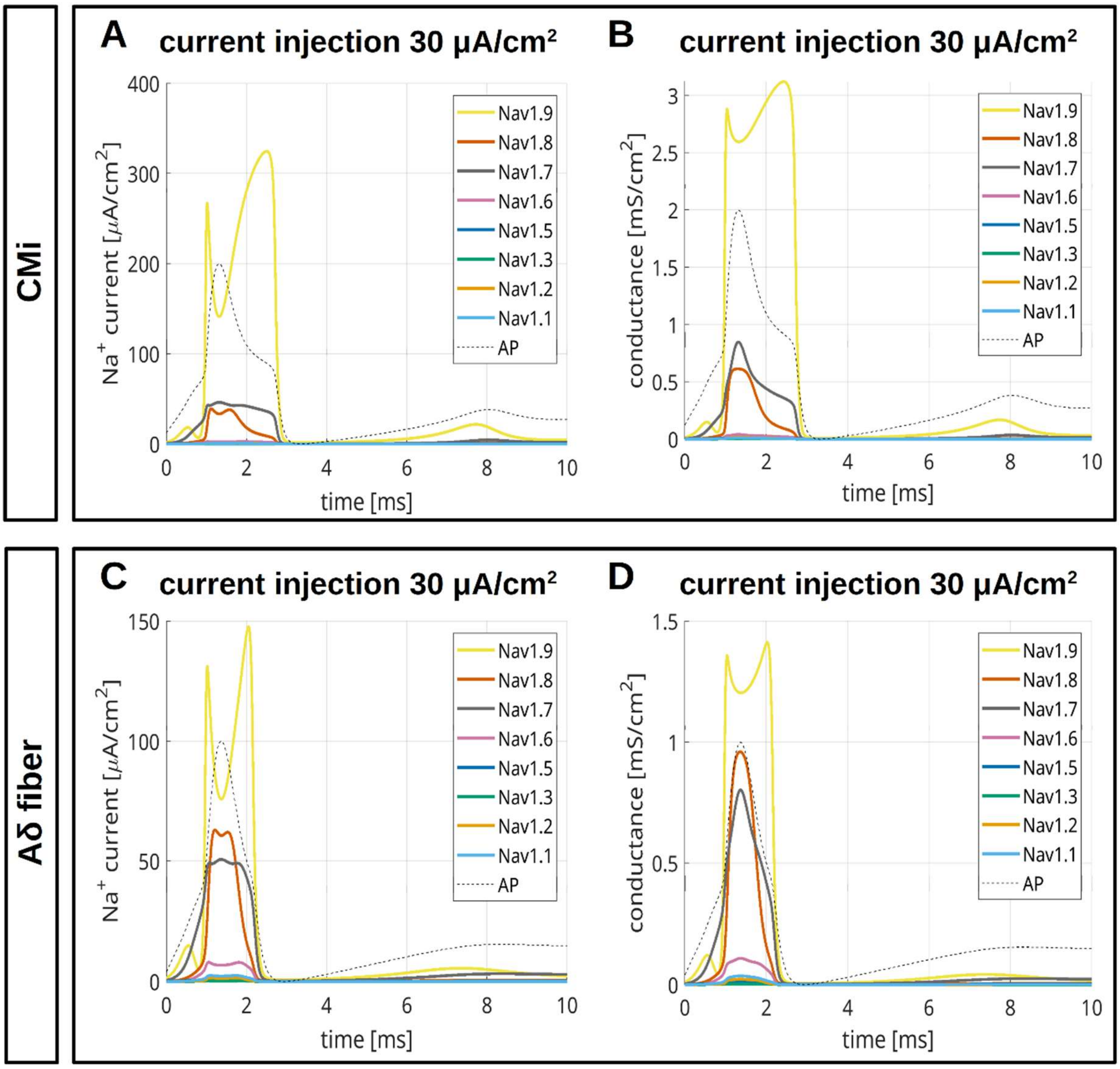


**Figure S7: Simulated Sodium Currents and Conductances in a CMi- and Aδ-fiber.** A: Absolute sodium currents over the course of the simulated AP of the CMi-fiber depicted in Fig. 9A, upper left panel. B: Conductances of the different VGSC isoforms over the course of the simulated AP of the CMi-fiber depicted in Fig. 9A, upper left panel. C: Absolute sodium currents over the course of the simulated AP of the Aδ-fiber depicted in Fig. 10A, upper left panel. D: Conductances of the different VGSC isoforms over the course of the simulated AP of the Aδ-fiber depicted in Fig. 10A, upper left panel. The current injection of 30 μA/cm² persist until the end of the simulation. The dotted black line visualizes the time course of the membrane potential during the AP.



Parts of this study were published as a preprint: Köster et al. 2023. Biophysics of sodium channels during subthreshold depolarization *in vitro* and *in silico*. bioRxiv. doi.org/10.1101/2023.09.03.556095 (Preprint posted September 6, 2023)
