## supplemental tables for "Sodium channels expressed in nociceptors contribute distinctly to action potential subthreshold phase, upstroke and shoulder"

**Supplemental Table 1: Overview of all used cell lines and their culture media and supplements**

| cell line | cell type | culture media and supplements |
|---|---|---|
| HEK293T (purchased by Sigma-Aldrich, St. Louis, Missouri, USA) | human embryonic kidney cells | Dulbecco's modified Eagle medium (DMEM) with L-Glutamine (Gibco, Thermo Fisher Scientific, Waltham, Massachusetts, USA) + 10% fetal bovine serum (FBS; Biochrom AG, Berlin, Germany) |
| ND7/23 (purchased by Sigma-Aldrich, St. Louis, Missouri, USA) | mouse neuroblastoma (N18 tg 2) x rat dorsal root ganglion neuron hybrid cells | DMEM with high glucose level (4,5 g/l) and L-Glutamine (Gibco, Thermo Fisher Scientific, Waltham, Massachusetts, USA) + 10% FBS |
| HEK293 r$Na_v$1.3 (Cummins et al., 2001) | human embryonic kidney cells, stably transfected with pcDNA3-rNav1.3 plasmid | DMEM with high glucose level (4,5 g/L) and L-Glutamine + 10% FBS + 0,5 mg/ml Geneticin disulphate (G418; Carl Roth GmbH + Co. KG, Karlsruhe, Germany) |
| HEK293 h$Na_v$1.5 (Eberhardt et al., 2015) | human embryonic kidney cells, stably transfected with pTracer-hNav1.5 plasmid | Dulbecco's modified Eagle medium F-12 (DMEM/F-12) with L-Glutamine (Gibco, Thermo Fisher Scientific, Waltham, Massachusetts, USA) + 10% FBS + 100 µg/ml Zeocin (Invivogen, San Diego, California, USA) |
| HEK293 m$Na_v$1.6 (Herzog et al., 2003 and Laezza et al., 2009) | human embryonic kidney cells stably transfected with pCIN5-mNav1.6 plasmid | DMEM/F-12 with L-Glutamine + 10% FBS + 0,5 mg/ml G418 |
| HEK293 h$Na_v$1.7 (Körner et al., 2018; vector by Klugbauer et al., 1995) | human embryonic kidney cells stably transfected with pcDNA3-hNav1.7 plasmid | DMEM/F-12 with L-Glutamine + 10% FBS + 0,5 mg/ml G418 |

**Supplemental Table 2: Duration of command voltage ramps**

| Ramp rate [mV/ms] | Duration for 140mV [ms] |
|---:|---:|
| 0,1 | 1400 |
| 0,2 | 700 |
| 0,4 | 350 |
| 0,6 | 233,3333333 |
| 1,2 | 116,6666667 |
| 2 | 70 |
| 4 | 35 |
| 6 | 23,33333333 |

**Supplemental Table 3: Multiple comparisons of V50 values of voltage dependency of activation and fast inactivation measurement**

| Activation | Nav1.1 | | Nav1.2 | | Nav1.3 | | Nav1.5 | | Nav1.6 | | Nav1.7 | | Nav1.8 | | Inactivation |
|---|---|---|---|---|---|---|---|---|---|---|---|---|---|---|---|
| Nav1.1 | | | ns | >0,9999 | ns | >0,9999 | **** | <0,0001 | ns | >0,9999 | *** | 0,0005 | ns | 0,0538 | Nav1.1 |
| Nav1.2 | ns | >0,9999 | | | ns | >0,9999 | **** | <0,0001 | ns | >0,9999 | ns | 0,1267 | *** | 0,0002 | Nav1.2 |
| Nav1.3 | ns | 0,9995 | ns | >0,9999 | | | **** | <0,0001 | ns | >0,9999 | * | 0,0178 | ** | 0,0021 | Nav1.3 |
| Nav1.5 | **** | <0,0001 | **** | <0,0001 | **** | <0,0001 | | | **** | <0,0001 | ns | 0,0616 | **** | <0,0001 | Nav1.5 |
| Nav1.6 | ** | 0,0038 | ** | 0,0081 | * | 0,0164 | **** | <0,0001 | | | ns | 0,1855 | **** | <0,0001 | Nav1.6 |
| Nav1.7 | * | 0,0127 | * | 0,025 | * | 0,0468 | **** | <0,0001 | ns | 0,9998 | | | **** | <0,0001 | Nav1.7 |
| Nav1.8 | **** | <0,0001 | **** | <0,0001 | **** | <0,0001 | **** | <0,0001 | **** | <0,0001 | **** | <0,0001 | | | Nav1.8 |
| | Nav1.1 | | Nav1.2 | | Nav1.3 | | Nav1.5 | | Nav1.6 | | Nav1.7 | | Nav1.8 | | |

Activation data: adjusted P value of Tukey's multiple comparisons test

Inactivation data: adjusted P value of Dunn's multiple comparisons test

**Supplemental Table 4: Multiple comparisons of slope values (k) of voltage dependency of activation and fast inactivation measurement**

| | | Nav1.1 | | Nav1.2 | | Nav1.3 | | Nav1.5 | | Nav1.6 | | Nav1.7 | | Nav1.8 | | | |
|---|---|---|---|---|---|---|---|---|---|---|---|---|---|---|---|---|---|
| | Nav1.1 | | | ns | >0,9999 | ** | 0,0015 | ns | >0,9999 | ** | 0,0039 | * | 0,0332 | **** | <0,0001 | Nav1.1 | |
| | Nav1.2 | ns | >0,9999 | | | * | 0,0131 | ns | >0,9999 | * | 0,03 | ns | 0,189 | **** | <0,0001 | Nav1.2 | |
| | Nav1.3 | ns | >0,9999 | ns | >0,9999 | | | ** | 0,0036 | ns | >0,9999 | ns | >0,9999 | ** | 0,0028 | Nav1.3 | |
| Activation | Nav1.5 | ns | >0,9999 | ns | >0,9999 | ns | >0,9999 | | | ** | 0,009 | ns | 0,0676 | **** | <0,0001 | Nav1.5 | Inactivation |
| | Nav1.6 | ** | 0,0063 | ns | 0,124 | ns | 0,9217 | ns | >0,9999 | | | ns | >0,9999 | ** | 0,0011 | Nav1.6 | |
| | Nav1.7 | ns | >0,9999 | ns | >0,9999 | ns | >0,9999 | ns | >0,9999 | ns | 0,3398 | | | **** | <0,0001 | Nav1.7 | |
| | Nav1.8 | **** | <0,0001 | **** | <0,0001 | **** | <0,0001 | **** | <0,0001 | **** | <0,0001 | **** | <0,0001 | | | Nav1.8 | |
| | | Nav1.1 | | Nav1.2 | | Nav1.3 | | Nav1.5 | | Nav1.6 | | Nav1.7 | | Nav1.8 | | | |

Data: adjusted P value of Dunn's multiple comparisons test

**Supplemental Table 5: Multiple comparisons of window current AUC and intersection voltage of activation and SSFI Boltzmann fit curves**

| window current AUC | Nav1.1 | | Nav1.2 | | Nav1.3 | | Nav1.5 | | Nav1.6 | | Nav1.7 | | Nav1.8 | | intersection voltages of act/SSFI curves |
|---|---|---|---|---|---|---|---|---|---|---|---|---|---|---|---|
| Nav1.1 | | | ns | 0,9045 | ns | 0,9297 | **** | <0,0001 | ns | 0,9999 | ** | 0,0015 | **** | <0,0001 | Nav1.1 |
| Nav1.2 | ns | >0,9999 | | | ns | 0,2682 | **** | <0,0001 | ns | 0,9822 | ns | 0,0676 | **** | <0,0001 | Nav1.2 |
| Nav1.3 | ns | >0,9999 | ns | >0,9999 | | | **** | <0,0001 | ns | 0,7801 | **** | <0,0001 | **** | <0,0001 | Nav1.3 |
| Nav1.5 | ns | >0,9999 | ns | >0,9999 | ns | >0,9999 | | | **** | <0,0001 | **** | <0,0001 | **** | <0,0001 | Nav1.5 |
| Nav1.6 | ns | >0,9999 | * | 0,0489 | * | 0,0493 | ns | >0,9999 | | | ** | 0,0053 | **** | <0,0001 | Nav1.6 |
| Nav1.7 | ns | >0,9999 | ns | >0,9999 | ns | >0,9999 | ns | >0,9999 | ns | >0,9999 | | | **** | <0,0001 | Nav1.7 |
| Nav1.8 | ** | 0,0051 | **** | <0,0001 | **** | <0,0001 | ** | 0,0092 | ns | 0,6778 | * | 0,0175 | | | Nav1.8 |
| | Nav1.1 | | Nav1.2 | | Nav1.3 | | Nav1.5 | | Nav1.6 | | Nav1.7 | | Nav1.8 | | |

window current AUC data: adjusted P value of Dunn's multiple comparisons test
act/SSFI intersection data: adjusted P value of Tukey's multiple comparisons test

**Supplemental Table 6: Multiple comparisons of fraction of channels remaining open after SSFI**

fraction of channels remaining open after SSFI

| | Nav1.1 | | Nav1.2 | | Nav1.3 | | Nav1.5 | | Nav1.6 | | Nav1.7 | | Nav1.8 |
|---|---|---|---|---|---|---|---|---|---|---|---|---|---|
| **Nav1.1** | | | | | | | | | | | | | |
| **Nav1.2** | ns | >0,9999 | | | | | | | | | | | |
| **Nav1.3** | ns | >0,9999 | ns | >0,9999 | | | | | | | | | |
| **Nav1.5** | ns | >0,9999 | ns | 0,2983 | * | 0,0126 | | | | | | | |
| **Nav1.6** | ns | >0,9999 | ns | 0,3377 | * | 0,0148 | ns | >0,9999 | | | | | |
| **Nav1.7** | ns | >0,9999 | ns | >0,9999 | ns | 0,0887 | ns | >0,9999 | ns | >0,9999 | | | |
| **Nav1.8** | ns | 0,138 | ** | 0,0083 | *** | 0,0001 | ns | >0,9999 | ns | >0,9999 | ns | >0,9999 | |

Data: adjusted P value of Dunn's multiple comparisons test

**Supplemental Table 7: Descriptive statistics of parameters gathered from ramp current measurements**

Data is mean ± SD, n = sample size

| | ramp rate (mV/ms) | Imax(ramp)/ Imax(act) | U of Imax(ramp) (mV) | n | AUC (normalized) | n |
|---|---|---|---|---|---|---|
| Nav1.1 | **0,1** | 0,029 ± 0,033 | -47,6 ± 9 | 26 | 0,0007 ± 0,0036 | 24 |
| | **0,2** | 0,028 ± 0,025 | -48,1 ± 9,8 | 26 | 0,0045 ± 0,0049 | 24 |
| | **0,4** | 0,035 ± 0,028 | -47,5 ± 7,3 | 26 | 0,0049 ± 0,0041 | 24 |
| | **0,6** | 0,045 ± 0,03 | -49,6 ± 4,8 | 27 | 0,0058 ± 0,0042 | 24 |
| | **1,2** | 0,08 ± 0,033 | -46 ± 4,3 | 27 | 0,0112 ± 0,0044 | 24 |
| | **2** | 0,131 ± 0,041 | -44,3 ± 4,4 | 27 | 0,0173 ± 0,0056 | 24 |
| | **4** | 0,248 ± 0,058 | -40,4 ± 4,6 | 27 | 0,0324 ± 0,0094 | 24 |
| | **6** | 0,343 ± 0,067 | -37,7 ± 4,7 | 27 | 0,0449 ± 0,01 | 24 |
| Nav1.2 | **0,1** | 0,027 ± 0,029 | -51,4 ± 9,2 | 26 | 0,0016 ± 0,0035 | 20 |
| | **0,2** | 0,028 ± 0,026 | -48,9 ± 7,8 | 26 | 0,0024 ± 0,0039 | 21 |
| | **0,4** | 0,034 ± 0,028 | -47,4 ± 6 | 27 | 0,004 ± 0,0033 | 22 |
| | **0,6** | 0,041 ± 0,026 | -46,9 ± 5,6 | 27 | 0,0049 ± 0,0047 | 22 |
| | **1,2** | 0,073 ± 0,035 | -44,6 ± 5,8 | 27 | 0,0103 ± 0,0074 | 22 |
| | **2** | 0,12 ± 0,048 | -43,2 ± 6,1 | 27 | 0,0173 ± 0,011 | 22 |
| | **4** | 0,229 ± 0,077 | -39,3 ± 5,5 | 27 | 0,0322 ± 0,0187 | 22 |
| | **6** | 0,323 ± 0,094 | -37 ± 5,8 | 27 | 0,0437 ± 0,0216 | 22 |
| Nav1.3 | **0,1** | 0,026 ± 0,02 | -47,5 ± 6,8 | 26 | 0,0022 ± 0,0046 | 23 |
| | **0,2** | 0,042 ± 0,019 | -46,8 ± 4,7 | 26 | 0,0053 ± 0,0045 | 23 |
| | **0,4** | 0,078 ± 0,019 | -45,1 ± 4,5 | 27 | 0,0098 ± 0,0034 | 24 |
| | **0,6** | 0,117 ± 0,031 | -43,7 ± 4,3 | 27 | 0,0151 ± 0,0055 | 25 |
| | **1,2** | 0,211 ± 0,052 | -40,9 ± 4,3 | 27 | 0,0268 ± 0,0067 | 25 |
| | **2** | 0,307 ± 0,072 | -39 ± 4,3 | 27 | 0,0381 ± 0,0093 | 25 |
| | **4** | 0,46 ± 0,095 | -35,7 ± 4,9 | 27 | 0,0599 ± 0,0121 | 25 |
| | **6** | 0,552 ± 0,104 | -33,6 ± 4,8 | 27 | 0,0718 ± 0,0153 | 25 |
| Nav1.5 | **0,1** | 0,017 ± 0,015 | -73,8 ± 7,6 | 18 | 0,0001 ± 0,0018 | 13 |
| | **0,2** | 0,025 ± 0,022 | -71,9 ± 10 | 22 | 0,0038 ± 0,0034 | 15 |
| | **0,4** | 0,032 ± 0,02 | -71,7 ± 4,4 | 26 | 0,0051 ± 0,004 | 15 |
| | **0,6** | 0,046 ± 0,026 | -70,5 ± 4,6 | 27 | 0,008 ± 0,005 | 16 |
| | **1,2** | 0,092 ± 0,047 | -67 ± 4,6 | 27 | 0,0194 ± 0,0098 | 16 |
| | **2** | 0,149 ± 0,07 | -64,5 ± 4 | 27 | 0,0304 ± 0,0115 | 16 |
| | **4** | 0,265 ± 0,109 | -60,2 ± 4,3 | 27 | 0,0567 ± 0,0184 | 16 |
| | **6** | 0,344 ± 0,138 | -57,4 ± 3,8 | 27 | 0,0744 ± 0,0241 | 17 |
| Nav1.6 | **0,1** | 0,032 ± 0,018 | -58,2 ± 11,5 | 27 | -0,0017 ± 0,0073 | 23 |
| | **0,2** | 0,036 ± 0,018 | -57,7 ± 11,3 | 26 | 0,0041 ± 0,0051 | 22 |
| | **0,4** | 0,039 ± 0,016 | -53,8 ± 6,9 | 27 | 0,0068 ± 0,0046 | 23 |
| | **0,6** | 0,051 ± 0,023 | -53,1 ± 6 | 27 | 0,008 ± 0,0045 | 23 |
| | **1,2** | 0,093 ± 0,032 | -47 ± 5,4 | 27 | 0,0172 ± 0,0075 | 21 |
| | **2** | 0,155 ± 0,046 | -44,2 ± 5,8 | 27 | 0,025 ± 0,008 | 23 |
| | **4** | 0,308 ± 0,069 | -40,6 ± 5,6 | 27 | 0,0464 ± 0,0151 | 22 |
| | **6** | 0,433 ± 0,085 | -37,4 ± 6,3 | 27 | 0,0654 ± 0,0147 | 22 |
| Nav1.7 | **0,1** | 0,035 ± 0,029 | -56,2 ± 8 | 27 | 0,0006 ± 0,0033 | 20 |
| | **0,2** | 0,037 ± 0,027 | -55,7 ± 6,2 | 27 | 0,0035 ± 0,0046 | 20 |
| | **0,4** | 0,05 ± 0,03 | -51,7 ± 8 | 27 | 0,0063 ± 0,004 | 20 |
| | **0,6** | 0,067 ± 0,035 | -51,7 ± 6,8 | 27 | 0,0083 ± 0,0036 | 21 |
| | **1,2** | 0,126 ± 0,046 | -48,1 ± 6,7 | 27 | 0,0183 ± 0,006 | 21 |
| | **2** | 0,201 ± 0,057 | -45,7 ± 6,3 | 27 | 0,0279 ± 0,0085 | 21 |
| | **4** | 0,36 ± 0,08 | -41,9 ± 6,6 | 27 | 0,0498 ± 0,0159 | 20 |
| | **6** | 0,48 ± 0,089 | -40 ± 6,9 | 27 | 0,0653 ± 0,0184 | 20 |
| Nav1.8 | **0,1** | 0,117 ± 0,1 | -0,3 ± 8,7 | 24 | 0,0247 ± 0,0265 | 24 |
| | **0,2** | 0,123 ± 0,101 | -0,4 ± 8,6 | 26 | 0,0306 ± 0,0335 | 26 |
| | **0,4** | 0,133 ± 0,093 | -2,6 ± 10,7 | 27 | 0,0313 ± 0,0299 | 26 |
| | **0,6** | 0,158 ± 0,11 | -3,5 ± 9,6 | 27 | 0,0365 ± 0,0358 | 26 |
| | **1,2** | 0,209 ± 0,087 | -5,9 ± 8 | 27 | 0,0443 ± 0,0367 | 27 |
| | **2** | 0,278 ± 0,083 | -6,3 ± 5,6 | 27 | 0,0514 ± 0,0351 | 27 |
| | **4** | 0,417 ± 0,095 | -4,1 ± 5,3 | 27 | 0,0728 ± 0,0304 | 27 |
| | **6** | 0,512 ± 0,109 | -1,8 ± 6,3 | 27 | 0,0882 ± 0,0308 | 27 |

**Supplemental Table 8: Multiple comparisons of normalized maximum inward current values from ramp current measurements**

| ramp rate (mV/ms) | VGSC isoform | Nav1.1 | | Nav1.2 | | Nav1.3 | | Nav1.5 | | Nav1.6 | | Nav1.7 | | Nav1.8 | | VGSC isoform | ramp rate (mV/ms) |
|---|---|---|---|---|---|---|---|---|---|---|---|---|---|---|---|---|---|
| | Nav1.1 | | | ns | >0,9999 | ns | 0,9832 | ns | >0,9999 | ns | 0,9993 | ns | 0,9984 | **** | <0,0001 | Nav1.1 | |
| | Nav1.2 | ns | >0,9999 | | | ns | 0,9799 | ns | >0,9999 | ns | 0,999 | ns | 0,9979 | **** | <0,0001 | Nav1.2 | |
| | Nav1.3 | ns | >0,9999 | ns | >0,9999 | | | ns | 0,9614 | ns | 0,9998 | ns | >0,9999 | **** | <0,0001 | Nav1.3 | |
| 0,1 | Nav1.5 | ns | 0,9968 | ns | 0,9989 | ns | 0,9991 | | | ns | 0,996 | ns | 0,9932 | **** | <0,0001 | Nav1.5 | 0,2 |
| | Nav1.6 | ns | >0,9999 | ns | >0,9999 | ns | >0,9999 | ns | 0,9877 | | | ns | >0,9999 | **** | <0,0001 | Nav1.6 | |
| | Nav1.7 | ns | 0,9998 | ns | 0,999 | ns | 0,9987 | ns | 0,9649 | ns | >0,9999 | | | **** | <0,0001 | Nav1.7 | |
| | Nav1.8 | **** | <0,0001 | **** | <0,0001 | **** | <0,0001 | **** | <0,0001 | **** | <0,0001 | **** | <0,0001 | | | Nav1.8 | |
| | Nav1.1 | | | ns | >0,9999 | *** | 0,0005 | ns | >0,9999 | ns | 0,9998 | ns | 0,8608 | **** | <0,0001 | Nav1.1 | |
| | Nav1.2 | ns | >0,9999 | | | *** | 0,0002 | ns | >0,9999 | ns | 0,9979 | ns | 0,7602 | **** | <0,0001 | Nav1.2 | |
| | Nav1.3 | ns | 0,17 | ns | 0,1461 | | | *** | 0,0008 | ** | 0,0024 | ns | 0,0538 | ns | 0,1962 | Nav1.3 | |
| 0,4 | Nav1.5 | ns | >0,9999 | ns | >0,9999 | ns | 0,1165 | | | ns | >0,9999 | ns | 0,8985 | **** | <0,0001 | Nav1.5 | 0,6 |
| | Nav1.6 | ns | >0,9999 | ns | >0,9999 | ns | 0,2728 | ns | 0,9996 | | | ns | 0,9695 | **** | <0,0001 | Nav1.6 | |
| | Nav1.7 | ns | 0,9774 | ns | 0,9699 | ns | 0,6681 | ns | 0,9477 | ns | 0,9961 | | | **** | <0,0001 | Nav1.7 | |
| | Nav1.8 | **** | <0,0001 | **** | <0,0001 | * | 0,0207 | **** | <0,0001 | **** | <0,0001 | **** | <0,0001 | | | Nav1.8 | |
| | Nav1.1 | | | ns | 0,9946 | **** | <0,0001 | ns | 0,9455 | ns | 0,8103 | *** | 0,001 | **** | <0,0001 | Nav1.1 | |
| | Nav1.2 | ns | 0,9996 | | | **** | <0,0001 | ns | 0,6173 | ns | 0,386 | **** | <0,0001 | **** | <0,0001 | Nav1.2 | |
| | Nav1.3 | **** | <0,0001 | **** | <0,0001 | | | **** | <0,0001 | **** | <0,0001 | **** | <0,0001 | ns | 0,6188 | Nav1.3 | |
| 1,2 | Nav1.5 | ns | 0,9917 | ns | 0,9205 | **** | <0,0001 | | | ns | 0,9999 | * | 0,0393 | **** | <0,0001 | Nav1.5 | 2 |
| | Nav1.6 | ns | 0,9857 | ns | 0,8933 | **** | <0,0001 | ns | >0,9999 | | | ns | 0,1022 | **** | <0,0001 | Nav1.6 | |
| | Nav1.7 | ns | 0,0998 | * | 0,0317 | **** | <0,0001 | ns | 0,427 | ns | 0,4782 | | | *** | 0,0002 | Nav1.7 | |
| | Nav1.8 | **** | <0,0001 | **** | <0,0001 | ns | >0,9999 | **** | <0,0001 | **** | <0,0001 | **** | <0,0001 | | | Nav1.8 | |
| | Nav1.1 | | | ns | 0,9143 | **** | <0,0001 | ns | >0,9999 | **** | <0,0001 | **** | <0,0001 | **** | <0,0001 | Nav1.1 | |
| | Nav1.2 | ns | 0,9253 | | | **** | <0,0001 | ns | 0,8784 | **** | <0,0001 | **** | <0,0001 | **** | <0,0001 | Nav1.2 | |
| | Nav1.3 | **** | <0,0001 | **** | <0,0001 | | | **** | <0,0001 | **** | <0,0001 | *** | 0,0005 | ns | 0,2288 | Nav1.3 | |
| 4 | Nav1.5 | ns | 0,9561 | ns | 0,353 | **** | <0,0001 | | | **** | <0,0001 | **** | <0,0001 | **** | <0,0001 | Nav1.5 | 6 |
| | Nav1.6 | ** | 0,0093 | **** | <0,0001 | **** | <0,0001 | ns | 0,1632 | | | ns | 0,0888 | **** | <0,0001 | Nav1.6 | |
| | Nav1.7 | **** | <0,0001 | **** | <0,0001 | **** | <0,0001 | **** | <0,0001 | * | 0,0404 | | | ns | 0,4948 | Nav1.7 | |
| | Nav1.8 | **** | <0,0001 | **** | <0,0001 | ns | 0,1624 | **** | <0,0001 | **** | <0,0001 | * | 0,0181 | | | Nav1.8 | |
| ramp rate (mV/ms) | VGSC isoform | Nav1.1 | | Nav1.2 | | Nav1.3 | | Nav1.5 | | Nav1.6 | | Nav1.7 | | Nav1.8 | | VGSC isoform | ramp rate (mV/ms) |

Data: adjusted P value of Tukey's multiple comparisons test

**Supplemental Table 9: Multiple comparisons of the voltage dependence of maximum inward current from ramp current measurements**

| ramp rate (mV/ms) | VGSC isoform | Nav1.1 | | Nav1.2 | | Nav1.3 | | Nav1.5 | | Nav1.6 | | Nav1.7 | | Nav1.8 | | VGSC isoform | ramp rate (mV/ms) |
|---|---|---|---|---|---|---|---|---|---|---|---|---|---|---|---|---|---|
| | Nav1.1 | | | ns | 0,9997 | ns | 0,9905 | **** | <0,0001 | **** | <0,0001 | *** | 0,0007 | **** | <0,0001 | Nav1.1 | |
| | Nav1.2 | ns | 0,3775 | | | ns | 0,9169 | **** | <0,0001 | **** | <0,0001 | ** | 0,0037 | **** | <0,0001 | Nav1.2 | |
| | Nav1.3 | ns | >0,9999 | ns | 0,3334 | | | **** | <0,0001 | **** | <0,0001 | **** | <0,0001 | **** | <0,0001 | Nav1.3 | |
| 0,1 | Nav1.5 | **** | <0,0001 | **** | <0,0001 | **** | <0,0001 | | | **** | <0,0001 | **** | <0,0001 | **** | <0,0001 | Nav1.5 | 0,2 |
| | Nav1.6 | **** | <0,0001 | ** | 0,0043 | **** | <0,0001 | **** | <0,0001 | | | ns | 0,9341 | **** | <0,0001 | Nav1.6 | |
| | Nav1.7 | **** | <0,0001 | ns | 0,1246 | **** | <0,0001 | **** | <0,0001 | ns | 0,9288 | | | **** | <0,0001 | Nav1.7 | |
| | Nav1.8 | **** | <0,0001 | **** | <0,0001 | **** | <0,0001 | **** | <0,0001 | **** | <0,0001 | **** | <0,0001 | | | Nav1.8 | |
| | Nav1.1 | | | ns | 0,7823 | * | 0,0243 | **** | <0,0001 | ns | 0,4405 | ns | 0,9105 | **** | <0,0001 | Nav1.1 | |
| | Nav1.2 | ns | >0,9999 | | | ns | 0,5781 | **** | <0,0001 | * | 0,0124 | ns | 0,1293 | **** | <0,0001 | Nav1.2 | |
| | Nav1.3 | ns | 0,8529 | ns | 0,8762 | | | **** | <0,0001 | **** | <0,0001 | *** | 0,0003 | **** | <0,0001 | Nav1.3 | |
| 0,4 | Nav1.5 | **** | <0,0001 | **** | <0,0001 | **** | <0,0001 | | | **** | <0,0001 | **** | <0,0001 | **** | <0,0001 | Nav1.5 | 0,6 |
| | Nav1.6 | * | 0,0116 | ** | 0,0083 | **** | <0,0001 | **** | <0,0001 | | | ns | 0,9845 | **** | <0,0001 | Nav1.6 | |
| | Nav1.7 | ns | 0,2463 | ns | 0,206 | ** | 0,0055 | **** | <0,0001 | ns | 0,9188 | | | **** | <0,0001 | Nav1.7 | |
| | Nav1.8 | **** | <0,0001 | **** | <0,0001 | **** | <0,0001 | **** | <0,0001 | **** | <0,0001 | **** | <0,0001 | | | Nav1.8 | |
| | Nav1.1 | | | ns | 0,9971 | ns | 0,0528 | **** | <0,0001 | ns | >0,9999 | ns | 0,9861 | **** | <0,0001 | Nav1.1 | |
| | Nav1.2 | ns | 0,9852 | | | ns | 0,2247 | **** | <0,0001 | ns | 0,9981 | ns | 0,8128 | **** | <0,0001 | Nav1.2 | |
| | Nav1.3 | ns | 0,0777 | ns | 0,418 | | | **** | <0,0001 | ns | 0,0598 | ** | 0,0039 | **** | <0,0001 | Nav1.3 | |
| 1,2 | Nav1.5 | **** | <0,0001 | **** | <0,0001 | **** | <0,0001 | | | **** | <0,0001 | **** | <0,0001 | **** | <0,0001 | Nav1.5 | 2 |
| | Nav1.6 | ns | 0,9985 | ns | 0,8414 | * | 0,0162 | **** | <0,0001 | | | ns | 0,9815 | **** | <0,0001 | Nav1.6 | |
| | Nav1.7 | ns | 0,9106 | ns | 0,4456 | ** | 0,0016 | **** | <0,0001 | ns | 0,9958 | | | **** | <0,0001 | Nav1.7 | |
| | Nav1.8 | **** | <0,0001 | **** | <0,0001 | **** | <0,0001 | **** | <0,0001 | **** | <0,0001 | **** | <0,0001 | | | Nav1.8 | |
| | Nav1.1 | | | ns | 0,9997 | ns | 0,2738 | **** | <0,0001 | ns | >0,9999 | ns | 0,868 | **** | <0,0001 | Nav1.1 | |
| | Nav1.2 | ns | 0,9963 | | | ns | 0,5145 | **** | <0,0001 | ns | >0,9999 | ns | 0,6452 | **** | <0,0001 | Nav1.2 | |
| | Nav1.3 | ns | 0,1262 | ns | 0,4242 | | | **** | <0,0001 | ns | 0,3626 | ** | 0,0085 | **** | <0,0001 | Nav1.3 | |
| 4 | Nav1.5 | **** | <0,0001 | **** | <0,0001 | **** | <0,0001 | | | **** | <0,0001 | **** | <0,0001 | **** | <0,0001 | Nav1.5 | 6 |
| | Nav1.6 | ns | >0,9999 | ns | 0,9924 | ns | 0,1014 | **** | <0,0001 | | | ns | 0,79 | **** | <0,0001 | Nav1.6 | |
| | Nav1.7 | ns | 0,9828 | ns | 0,7818 | * | 0,0114 | **** | <0,0001 | ns | 0,9906 | | | **** | <0,0001 | Nav1.7 | |
| | Nav1.8 | **** | <0,0001 | **** | <0,0001 | **** | <0,0001 | **** | <0,0001 | **** | <0,0001 | **** | <0,0001 | | | Nav1.8 | |
| ramp rate (mV/ms) | VGSC isoform | Nav1.1 | | Nav1.2 | | Nav1.3 | | Nav1.5 | | Nav1.6 | | Nav1.7 | | Nav1.8 | | VGSC isoform | ramp rate (mV/ms) |

Data: adjusted P value of Tukey's multiple comparisons test

**Supplemental Table 10: Multiple comparisons of normalized AUC before maximum inward current values from ramp current measurements**

| ramp rate (mV/ms) | VGSC isoform | Nav1.1 | | Nav1.2 | | Nav1.3 | | Nav1.5 | | Nav1.6 | | Nav1.7 | | Nav1.8 | | VGSC isoform | ramp rate (mV/ms) |
|---|---|---|---|---|---|---|---|---|---|---|---|---|---|---|---|---|---|
| | Nav1.1 | | | ns | 0,9994 | ns | >0,9999 | ns | >0,9999 | ns | >0,9999 | ns | >0,9999 | **** | <0,0001 | Nav1.1 | |
| | Nav1.2 | ns | >0,9999 | | | ns | 0,9966 | ns | >0,9999 | ns | 0,9998 | ns | >0,9999 | **** | <0,0001 | Nav1.2 | |
| | Nav1.3 | ns | >0,9999 | ns | >0,9999 | | | ns | >0,9999 | ns | >0,9999 | ns | 0,9998 | **** | <0,0001 | Nav1.3 | |
| 0,1 | Nav1.5 | ns | >0,9999 | ns | >0,9999 | ns | 0,9998 | | | ns | >0,9999 | ns | >0,9999 | **** | <0,0001 | Nav1.5 | 0,2 |
| | Nav1.6 | ns | 0,9985 | ns | 0,9941 | ns | 0,9837 | ns | 0,9999 | | | ns | >0,9999 | **** | <0,0001 | Nav1.6 | |
| | Nav1.7 | ns | >0,9999 | ns | >0,9999 | ns | >0,9999 | ns | >0,9999 | ns | 0,9992 | | | **** | <0,0001 | Nav1.7 | |
| | Nav1.8 | **** | <0,0001 | **** | <0,0001 | **** | <0,0001 | *** | 0,0002 | **** | <0,0001 | **** | <0,0001 | | | Nav1.8 | |
| | Nav1.1 | | | ns | >0,9999 | ns | 0,4036 | ns | 0,9996 | ns | 0,9993 | ns | 0,9985 | **** | <0,0001 | Nav1.1 | |
| | Nav1.2 | ns | >0,9999 | | | ns | 0,3142 | ns | 0,9974 | ns | 0,9954 | ns | 0,9926 | **** | <0,0001 | Nav1.2 | |
| | Nav1.3 | ns | 0,9446 | ns | 0,884 | | | ns | 0,8098 | ns | 0,7267 | ns | 0,7915 | **** | <0,0001 | Nav1.3 | |
| 0,4 | Nav1.5 | ns | >0,9999 | ns | >0,9999 | ns | 0,9754 | | | ns | >0,9999 | ns | >0,9999 | **** | <0,0001 | Nav1.5 | 0,6 |
| | Nav1.6 | ns | 0,9997 | ns | 0,997 | ns | 0,9957 | ns | >0,9999 | | | ns | >0,9999 | **** | <0,0001 | Nav1.6 | |
| | Nav1.7 | ns | >0,9999 | ns | 0,9992 | ns | 0,9915 | ns | >0,9999 | ns | >0,9999 | | | **** | <0,0001 | Nav1.7 | |
| | Nav1.8 | **** | <0,0001 | **** | <0,0001 | **** | <0,0001 | **** | <0,0001 | **** | <0,0001 | **** | <0,0001 | | | Nav1.8 | |
| | Nav1.1 | | | ns | >0,9999 | *** | 0,0001 | ns | 0,1526 | ns | 0,6553 | ns | 0,2951 | **** | <0,0001 | Nav1.1 | |
| | Nav1.2 | ns | >0,9999 | | | *** | 0,0002 | ns | 0,1648 | ns | 0,6709 | ns | 0,3138 | **** | <0,0001 | Nav1.2 | |
| | Nav1.3 | * | 0,0129 | ** | 0,0085 | | | ns | 0,7523 | ns | 0,075 | ns | 0,3326 | * | 0,0455 | Nav1.3 | |
| 1,2 | Nav1.5 | ns | 0,6931 | ns | 0,5962 | ns | 0,7871 | | | ns | 0,9475 | ns | 0,9992 | *** | 0,0007 | Nav1.5 | 2 |
| | Nav1.6 | ns | 0,8749 | ns | 0,7963 | ns | 0,4089 | ns | 0,9996 | | | ns | 0,997 | **** | <0,0001 | Nav1.6 | |
| | Nav1.7 | ns | 0,7572 | ns | 0,6585 | ns | 0,5642 | ns | >0,9999 | ns | >0,9999 | | | **** | <0,0001 | Nav1.7 | |
| | Nav1.8 | **** | <0,0001 | **** | <0,0001 | ** | 0,0018 | **** | <0,0001 | **** | <0,0001 | **** | <0,0001 | | | Nav1.8 | |
| | Nav1.1 | | | ns | >0,9999 | **** | <0,0001 | **** | <0,0001 | *** | 0,0003 | *** | 0,0006 | **** | <0,0001 | Nav1.1 | |
| | Nav1.2 | ns | >0,9999 | | | **** | <0,0001 | **** | <0,0001 | *** | 0,0002 | *** | 0,0003 | **** | <0,0001 | Nav1.2 | |
| | Nav1.3 | **** | <0,0001 | **** | <0,0001 | | | ns | 0,9985 | ns | 0,8316 | ns | 0,8333 | ** | 0,0045 | Nav1.3 | |
| 4 | Nav1.5 | **** | <0,0001 | **** | <0,0001 | ns | 0,9962 | | | ns | 0,6002 | ns | 0,6058 | ns | 0,0828 | Nav1.5 | 6 |
| | Nav1.6 | ns | 0,0508 | ns | 0,054 | ns | 0,0633 | ns | 0,449 | | | ns | >0,9999 | **** | <0,0001 | Nav1.6 | |
| | Nav1.7 | ** | 0,0067 | ** | 0,0075 | ns | 0,3597 | ns | 0,8629 | ns | 0,9934 | | | **** | <0,0001 | Nav1.7 | |
| | Nav1.8 | **** | <0,0001 | **** | <0,0001 | ns | 0,0607 | * | 0,0263 | **** | <0,0001 | **** | <0,0001 | | | Nav1.8 | |
| ramp rate (mV/ms) | VGSC isoform | Nav1.1 | | Nav1.2 | | Nav1.3 | | Nav1.5 | | Nav1.6 | | Nav1.7 | | Nav1.8 | | VGSC isoform | ramp rate (mV/ms) |

Data: adjusted P value of Tukey's multiple comparisons test

**Supplemental Table 11: Correlation of window current values and AUC values from ramp current measurements**

Data: Spearman r with 95% confidence interval, p-values, n = sample size

| | | Nav1.1 | Nav1.2 | Nav1.3 | Nav1.5 | Nav1.6 | Nav1.7 | Nav1.8 |
|---|---|---|---|---|---|---|---|---|
| 0.1 | r | 0.4583 [0.0547; 0.7331] | 0.6902 [0.3443; 0.8711] | 0.2125 [-0.2312; 0.5830] | 0.2857 [-0.3312; 0.7315] | -0.1413 [-0.5324; 0.2995] | 0.3504 [-0.1229; 0.6938] | 0.4161 [-0.0199; 0.7192] |
| | n | 24 | 20 | 23 | 13 | 23 | 20 | 22 |
| | p | 0.0243 (*) | 0.0008 (***) | 0.3304 (ns) | 0.3436 (ns) | 0.5201 (ns) | 0.1299 (ns) | 0.0541 (ns) |
| 0.2 | r | 0.7243 [0.4433; 0.8757] | 0.4506 [0.0099; 0.7448] | 0.3419 [-0.0947; 0.6682] | 0.9000 [0.7112; 0.9677] | -0.0762 [-0.4925; 0.3684] | 0.4887 [0.0449; 0.7714] | 0.5339 [0.1540; 0.7763] |
| | n | 24 | 21 | 23 | 15 | 22 | 20 | 24 |
| | p | < 0.0001 (****) | 0.0403 (*) | 0.1103 (ns) | < 0.0001 (****) | 0.7360 (ns) | 0.0288 (*) | 0.0072 (**) |
| 0.4 | r | 0.8037 [0.5840; 0.9137] | 0.7595 [0.4869; 0.8972] | 0.4991 [0.1074; 0.7567] | 0.6214 [0.1438; 0.8642] | 0.5474 (0.1620; 0.7879) | 0.6030 [0.2055; 0.8297] | 0.5782 [0.2160; 0.8006] |
| | n | 24 | 22 | 24 | 15 | 23 | 20 | 24 |
| | p | < 0.0001 (****) | < 0.0001 (****) | 0.0130 (*) | 0.0155 (*) | 0.0069 (**) | 0.0049 (*) | 0.0031 (**) |
| 0.6 | r | 0.7965 [0.5708; 0.9103] | 0.4828 [0.0636; 0.7572] | 0.3546 [-0.0594; 0.3521] | 0.6441 [0.2027; 0.8680] | 0.3123 [-0.1275; 0.6494] | 0.7260 [0.4174; 0.8844] | 0.6574 [0.3345; 0.8422] |
| | n | 24 | 22 | 25 | 16 | 23 | 21 | 24 |
| | p | < 0.0001 (****) | 0.0229 (*) | 0.0820 (ns) | 0.0085 (**) | 0.1469 (ns) | 0.0002 (***) | 0.0005 (***) |
| 1.2 | r | 0.7009 [0.4042; 0.8641] | 0.8611 [0.6830; 0.9426] | -0.0623 [-0.4563; 0.3521] | 0.6000 [0.1327; 0.8491] | 0.3260 (-0.1365; 0.6718] | 0.7000 [0.3728; 0.8724] | 0.6962 [0.4050; 0.8591] |
| | n | 24 | 22 | 25 | 16 | 21 | 21 | 25 |
| | p | < 0.0001 (****) | < 0.0001 (****) | 0.7673 (ns) | 0.0159 (*) | 0.1493 (ns) | 0.0004 (***) | < 0.0001 (****) |
| 2 | r | 0.7757 [0.5328; 0.9005] | 0.8543 [0.6690; 0.9396] | -0.0538 [-0.4489; 0.3602] | 0.4059 [-0.1283; 0.7575] | 0.3547 [-0.0802; 0.6762] | 0.4338 [-0.0111; 0.7353] | 0.7938 [0.5727; 0.9073] |
| | n | 24 | 22 | 25 | 16 | 23 | 21 | 25 |
| | p | < 0.0001 (****) | < 0.0001 (****) | 0.8011 (ns) | 0.1201 (ns) | 0.0967 (ns) | 0.0485 (*) | < 0.0001 (****) |
| 4 | r | 0.6652 [0.3468; 0.8462] | 0.8046 [0.5708; 0.9177] | -0.0554 [-0.4508; 0.3582] | 0.5147 [0.0094; 0.8106] | 0.3405 [-0.1079; 0.6738] | 0.5564 [0.1373; 0.8065] | 0.5900 [0.2425; 0.8033] |
| | n | 24 | 22 | 25 | 16 | 22 | 20 | 25 |
| | p | 0.0004 (***) | < 0.0001 (****) | 0.7926 (ns) | 0.0436 (*) | 0.1210 (ns) | 0.0108 (*) | 0.0019 (**) |
| 6 | r | 0.6235 [0.2824; 0.8246] | 0.7357 [0.4448; 0.8862] | -0.0569 [-0.4520; 0.3568] | 0.4093 [-0.1041; 0.7505] | 0.3416 [-0.1066; 0.6744] | 0.3744 [-0.0956; 0.7079] | 0.5338 [0.1638; 0.7722] |
| | n | 24 | 22 | 25 | 17 | 22 | 20 | 25 |
| | p | 0.0011 (**) | < 0.0001 (****) | 0.7870 (ns) | 0.1041 (ns) | 0.1197 (ns) | 0.1038 (ns) | 0.0060 (**) |

**Supplemental Table 12: Descriptive statistics of parameters gathered from AP clamping measurements**

Data is mean ± SD, n = sample size

| | Nav isoform | Imax(AP)/ Imax(act) | n | timepoint of Imax(AP)/Ima x(act) (ms) | n | AUC (total) | n | Imax(AP, subthr)/ Imax(act) | n | AUC (subthr) | n | subthr slope (1/ms) | n |
|---|---|---|---|---|---|---|---|---|---|---|---|---|---|
| AP1 | **Nav1.1** | 0,097 ± 0,065 | 27 | 149.6 ± 0.19 | 25 | 0,0013 ± 0,0009 | 26 | 0,0085 ± 0,0064 | 27 | -0,001 ± 0,0025 | 26 | -0,00024 ± 0,00022 | 26 |
| | **Nav1.2** | 0,101 ± 0,052 | 27 | 149.6 ± 0.16 | 23 | 0,0012 ± 0,0009 | 23 | 0,0071 ± 0,0059 | 26 | 0 ± 0,0033 | 25 | -0,00015 ± 0,00013 | 24 |
| | **Nav1.3** | 0,216 ± 0,096 | 27 | 149.5 ± 0.13 | 27 | 0,0048 ± 0,0022 | 27 | 0,0123 ± 0,006 | 26 | 0,0021 ± 0,0038 | 27 | -0,00022 ± 0,00013 | 26 |
| | **Nav1.5** | 0,028 ± 0,016 | 23 | 150.4 ± 0.91 | 20 | 0,0019 ± 0,002 | 24 | 0,0107 ± 0,0094 | 25 | 0,0005 ± 0,006 | 24 | -0,00033 ± 0,00028 | 24 |
| | **Nav1.6** | 0,07 ± 0,046 | 27 | 149.8 ± 0.63 | 23 | 0,0021 ± 0,0031 | 27 | 0,0145 ± 0,0072 | 26 | -0,0008 ± 0,0046 | 26 | -0,00047 ± 0,00032 | 26 |
| | **Nav1.7** | 0,098 ± 0,068 | 26 | 149.8 ± 0.55 | 24 | 0,0029 ± 0,0027 | 25 | 0,0124 ± 0,0089 | 26 | 0,0017 ± 0,004 | 25 | -0,00035 ± 0,00027 | 26 |
| | **Nav1.8** | 0,505 ± 0,14 | 27 | 151.5 ± 0.46 | 27 | 0,0088 ± 0,0034 | 26 | 0,0062 ± 0,0056 | 27 | -0,0021 ± 0,0033 | 25 | -0,00018 ± 0,00015 | 25 |
| AP2 | **Nav1.1** | 0,232 ± 0,1 | 27 | 64.12 ± 0.1 | 27 | 0,0022 ± 0,0012 | 23 | 0,0087 ± 0,0071 | 27 | 0,002 ± 0,0042 | 26 | -0,00151 ± 0,00094 | 25 |
| | **Nav1.2** | 0,216 ± 0,1 | 27 | 64.15 ± 0.14 | 27 | 0,0022 ± 0,002 | 26 | 0,0075 ± 0,0065 | 26 | 0,0014 ± 0,0042 | 26 | -0,00163 ± 0,00149 | 26 |
| | **Nav1.3** | 0,329 ± 0,096 | 27 | 64.14 ± 0.1 | 27 | 0,0041 ± 0,0016 | 27 | 0,0114 ± 0,0069 | 25 | 0,0038 ± 0,0056 | 26 | -0,00206 ± 0,00129 | 26 |
| | **Nav1.5** | 0,033 ± 0,021 | 22 | 64.47 ± 0.28 | 23 | 0,0014 ± 0,0016 | 23 | 0,0066 ± 0,009 | 24 | 0,0009 ± 0,0081 | 24 | -0,00179 ± 0,00151 | 24 |
| | **Nav1.6** | 0,141 ± 0,084 | 27 | 64.14 ± 0.1 | 27 | 0,0028 ± 0,0031 | 26 | 0,0129 ± 0,01 | 26 | 0,003 ± 0,0076 | 26 | -0,00264 ± 0,00151 | 27 |
| | **Nav1.7** | 0,207 ± 0,121 | 26 | 64.12 ± 0.14 | 26 | 0,0044 ± 0,0035 | 25 | 0,021 ± 0,0157 | 26 | 0,008 ± 0,0115 | 26 | -0,00368 ± 0,00259 | 26 |
| | **Nav1.8** | 0,71 ± 0,179 | 27 | 65.49 ± 0.27 | 27 | 0,0104 ± 0,0042 | 27 | 0,0006 ± 0,0057 | 26 | -0,0031 ± 0,0059 | 25 | -0,00123 ± 0,00088 | 26 |
| AP3 | **Nav1.1** | 0,056 ± 0,023 | 25 | 63.13 ± 3.92 | 27 | 0,0058 ± 0,0035 | 25 | 0,0506 ± 0,0216 | 27 | 0,0232 ± 0,0108 | 27 | -0,00516 ± 0,00221 | 27 |
| | **Nav1.2** | 0,072 ± 0,053 | 27 | 64.68 ± 3.56 | 27 | 0,0035 ± 0,0024 | 23 | 0,0441 ± 0,0259 | 27 | 0,0175 ± 0,0123 | 27 | -0,0043 ± 0,00238 | 27 |
| | **Nav1.3** | 0,196 ± 0,051 | 27 | 63.57 ± 3.13 | 27 | 0,0112 ± 0,0045 | 26 | 0,1752 ± 0,0617 | 27 | 0,0767 ± 0,0372 | 27 | -0,01473 ± 0,00481 | 27 |
| | **Nav1.5** | 0,042 ± 0,029 | 21 | 68.23 ± 0.25 | 21 | 0,0043 ± 0,0039 | 22 | 0,0107 ± 0,0089 | 24 | 0,0015 ± 0,0064 | 24 | -0,00164 ± 0,00143 | 24 |
| | **Nav1.6** | 0,061 ± 0,029 | 24 | 67.71 ± 4.59 | 26 | 0,005 ± 0,0036 | 23 | 0,0425 ± 0,0199 | 26 | 0,0186 ± 0,0096 | 26 | -0,00423 ± 0,00196 | 25 |
| | **Nav1.7** | 0,101 ± 0,058 | 26 | 64.05 ± 4.62 | 26 | 0,0081 ± 0,0073 | 26 | 0,0631 ± 0,0303 | 26 | 0,0305 ± 0,0144 | 26 | -0,00677 ± 0,00321 | 26 |
| | **Nav1.8** | 0,317 ± 0,075 | 24 | 71.72 ± 1.9 | 27 | 0,0157 ± 0,0057 | 26 | 0,0162 ± 0,0102 | 25 | -0,0004 ± 0,0071 | 26 | -0,00154 ± 0,00104 | 24 |

**Supplemental Table 13: Multiple comparisons of parameters calculated from AP clamping measurements**

| | VGSC isoform | Nav1.1 | | Nav1.2 | | Nav1.3 | | Nav1.5 | | Nav1.6 | | Nav1.7 | | Nav1.8 | | VGSC isoform | |
|---|---|---|---|---|---|---|---|---|---|---|---|---|---|---|---|---|---|
| Imax(AP)/Imax(act) AP1 | Nav1.1 | | | ns | >0,9999 | ns | 0,6371 | **** | <0,0001 | ns | 0,4575 | ns | >0,9999 | **** | <0,0001 | Nav1.1 | Imax(AP)/Imax(act) AP2 |
| | Nav1.2 | ns | >0,9999 | | | ns | 0,234 | **** | <0,0001 | ns | >0,9999 | ns | >0,9999 | **** | <0,0001 | Nav1.2 | |
| | Nav1.3 | ** | 0,0078 | * | 0,0336 | | | **** | <0,0001 | *** | 0,0002 | ns | 0,1079 | * | 0,0482 | Nav1.3 | |
| | Nav1.5 | ** | 0,006 | ** | 0,0013 | **** | <0,0001 | | | ns | 0,0504 | *** | 0,0001 | **** | <0,0001 | Nav1.5 | |
| | Nav1.6 | ns | >0,9999 | ns | >0,9999 | **** | <0,0001 | ns | 0,272 | | | ns | >0,9999 | **** | <0,0001 | Nav1.6 | |
| | Nav1.7 | ns | >0,9999 | ns | >0,9999 | ** | 0,0064 | ** | 0,0093 | ns | >0,9999 | | | **** | <0,0001 | Nav1.7 | |
| | Nav1.8 | **** | <0,0001 | **** | <0,0001 | ns | 0,2026 | **** | <0,0001 | **** | <0,0001 | **** | <0,0001 | | | Nav1.8 | |
| Imax(AP)/Imax(act) AP3 | Nav1.1 | | | ns | >0,9999 | ns | >0,9999 | ** | 0,002 | ns | 0,7752 | ns | >0,9999 | **** | <0,0001 | Nav1.1 | timepoint of Imax(AP)/Imax(act) AP1 |
| | Nav1.2 | ns | >0,9999 | | | ns | >0,9999 | ** | 0,004 | ns | >0,9999 | ns | >0,9999 | **** | <0,0001 | Nav1.2 | |
| | Nav1.3 | **** | <0,0001 | **** | <0,0001 | | | **** | <0,0001 | * | 0,0125 | ns | 0,7184 | **** | <0,0001 | Nav1.3 | |
| | Nav1.5 | ns | >0,9999 | ns | >0,9999 | **** | <0,0001 | | | ns | >0,9999 | * | 0,0364 | ns | 0,2729 | Nav1.5 | |
| | Nav1.6 | ns | >0,9999 | ns | >0,9999 | **** | <0,0001 | ns | >0,9999 | | | ns | >0,9999 | **** | <0,0001 | Nav1.6 | |
| | Nav1.7 | ns | 0,3866 | ns | >0,9999 | * | 0,0254 | * | 0,013 | ns | >0,9999 | | | **** | <0,0001 | Nav1.7 | |
| | Nav1.8 | **** | <0,0001 | **** | <0,0001 | ns | >0,9999 | **** | <0,0001 | **** | <0,0001 | **** | <0,0001 | | | Nav1.8 | |
| timepoint of Imax(AP)/Imax(act) AP2 | Nav1.1 | | | ns | >0,9999 | ns | >0,9999 | *** | 0,0003 | * | 0,047 | ns | >0,9999 | **** | <0,0001 | Nav1.1 | timepoint of Imax(AP)/Imax(act) AP3 |
| | Nav1.2 | ns | >0,9999 | | | ns | >0,9999 | * | 0,011 | ns | 0,7437 | ns | >0,9999 | **** | <0,0001 | Nav1.2 | |
| | Nav1.3 | ns | >0,9999 | ns | >0,9999 | | | **** | <0,0001 | * | 0,0218 | ns | >0,9999 | **** | <0,0001 | Nav1.3 | |
| | Nav1.5 | *** | 0,0007 | ** | 0,0089 | ** | 0,0077 | | | ns | >0,9999 | ** | 0,006 | ns | 0,2 | Nav1.5 | |
| | Nav1.6 | ns | >0,9999 | ns | >0,9999 | ns | >0,9999 | ** | 0,0071 | | | ns | 0,4707 | *** | 0,0003 | Nav1.6 | |
| | Nav1.7 | ns | >0,9999 | ns | >0,9999 | ns | >0,9999 | ** | 0,0013 | ns | >0,9999 | | | **** | <0,0001 | Nav1.7 | |
| | Nav1.8 | **** | <0,0001 | **** | <0,0001 | **** | <0,0001 | ns | 0,0794 | **** | <0,0001 | **** | <0,0001 | | | Nav1.8 | |
| AUC (total) AP1 | Nav1.1 | | | ns | >0,9999 | * | 0,0312 | ns | >0,9999 | ns | >0,9999 | ns | 0,2419 | **** | <0,0001 | Nav1.1 | AUC (total) AP2 |
| | Nav1.2 | ns | >0,9999 | | | * | 0,0233 | ns | >0,9999 | ns | >0,9999 | ns | 0,2038 | **** | <0,0001 | Nav1.2 | |
| | Nav1.3 | **** | <0,0001 | **** | <0,0001 | | | **** | <0,0001 | ns | 0,5491 | ns | >0,9999 | * | 0,0111 | Nav1.3 | |
| | Nav1.5 | ns | >0,9999 | ns | >0,9999 | ** | 0,0019 | | | ns | 0,3289 | ** | 0,0019 | **** | <0,0001 | Nav1.5 | |
| | Nav1.6 | ns | >0,9999 | ns | >0,9999 | ** | 0,0024 | ns | >0,9999 | | | ns | >0,9999 | **** | <0,0001 | Nav1.6 | |
| | Nav1.7 | ns | 0,4951 | ns | 0,1524 | ns | 0,3604 | ns | >0,9999 | ns | >0,9999 | | | ** | 0,0012 | Nav1.7 | |
| | Nav1.8 | **** | <0,0001 | **** | <0,0001 | ns | 0,8336 | **** | <0,0001 | **** | <0,0001 | *** | 0,0003 | | | Nav1.8 | |
| AUC (total) AP3 | Nav1.1 | | | ns | >0,9999 | ns | 0,3892 | ns | >0,9999 | * | 0,0251 | ns | >0,9999 | ns | >0,9999 | Nav1.1 | Imax(AP, subthr)/ Imax(act) AP1 |
| | Nav1.2 | ns | 0,6046 | | | * | 0,026 | ns | >0,9999 | *** | 0,0008 | ns | 0,1788 | ns | >0,9999 | Nav1.2 | |
| | Nav1.3 | * | 0,0139 | **** | <0,0001 | | | ns | >0,9999 | ns | >0,9999 | ns | >0,9999 | * | 0,0102 | Nav1.3 | |
| | Nav1.5 | ns | >0,9999 | ns | >0,9999 | **** | <0,0001 | | | ns | 0,6058 | ns | >0,9999 | ns | 0,7048 | Nav1.5 | |
| | Nav1.6 | ns | >0,9999 | ns | >0,9999 | *** | 0,0008 | ns | >0,9999 | | | ns | >0,9999 | *** | 0,0003 | Nav1.6 | |
| | Nav1.7 | ns | >0,9999 | ns | 0,0779 | ns | 0,1366 | ns | 0,6611 | ns | >0,9999 | | | ns | 0,0823 | Nav1.7 | |
| | Nav1.8 | **** | <0,0001 | **** | <0,0001 | ns | >0,9999 | **** | <0,0001 | **** | <0,0001 | *** | 0,0004 | | | Nav1.8 | |
| Imax(AP, subthr)/ Imax(act) AP2 | Nav1.1 | | | ns | >0,9999 | *** | 0,0002 | **** | <0,0001 | ns | >0,9999 | ns | >0,9999 | *** | 0,0001 | Nav1.1 | Imax(AP, subthr)/ Imax(act) AP3 |
| | Nav1.2 | ns | >0,9999 | | | **** | <0,0001 | *** | 0,0002 | ns | >0,9999 | ns | >0,9999 | ** | 0,0075 | Nav1.2 | |
| | Nav1.3 | ns | >0,9999 | ns | >0,9999 | | | **** | <0,0001 | **** | <0,0001 | ** | 0,005 | **** | <0,0001 | Nav1.3 | |
| | Nav1.5 | ns | >0,9999 | ns | >0,9999 | ns | 0,632 | | | *** | 0,0002 | **** | <0,0001 | ns | >0,9999 | Nav1.5 | |
| | Nav1.6 | ns | >0,9999 | ns | 0,6664 | ns | >0,9999 | ns | 0,322 | | | ns | >0,9999 | ** | 0,0071 | Nav1.6 | |
| | Nav1.7 | ns | 0,0566 | ** | 0,0019 | ns | 0,985 | *** | 0,0007 | ns | >0,9999 | | | **** | <0,0001 | Nav1.7 | |
| | Nav1.8 | *** | 0,0005 | * | 0,0259 | **** | <0,0001 | ns | 0,0931 | **** | <0,0001 | **** | <0,0001 | | | Nav1.8 | |
| AUC (subthr) AP1 | Nav1.1 | | | ns | >0,9999 | ns | >0,9999 | ns | >0,9999 | ns | >0,9999 | ns | 0,1846 | * | 0,0148 | Nav1.1 | AUC (subthr) AP2 |
| | Nav1.2 | ns | >0,9999 | | | ns | >0,9999 | ns | >0,9999 | ns | >0,9999 | * | 0,0206 | ns | 0,1379 | Nav1.2 | |
| | Nav1.3 | ns | 0,087 | ns | 0,982 | | | ns | >0,9999 | ns | >0,9999 | ns | >0,9999 | *** | 0,0004 | Nav1.3 | |
| | Nav1.5 | ns | >0,9999 | ns | >0,9999 | ns | >0,9999 | | | ns | >0,9999 | * | 0,0311 | ns | 0,1393 | Nav1.5 | |
| | Nav1.6 | ns | >0,9999 | ns | >0,9999 | ns | 0,4136 | ns | >0,9999 | | | ns | 0,4746 | ** | 0,0041 | Nav1.6 | |
| | Nav1.7 | ns | 0,1794 | ns | >0,9999 | ns | >0,9999 | ns | >0,9999 | ns | 0,7415 | | | **** | <0,0001 | Nav1.7 | |
| | Nav1.8 | ns | >0,9999 | ns | >0,9999 | ** | 0,003 | ns | 0,2227 | ns | >0,9999 | ** | 0,008 | | | Nav1.8 | |
| AUC (subthr) AP3 | Nav1.1 | | | ns | >0,9999 | ns | >0,9999 | ns | >0,9999 | ** | 0,0073 | ns | >0,9999 | ns | >0,9999 | Nav1.1 | subthr slope AP1 |
| | Nav1.2 | ns | >0,9999 | | | ns | >0,9999 | ns | 0,1638 | **** | <0,0001 | * | 0,0441 | ns | >0,9999 | Nav1.2 | |
| | Nav1.3 | ** | 0,008 | **** | <0,0001 | | | ns | >0,9999 | ns | 0,07 | ns | >0,9999 | ns | >0,9999 | Nav1.3 | |
| | Nav1.5 | **** | <0,0001 | ** | 0,0038 | **** | <0,0001 | | | ns | 0,8171 | ns | >0,9999 | ns | 0,5191 | Nav1.5 | |
| | Nav1.6 | ns | >0,9999 | ns | >0,9999 | *** | 0,0001 | *** | 0,0009 | | | ns | >0,9999 | *** | 0,0003 | Nav1.6 | |
| | Nav1.7 | ns | >0,9999 | ns | 0,2842 | ns | 0,288 | **** | <0,0001 | ns | 0,848 | | | ns | 0,1652 | Nav1.7 | |
| | Nav1.8 | **** | <0,0001 | *** | 0,0005 | **** | <0,0001 | ns | >0,9999 | **** | <0,0001 | **** | <0,0001 | | | Nav1.8 | |
| subthr slope AP2 | Nav1.1 | | | ns | >0,9999 | *** | 0,0005 | **** | <0,0001 | ns | >0,9999 | ns | >0,9999 | **** | <0,0001 | Nav1.1 | subthr slope AP3 |
| | Nav1.2 | ns | >0,9999 | | | **** | <0,0001 | ** | 0,0087 | ns | >0,9999 | ns | 0,4518 | ** | 0,0069 | Nav1.2 | |
| | Nav1.3 | ns | >0,9999 | ns | >0,9999 | | | **** | <0,0001 | **** | <0,0001 | * | 0,0309 | **** | <0,0001 | Nav1.3 | |
| | Nav1.5 | ns | >0,9999 | ns | >0,9999 | ns | >0,9999 | | | ** | 0,0056 | **** | <0,0001 | ns | >0,9999 | Nav1.5 | |
| | Nav1.6 | ns | 0,129 | * | 0,0333 | ns | >0,9999 | ns | 0,3252 | | | ns | 0,7992 | ** | 0,0045 | Nav1.6 | |
| | Nav1.7 | ** | 0,0086 | ** | 0,0016 | ns | 0,8037 | * | 0,028 | ns | >0,9999 | | | **** | <0,0001 | Nav1.7 | |
| | Nav1.8 | ns | >0,9999 | ns | >0,9999 | ns | 0,2756 | ns | >0,9999 | ** | 0,0036 | *** | 0,0001 | | | Nav1.8 | |
| timepoint of Imax(AP)/Imax(act) AP3 after exclusion of measurements with transient current artifacts | Nav1.1 | | | | | | | | | | | | | | | | |
| | Nav1.2 | ns | >0,9999 | | | | | | | | | | | | | | |
| | Nav1.3 | ns | >0,9999 | ns | >0,9999 | | | | | | | | | | | | |
| | Nav1.5 | **** | <0,0001 | ** | 0,0033 | ** | 0,0012 | | | | | | | | | | |
| | Nav1.6 | *** | 0,0006 | ns | 0,0548 | * | 0,0281 | ns | >0,9999 | | | | | | | | |
| | Nav1.7 | ns | >0,9999 | ns | >0,9999 | ns | >0,9999 | **** | <0,0001 | ** | 0,0013 | | | | | | |
| | Nav1.8 | **** | <0,0001 | **** | <0,0001 | **** | <0,0001 | ns | 0,2665 | ** | 0,0037 | **** | <0,0001 | | | | |
| | VGSC isoform | Nav1.1 | | Nav1.2 | | Nav1.3 | | Nav1.5 | | Nav1.6 | | Nav1.7 | | Nav1.8 | | VGSC isoform | |

Data: adjusted P value of Dunn's multiple comparisons test
